# SpinForecast: chain-free probabilistic backbone assignment of intrinsically disordered proteins from NMR chemical shifts

**DOI:** 10.64898/2026.07.31.740808

**Authors:** James T. Eaton, Jasmine Cornish, Kirsten M. Silvey, Pijush Chakraborty, Thomas Löhr, Gogulan Karunanithy, Gabriella T. Heller

## Abstract

Assigning peaks in NMR spectra to specific residues is an essential but time-consuming step in the study of intrinsically disordered proteins (IDPs). Conventional approaches rely on building chains of sequential connectivities between peaks, which are particularly prone to failure in disordered systems due to spectral overlap, missing peaks, and proline-rich sequences. Here we present SpinForecast, a tool that performs chain-free probabilistic backbone assignment of IDPs from chemical shifts alone, without requiring peaks to be linked into sequential chains. SpinForecast predicts residue-specific chemical shift distributions from a disorder-filtered subset of the Biomolecular Magnetic Resonance Data Bank (BMRB), incorporating nearest-neighbour residue effects and corrections for temperature and pH. Experimental chemical shifts are then assigned to residues by Bayes’ theorem, using residue-specific chemical shift distributions as likelihoods. We validate SpinForecast on three disordered proteins, IAPP (37 residues), NUPR1 (82 residues), and JPT2 (218 residues), achieving 100% confidence assignments for 74%, 57%, and 44% of in-distribution peaks, respectively, all with at least 99% accuracy. Where single assignments cannot be determined for these systems, the correct assignment was contained within the returned set of candidate assignments in greater than 97% of cases. SpinForecast is freely available at https://tools.bindresearch.org/bindbox/SpinForecast.

## Introduction

Intrinsically disordered proteins (IDPs) and intrinsically disordered regions (IDRs), which account for approximately 30% of the human proteome,^1^ play central roles in cellular signalling,^2^ transcriptional regulation,^3^ and disease, yet their structural characterisation remains challenging.^1,4^ Solution-state NMR spectroscopy is uniquely suited to studying these dynamic systems, providing atomic-resolution information on conformational ensembles in solution.^5–13^ However, extracting residue-specific information from NMR spectra requires that peaks be assigned to specific atoms and residues, a process that can take weeks to months even for an experienced spectroscopist.

The backbone assignment of IDPs presents several challenges not encountered in structured systems. The conformational averaging inherent to IDPs compresses chemical shifts towards random-coil values, producing severe spectral overlap that increases ambiguity in both peak picking and sequential assignment.^11,12,14^ IDPs are also enriched in proline residues relative to structured proteins,^15^ which interrupts the amide proton magnetisation transfer pathway and can give rise to additional minor-population peaks through cis-trans proline isomerism.^16^ Furthermore, conventional assignment strategies rely on building chains of sequential connectivities between peaks from neighbouring residues either using through-bond^17^ or through-space^18^ inter-residue correlation experiments. A single missing peak breaks the chain, potentially leaving large stretches of the sequence unassigned.

Backbone assignment typically requires double (^13^C, ^15^N) or triple (^2^H, ^13^C, ^15^N) isotopically labelled samples and a suite of multidimensional NMR experiments.^19^ The resulting spectra are analysed using software packages such as CCPN AnalysisAssign,^20^ CARA,^21^ MARS,^22^ NMRFAM-SPARKY,^23^ and POKY,^24^ which guide the user through the process of building sequential connectivity networks. More recently, fully automated approaches such as ARTINA^25,26^ have been developed that integrate peak picking and assignment, though these require careful manual validation, particularly for IDPs where spectral overlap and missing peaks increase assignment ambiguity. Methods utilising higher dimensionality (>3D) NMR experiments^27–34^ and direct detection of ^1^HA,^31,35–37^ ^13^C,^30,32,37–41^ or ^15^N,^42–44^ nuclei have been shown to increase resolution in highly overlapped spectral areas of disordered proteins, improving assignment coverage. Furthermore, the ^1^HA,^13^C and ^15^N direct detection methods enable proline residues to be directly observed, which is not possible for amide proton detected experiments. However, despite these technical advancements, assignment of disordered proteins remains challenging. Chemical shift prediction tools such as SHIFTX2^45^ and SPARTA+^46^ can aid assignment in structured proteins but require a static structure and are therefore not applicable to IDPs. For disordered proteins, random coil chemical shift predictors, such as POTENCI,^47^ provide sequence, temperature and pH corrected chemical shift predictions, but return single-point estimates rather than distributions, limiting their use in probabilistic assignment workflows.

Here we present SpinForecast, a freely available online tool for chain-free probabilistic backbone assignment of IDPs from chemical shifts, which exploits the largely untapped chemical shift data held in the Biomolecular Magnetic Resonance Data Bank (BMRB),^48^ the primary repository for assigned protein NMR chemical shifts. SpinForecast filters these data by residue type and nearest-neighbour residue type to build residue-specific chemical shift distributions. To ensure that only genuinely disordered chemical shifts are retained, disorder is classified using AlphaFold2^49^ predicted local distance difference test (pLDDT) scores, retaining only residues within stretches of at least 30 consecutive residues with pLDDT below 70, a conservative threshold given that AlphaFold2 is known to over-fold disordered regions. Temperature and pH corrections are then applied to these distributions following the approach of POTENCI.^47^ Experimental chemical shifts are assigned to residues by applying Bayes’ theorem, using the residue-specific chemical shift distributions as likelihoods. Crucially, each peak is treated independently, making SpinForecast robust to missing peaks and applicable to minor population states such as those arising from cis-trans proline isomerism. We validate SpinForecast on three IDPs of increasing size, IAPP (37 residues), NUPR1 (82 residues), and JPT2 (218 residues), and assess its performance on a mixed ordered/disordered system, CBP-TAZ4 (306 residues).

## Materials and Methods

### Disorder classification of BMRB chemical shifts

A pipeline was developed to extract chemical shift assignments from the BMRB^48^ corresponding to residues within predicted IDRs. The pipeline integrates BMRB sequence data, AlphaFold2-predicted per-residue confidence scores, and sequence-based mapping between BMRB entries and UniProt entries. The BMRB was accessed in August 2025, yielding 17,054 entries. For each entry, protein sequences were extracted from the *polymer_sequences* loop, retaining only entities with polymer type *polypeptide(L);* DNA, RNA, and other non-polypeptide polymers were excluded. Each entity within a multi-entity entry (e.g., protein complexes) was treated as an independent unit in all subsequent analyses, and chemical shift assignments were retrieved on a per-entity basis using the corresponding *Entity_ID*. BMRB entities were then grouped by source organism and mapped to UniProt reference proteomes, retaining only proteomes associated with more than ten BMRB entities.

Predicted protein structures for each retained proteome were obtained from the AlphaFold Protein Structure Database (AFDB) version 4,^50^ generated using AlphaFold2.^49^ Per-residue Predicted Local Distance Difference Test (pLDDT) confidence scores were extracted from the B-factor column of the coordinate files. IDRs were defined as stretches of at least 30 consecutive residues with pLDDT below 70,^4,51–53^ and proteins lacking any region meeting this criterion were excluded from further analysis. Full-length canonical UniProt sequences for all entries containing at least one predicted disordered region were collated and used to build a DIAMOND^54^ sequence database. Each BMRB entity sequence was then queried against this database using *diamond blastp* in default sensitivity mode. For each BMRB entity, the UniProt hit with the highest percent identity was retained and matches with percent identity below 90% were discarded, ensuring near-identical sequence correspondence and accounting for construct-level modifications such as purification tags, truncations, or point mutations.

Residue-level correspondence for each BMRB entity and UniProt pair passing the identity threshold was established using *SequenceMatcher* from Python’s *difflib* module. The aligned residue positions were then cross-referenced against the predicted disordered regions of the UniProt sequence, and every chemical shift entry in the BMRB *Atom_chem_shift* loop corresponding to residues falling within a predicted disordered region was retained. Assignments were counted without deduplication across experimental conditions, samples, or separate BMRB entries mapping to the same UniProt position. The resulting dataset reflects the total volume of NMR chemical shift data deposited for predicted disordered residues, rather than the number of unique disordered atoms assigned at least once. In total, 10,232,379 chemical shifts from 12,883 unique sequences were retrieved from the BMRB, of which 480,599 (from 2,802 unique sequences) corresponded to predicted disordered residues. Removing the disordered chemical shifts from the total set obtained from the BMRB and removing entries from proteins described as having a non-structured physical-state (denatured, disordered, intrinsically disordered, partially disordered, molten globule or unfolded) led to 9,573,332 non-disordered chemical shifts (from 12,439 unique sequences).

### Construction of a referenced disordered chemical shift dataset

To construct residue-specific chemical shift distributions, the disorder-filtered BMRB dataset was first corrected for temperature and pH following the approach of POTENCI.^47,55–57^ These corrections, denoted by the vector *c̄*, convert all chemical shifts, *δ̄*, to a reference state of T=298K and pH=7.0, giving corrected chemical shifts *δ̄*_*corr*_ = *δ̄* + *c̄*. After standardising the temperature, ionic strength and pressure units for each entry to Kelvin (K), molar (M) and atmosphere (atm), respectively, chemical shifts from BMRB entries with experimental conditions outside the following ranges were excluded prior to correction: pH 1-14, 273-330 K, and ionic strength 0-8 M. Entries with missing values for pressure, temperature, pH or ionic strength were also excluded, as were entries measured at pressures other than 1 atmosphere. Entries containing more than one entity, such as those involving a potential binding partner were also removed. Only chemical shifts from H, N, CA, CB, C, HA and HB atoms were retained.

Different research groups may use slightly different referencing standards when depositing chemical shifts, leading to systematic offsets between entries that would artificially broaden the residue-specific chemical shift distributions if uncorrected. To correct for this, a referencing correction was calculated for each atom type (CA, CB, N, C, H, HA, HB) within a given entry. For each backbone atom type, a global mean chemical shift across the entire set of disordered residues was first calculated as a function of residue type, denoted as 〈*δ̄*_*corr*_ (atom, residue)〉, where 〈 〉 denotes the mean over all values. The referencing value for each atom type in each BMRB entry, denoted by ref(id, atom) was then calculated as follows, where N_residues_(id, atom) is the number of different residue types present for that atom type in the entry:

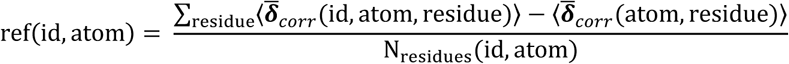

Referencing was only performed on each atom type within a given BMRB entry if more than ten chemical shifts were present for that atom type. Referencing corrections for different atom types within the same entry are not constrained to be equal, since chemical shifts for different atom types may have been obtained from different experiments with independent referencing errors. For example, CA and CO chemical shifts are often measured in separate experiments (such as HNCACB and HNCO respectively) which may carry independent referencing offsets. Following these corrections, the final disordered residue dataset contained 217,357 backbone chemical shifts from 847 unique sequences and 957 BMRB entries. Chemical shifts from C- and N-terminal residues were excluded from temperature, pH, and referencing corrections as their temperature and pH correction factors were not available through POTENCI. As a result, atoms from C- and N-terminal residues have wider predicted distributions.

### Calculating residue-specific disordered chemical shift distributions

To construct residue-specific chemical shift distributions for each atom type and residue within a given sequence, the corrected disordered chemical shift dataset was filtered by residue type (*i*) and nearest-neighbour residue type. For H and N atom types, the dataset was first filtered by the preceding residue type (*i* − 1), as these atoms are particularly sensitive to the identity of the preceding residue through its influence on the peptide bond conformation, and then additionally by the following residue type (*i* + 1) if at least 20 chemical shifts remained after the initial filter. For all other atom types, filtering was performed first by the following residue type (*i* + 1), and then by the preceding residue type (*i* − 1), again subject to the same minimum data threshold of 20 chemical shifts. This minimum threshold ensures that the resulting distributions are based on sufficient experimental data to be statistically meaningful. Temperature and pH corrections were then applied to the resulting distributions for the experimental conditions of the protein under study, following the same approach as POTENCI.^47,55–57^ A Gaussian kernel density estimation (KDE) was applied to each chemical shift distribution using the Scott bandwidth method in SciPy,^58^ producing a smooth probability density function for each atom type and residue position in the sequence.

### Probabilistic peak assignment

To calculate the posterior probabilities for peaks in a protein sequence of length N, a probabilistic assignment procedure was performed. For a given peak *i* in the peaklist, containing observed chemical shift values *δ_i_* for different atom types *j* (e.g. N, CA, CAi-1), the likelihood of observing each chemical shift given that the peak originated from residue *k*, P(*δ*_j,*i*_|*k*), was calculated by evaluating the residue-specific kernel density estimate (KDE) for residue *k* and atom type *j* at the observed chemical shift value *δ*_j,*i*_. Assuming conditional independence between atom types given residue identity, the overall likelihood of peak *i* originating from residue *k* was calculated as:

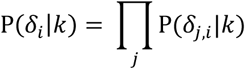

To convert these likelihoods into posterior probabilities, Bayes’ theorem was applied:

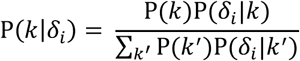

where the denominator represents the marginalisation over all candidate residues *k̄* in the sequence, ensuring that the posterior probabilities sum to one. Because the sequence contains a finite number of residues *N*, this marginalisation is computed directly. A uniform prior probability P(*k*) = 1/N was assigned to each residue, reflecting the absence of any prior information about the likely assignment. Since the prior was identical for all residues it cancels from both numerator and denominator, leaving:

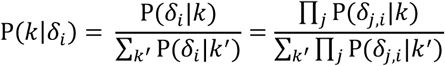

In practice, the posterior probabilities were calculated in log space to avoid numerical instability arising from the multiplication of many small probability values.

Defining:

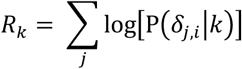

and

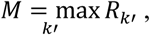

the posterior probability was then computed as:

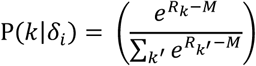

where *M* is a single fixed constant computed across all residues, and *k*′ is a dummy index running over all *N* residues in the denominator sum. Subtracting *M* from each exponent is a standard numerical trick to prevent underflow when exponentiating log probabilities, and does not affect the resulting posterior probabilities. This procedure was repeated for all residues *k*, and residues with posterior probabilities *P*(*k*|*δ_i_*) > 0.01 were retained as candidate assignments. We found that inclusion of the amide proton (^1^H) chemical shifts reduced assignment reliability; therefore, when included, amide proton chemical shifts were weighted by a factor of 0.5 in the assignment procedure by multiplying any P(*δ*_*j*=*H*,*i*_|*k*) value by 0.5. In the optimisation of atom type combinations procedure introduced below, amide proton chemical shifts were omitted entirely. After assignment prediction, peaks in the supplied peaklist were grouped into the following categories: 100% probability, greater than 75% probability, greater than 50% probability, less than 50% probability and out-of-distribution (described below), allowing highly confident assignments to be identified more readily. Posterior probabilities of 100% are rounded up from 99%. Assignments labelled as having 100% posterior probability have only a single candidate residue from the sequence that fits the data for that peak in the supplied peaklist (i.e. all other residues have less than 1% probability); it does not imply that this assignment will be 100% accurate.

To assess the contribution of individual atom types to a predicted assignment, a leave-one-out analysis was performed. For each peak *i* assigned to residue *k* with posterior probability *P*(*k*|*δ_i_*), the change in posterior probability upon removal of atom type a was calculated as:

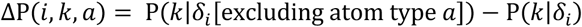

This procedure was repeated for all atom types *a*, producing *ΔP*(*i*, *k*, *a*) values on a scale from −1 to 1. Values close to zero indicate that removal of atom type *a* has little effect on the assignment probability for residue *k*. Positive values (Δ*P*(*i*, *k*, *a*) > 0) indicate that removing atom type *a* increases the assignment probability, implying that atom *a* disfavours the assignment. Negative values (Δ*P*(*i*, *k*, *a*) < 0) indicate that removing atom type *a* decreases the assignment probability, suggesting that atom *a* positively supports the assignment.

### Out-of-distribution classification

Following assignment, each peak is assessed to determine whether its chemical shifts are consistent with the residue-specific chemical shift distributions of the most probable assignment. If a chemical shift falls far outside the expected range for a given residue and atom type, this may indicate an incorrect assignment, a structured region, or a peak that cannot be reliably assigned using the available chemical shift data. Peaks meeting this criterion are classified as out-of-distribution and excluded from the reported results.

For each peak *i*, assigned to residue *k*, the percentile of each observed chemical shift *δ*_j,*i*_ within the residue-specific KDE for residue *k* and atom type *j*, denoted Percentile(δ*j,i*, *k*), was calculated as follows. The KDE for residue *k* and atom type *j* was constructed from a set of experimental chemical shifts {*D_k_*_,j_} with 10^th^ and 90^th^ percentile values denoted *l* and ℎ, respectively. A margin *m* = 0.2(ℎ − *l*) was defined, and extended integration bounds were set to *l*_ext_ = *l* − *m* and ℎ_ext_ = ℎ + *m*, ensuring that the integration region captures the tails of the distribution. These bounds were further extended to ensure inclusion of the observed chemical shift *δ*_*j,i*_, giving the final bounds *l*_final_ = min(*l*_ext_, *δ*_j,*i*_, − *λ*) and ℎ_final_ = max(ℎ_ext_, *δ*_j,*i*_ + λ), where λ is a small numerical tolerance set to 1×10^-6^. A truncated cumulative distribution (CDF) function was then evaluated as:

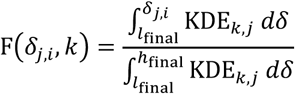

A truncated rather than full CDF was used to ensure numerical stability. The percentile of δ_*j,i*_ within the residue-specific KDE for residue *k* was then evaluated as:

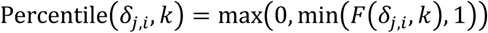

Chemical shifts with percentiles below 0.05 or above 0.95 were classed as outliers. If at least two outliers were identified across all atom types for the most probable assignment of peak *i*, the peak was classified as out-of-distribution and excluded from subsequent analysis.

### Optimisation of atom type combinations

To determine which combination of atom types yields the highest number of accurate assignments, a global optimisation was performed across the proteins used for validation: IAPP (37 residues), NUPR1 (82 residues), JPT2 (218 residues), and CBP-TAZ4 (306 residues). CBP-TAZ4 is a multi-domain protein containing both disordered and structured sections and was included as a stress test to assess the performance of SpinForecast outside of a purely disordered system. Assignments of IAPP, NUPR1, JPT2, and CBP-TAZ4 were not present in the disorder-filtered BMRB dataset used to construct residue-specific chemical shift distributions, ensuring that the validation is independent of the training data. Peaklists for IAPP and CBP-TAZ4 were obtained from previously reported BMRB entries (IAPP: 51259,^59^ CBP-TAZ4: 52655^60^), while peaklists for NUPR1 and JPT2 were obtained from the new assignments described in the following section. The protein sequences and experimental conditions used for each protein are given in **Supplementary Table 1**, and the peaklists are provided in **Supplementary Tables 3-6.** For each peak, the chemical shifts for H, N, Ni-1, Ni+1, CA, CAi-1, CB, CBi-1, C and Ci-1 chemical shifts are reported where available. Amide proton (^1^H) chemical shifts were excluded from assignment calculations in this procedure. Each peak was indexed by its correct assignment to allow comparison between the SpinForecast prediction and the true assignment; however, this index was not used by SpinForecast to influence any assignment predictions.

For each number of atom types from 2 to 9, all possible combinations of atom types were enumerated. For each combination, SpinForecast assignment predictions were performed on all four proteins. Where a peak was missing a chemical shift for a given atom, for example glycine residues without CB chemical shifts, assignment probabilities were calculated using the remaining atom types only. Peaks classified as out-of-distribution by the procedure were excluded prior to further analysis. The following metrics were then recorded for each combination: percentage of peaks assigned with 100% posterior probability (*n*), the accuracy of those 100% probability assignments (*y*), the accuracy of assignments with posterior probability greater than 75%, the accuracy of the set of candidate assignments returned for each peak, and the percentage of peaks classified as in-distribution. The optimal combination of atom types for each number of atom types was defined as that which maximised *n* + *y*, averaged across all four proteins, ensuring that the combination which balanced the largest number of high-confidence assignments and their accuracy was found. A table of results for the optimal combination for each number of atoms is included in **Supplementary Table 2**.

### Backbone assignment of NUPR1 and JPT2

The backbone (H,N,CA,CB,C) assignments of NUPR1 and JPT2 were performed independently and without the use of SpinForecast, providing ground truth assignments against which SpinForecast predictions could be validated. NMR spectra were processed using NMRPipe,^61^ with NUS reconstruction performed using SMILE.^62^ Peaks were picked and assigned using CCPNMR AnalysisAssign.^20^ Expression and purification protocols for NUPR1 and JPT2 are provided in the Supplementary Methods. Full details of sample composition, experimental conditions, NMR experiment types, and acquisition parameters used to assign each protein are provided in **Supplementary Tables 7** and **8**, respectively.

## Results

The BMRB^48^ contains a largely untapped resource of chemical shift data for IDPs and IDRs. Overlays of CA chemical shift distributions of all amino acid types reveal striking separation between residue types for disordered residues that is not observed for structured residues (**Figure 1**). In structured proteins, conformational diversity broadens chemical shift distributions and increases overlap between residue types. For IDPs and IDRs, conformational averaging towards random-coil values produces narrower, better-separated distributions for each amino acid type. We hypothesised that these narrow, well-separated distributions could be exploited to enable probabilistic backbone assignment of IDPs from chemical shifts alone, without requiring sequential connectivity. An interactive dashboard allowing users to explore these distributions and compare disordered and structured chemical shifts is available at https://tools.bindresearch.org/bindbox/BMRB_Chemical_Shifts. Analogous overlays for CB, C and N chemical shifts are provided in **Supplementary Figure 1**.

**Figure 1.**
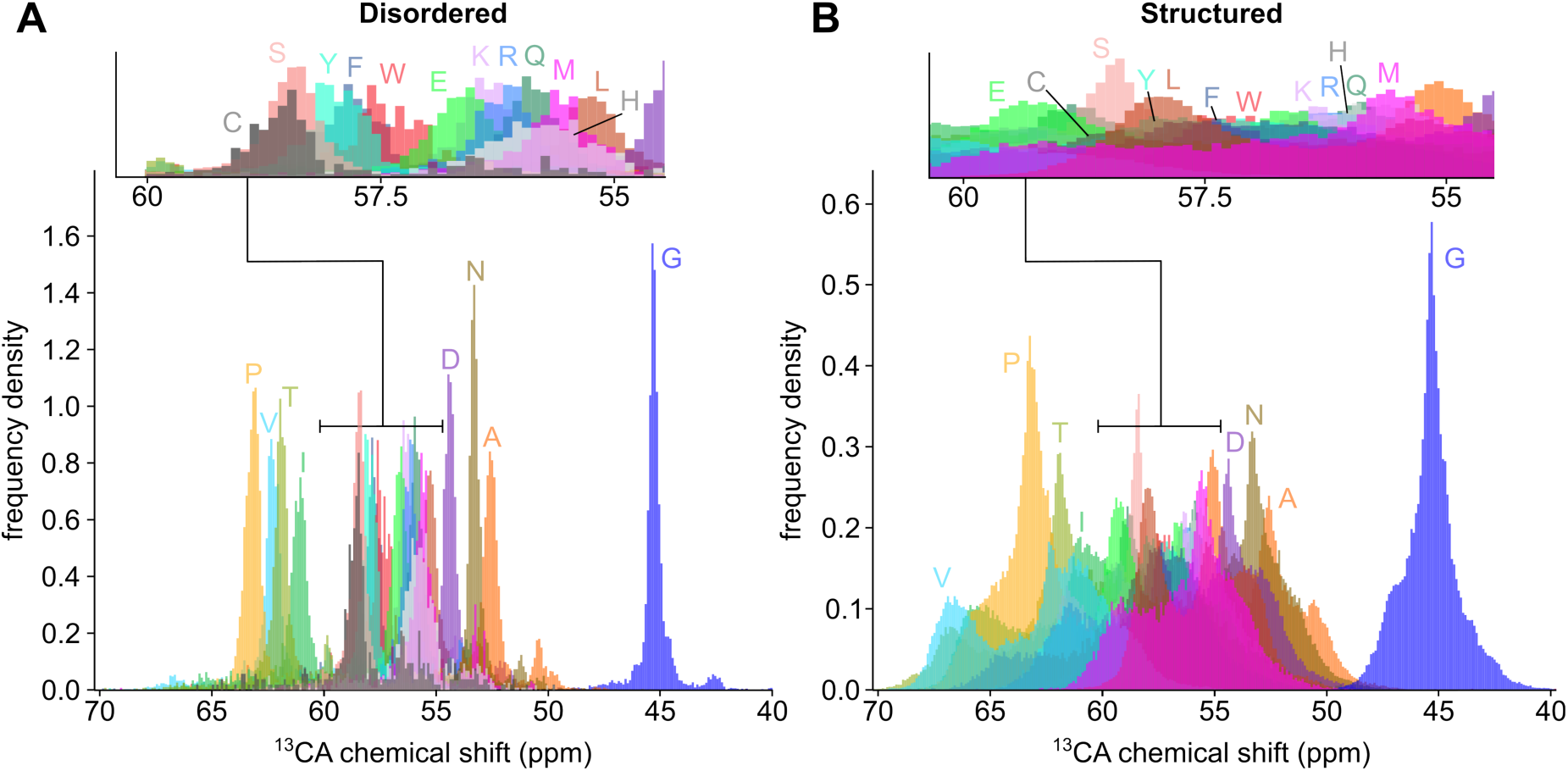
Overlays of CA chemical shift distributions for all amino acid types for disordered residues (**A**) and structured residues (**B**) within the BMRB database. There is substantial separation in CA chemical shift space between residue types for disordered residues that is not present for structured residues. Disordered chemical shifts were extracted using the disorder classification procedure described in the Materials and Methods, and the structured chemical shifts contain all values extracted from the BMRB not classified as disordered. The plots were produced using the interactive dashboard at https://tools.bindresearch.org/bindbox/BMRB_Chemical_Shifts using histogram bin widths of 0.1 ppm. Residue types are annotated by their single letter code. Analogous plots for CB, C and N chemical shifts are included in **Supplementary Figure 1**.

SpinForecast constructs residue-specific chemical shift distributions by filtering a disorder-classified subset of the BMRB by residue type and nearest-neighbour residue type, applying temperature and pH corrections using the supplied ionic strength, following the approach of POTENCI.^47^ Disorder classification is performed using AlphaFold2 pLDDT scores,^49,51–53^ retaining only residues within stretches of at least 30 consecutive residues with pLDDT below 70, a conservative threshold given that AlphaFold2 is known to over-fold disordered regions.^51,63^ Experimental chemical shifts are then assigned to residues by applying Bayes’ theorem, using the residue-specific chemical shift distributions as likelihoods, with each peak treated independently without requiring sequential connectivity between peaks (see **Materials and Methods**).

SpinForecast was validated on four proteins of increasing size and complexity: islet amyloid polypeptide (IAPP, 37 residues), nuclear protein 1 (NUPR1, 82 residues), jupiter microtubule associated homolog 2 (JPT2.3, 218 residues, referred to here as JPT2),^64,65^ CBP-TAZ4 (306 residues), a construct of the CREB-binding protein containing the transcriptional-adaptor zinc-finger-2 (TAZ2) domain and the disordered region ID4.^60^ Crucially, none of the four proteins were present in the disorder-filtered BMRB dataset used to construct the residue-specific chemical shift distributions; despite being disordered, residues within IAPP and NUPR1 were not classified as disordered by our stringent pLDDT filtering criteria, and assignments for JPT2 had not been deposited in the BMRB at the time of data collection. CBP-TAZ4 was included as a stress test to assess the performance of SpinForecast on a system containing disordered and ordered domains, with the ordered domain making up approximately 30% of the total sequence.

SpinForecast was benchmarked across all possible combinations of 2 to 9 atom types using the four validation proteins. As expected, increasing the number of atom types generally increases the percentage of peaks assigned with 100% confidence (**Figure 2A**). Importantly, this increased coverage does not come at the cost of accuracy. Assignments made with 100% confidence remained at least 97% accurate when more than 4 atom types were used (**Figure 2B**). With as few as 4 atom types (CA, CB, CBi−1, Ni+1), SpinForecast achieved 100% confidence assignments for 69%, 49%, 30% and 12% of peaks in IAPP, NUPR1, JPT2 and CBP-TAZ4 respectively, all with greater than 99% accuracy. Using all 9 available atom types, the percentage of 100% confidence assignments reached 74% (IAPP), 57% (NUPR1), 44% (JPT2) and 22% (CBP-TAZ4). The lower performance of SpinForecast on CBP-TAZ4 is consistent with the expectation that residue-specific chemical shift distributions derived from disordered residues are less well-suited to the structured regions of this mixed-domain protein. The accuracy of assignments with posterior probability greater than 75% exceeded 96% across all proteins when all atom types were used (**Figure 3A**). The accuracy of the set of candidate assignments returned for each peak exceeded 97% for all IDPs (**Figure 3B**), with the median set size decreasing as more atom types are included, reflecting the increased discriminatory power of additional chemical shift information. The percentage of peaks classified as in-distribution for the best atom type combinations is shown in **Supplementary Figure 2**.

**Figure 2.**
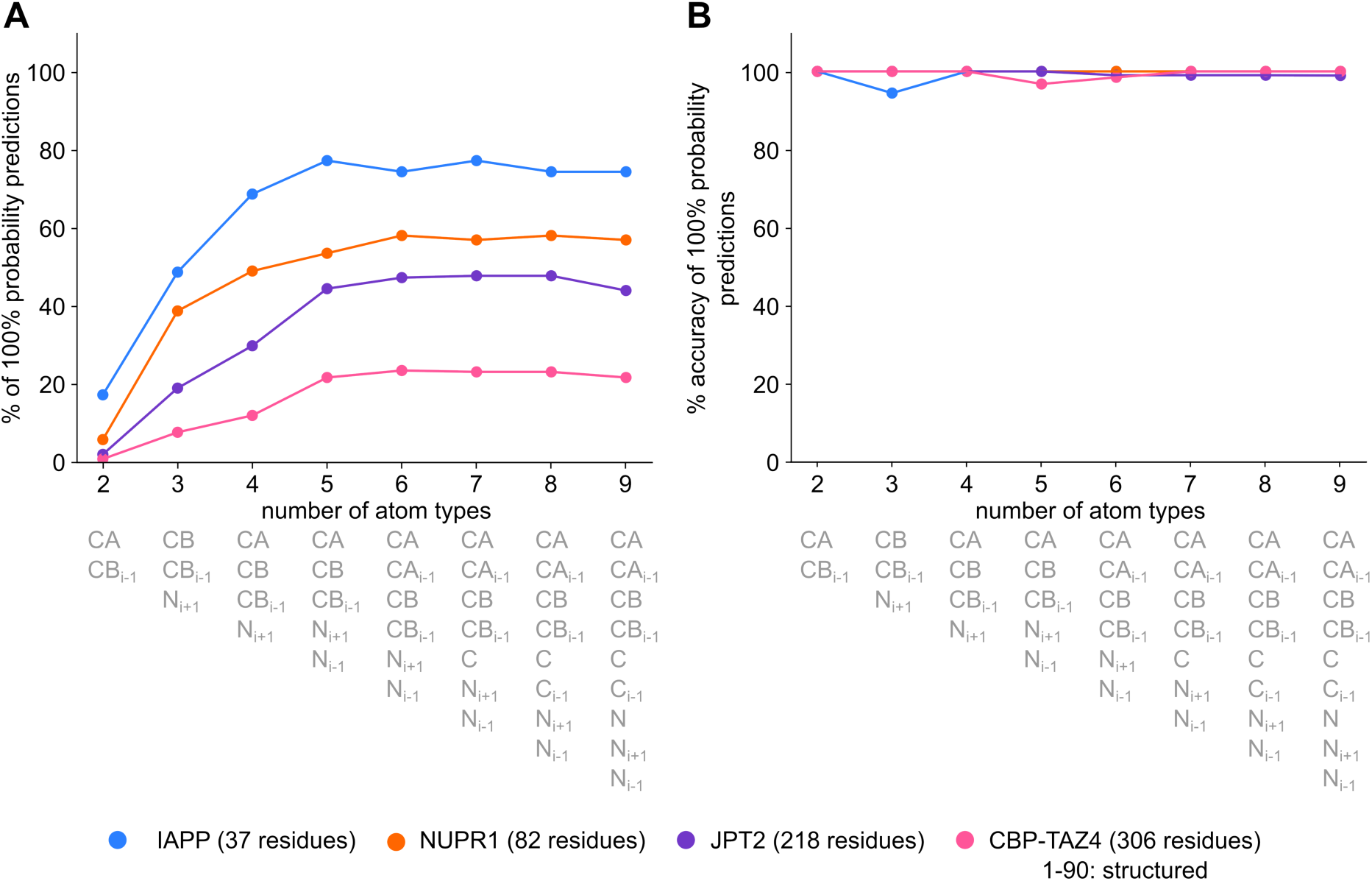
Performance statistics for assignments made by SpinForecast with 100% posterior probability when applied to IAPP, NUPR1, JPT2 and CBP-TAZ4, as a function of the number of atom types used. **A** The percentage of peaks in each peaklist assigned with 100% posterior probability as a function of the number of atom types. **B** The accuracy of those 100% posterior probability assignments as a function of the number of atom types. The atom type combinations shown below each data point show the optimal combination for that number of atom types. Here, the optimal combination was defined as the combination that maximises the sum of the percentage of peaks assigned with 100% posterior probability (panel **A**) and the accuracy of those assignments (panel **B**).

**Figure 3.**
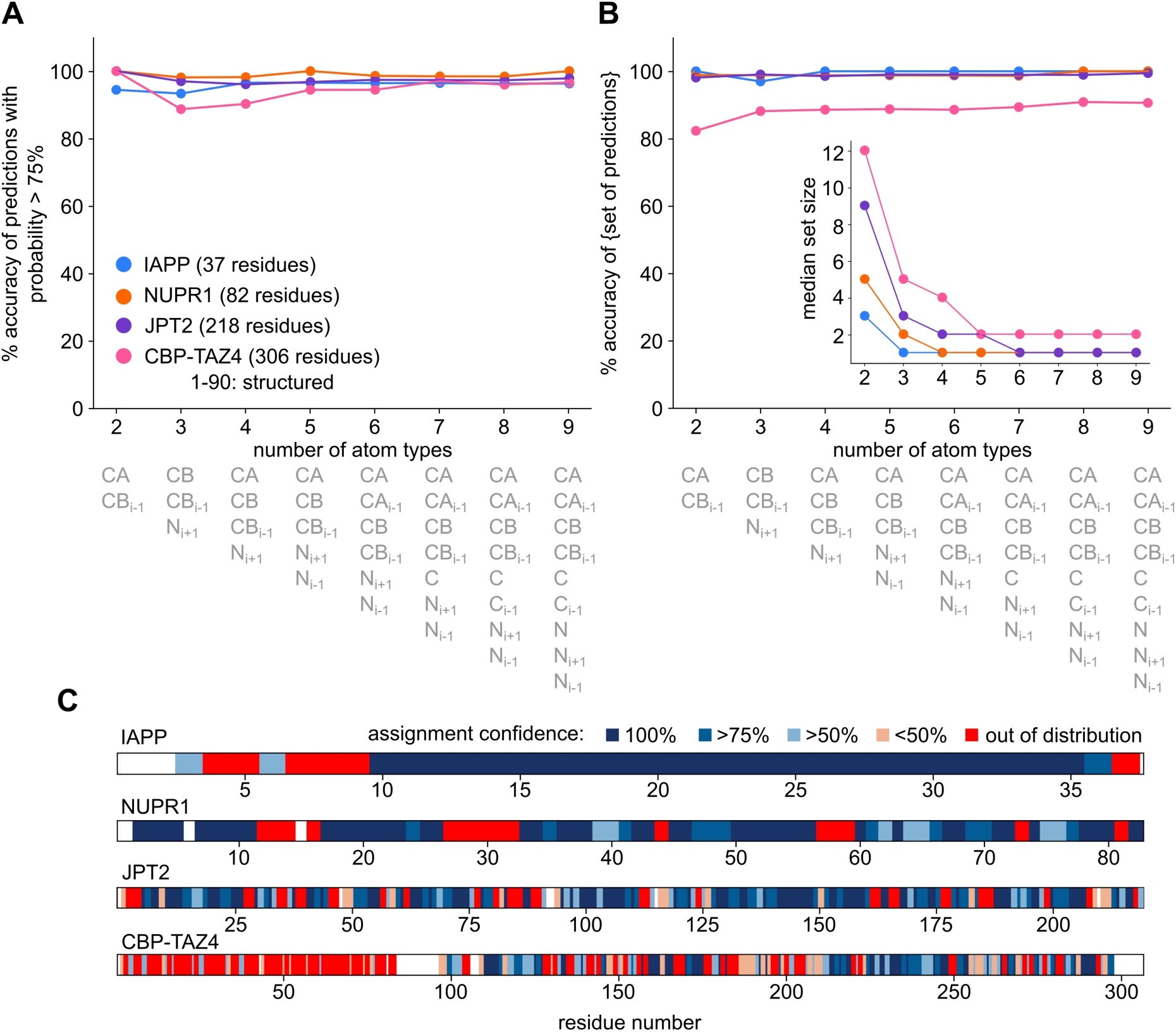
Additional performance statistics for assignments made by SpinForecast when applied to the proteins IAPP, NUPR1, JPT2 and CBP-TAZ4 as a function of the number of atom types used. **A** The accuracy of assignments with posterior probability greater than 75% as a function of number of the number of atom types. **B** The accuracy of the set of candidate assignments returned by SpinForecast as a function of number of atom types, where accuracy is calculated based on whether the correct assignment was contained within the returned set. The inset shows the median set size as a function of number of atom types; increasing the number of atom types reduces the median set size, reflecting the increased discriminatory power of additional chemical shift information. The atom type combinations shown below each data point represent the optimal combination for that number of atom types, defined as that which maximised the sum of the metrics in Figure 2A and **2B**. **C** The posterior probability of each peak assignment as a function of residue number for each validation protein when using all 9 available atom types. Peaks classified as out-of-distribution are also shown in red and unassigned residue numbers are shown as white. For CBP-TAZ4, residues 1-90 correspond to the structured zinc-finger domain.

Considering only peaks classified as in-distribution, assignments made with 100% posterior probability reached at least 99% accuracy across all validation datasets (**Figure 2B**). Where a single confident assignment could not be determined, the correct assignment was contained within the returned set of candidate assignments in greater than 97% of cases for IAPP, NUPR1, JPT2, and 82% of cases for CBP-TAZ4 (**Figure 3B**). The distribution of 100% confidence assignments across the protein sequences for each validation dataset is shown in **Figure 3C**.

Peaks classified as out-of-distribution by SpinForecast were assessed by comparing experimental CA chemical shifts to random coil values predicted by POTENCI^47^ (**Figure 4**). For IAPP, NUPR1 and CBP-TAZ4, out-of-distribution peaks showed a strong correlation with regions of secondary structure propensity, as indicated by sustained positive or negative deviations from random coil CA chemical shifts. This trend was less pronounced for JPT2, which shows little secondary structure propensity throughout its sequence.

**Figure 4.**
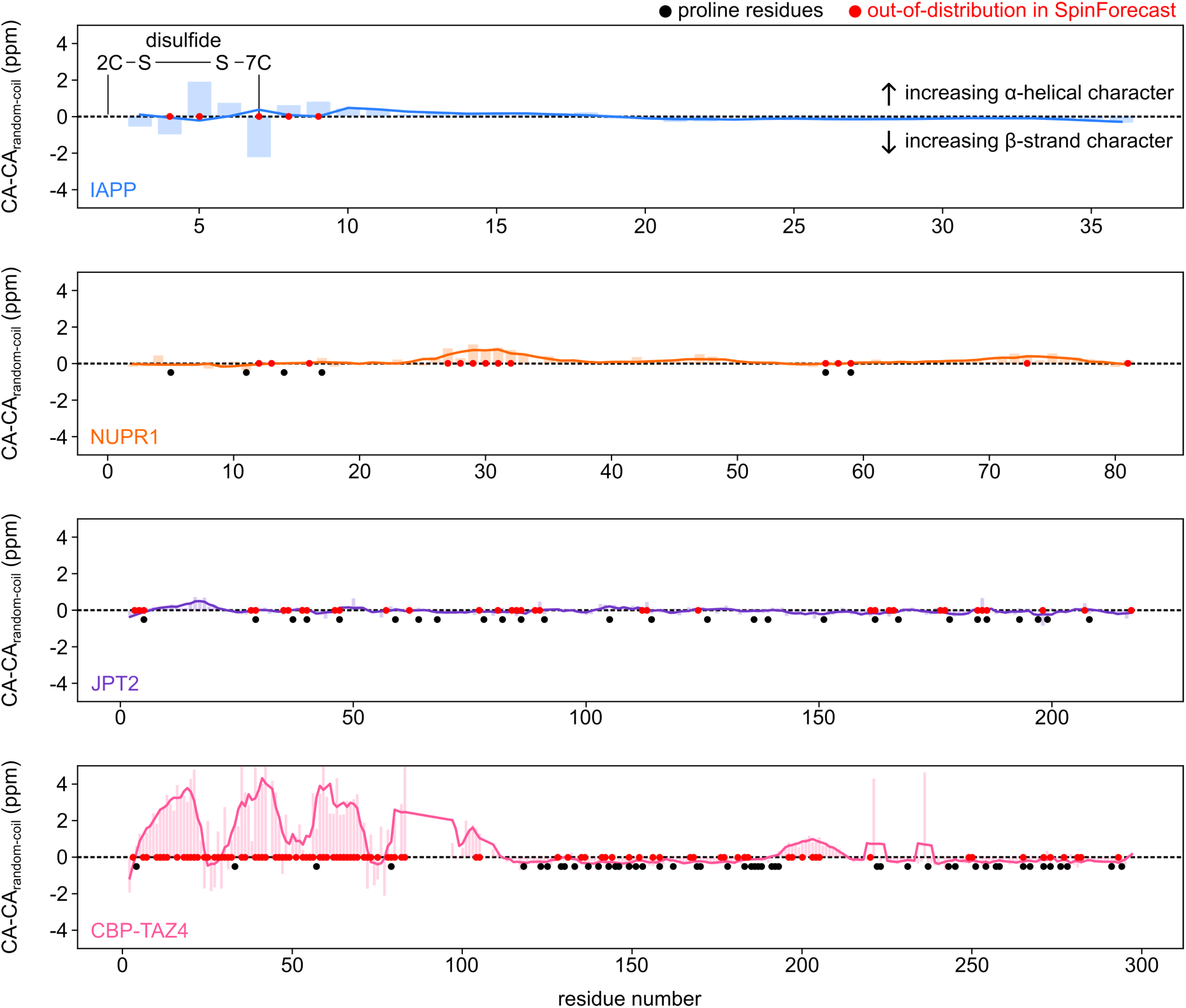
Deviations of experimental CA chemical shifts from their predicted random coil values for IAPP, NUPR1, JPT2 and CBP-TAZ4. Red circles indicate residues whose peaks were classified as out-of-distribution by SpinForecast when using all 9 available atom types. Black circles highlight residue numbers of proline amino acids. Random coil CA chemical shifts for each system were obtained from POTENCI using the sequences and experimental conditions in **Supplementary Table 1**. Sustained positive and negative deviations indicate regions with substantial α-helical and β-strand secondary structure propensity, respectively. IAPP has a disulfide bond between residues 2C and 7C which distorts the conformation of nearby residues.^71^ For IAPP, NUPR1 and CBP- TAZ4, out-of-distribution peaks (red circles) tend to coincide with regions of sustained secondary structure propensity or proximity to proline residues (black circle), though this trend does not hold for all residues. Data points are represented by bars, and trend lines were calculated using a Savitzky-Golay filter of window length 5 and a polynomial order of 1.

## Discussion and Conclusions

SpinForecast is a chain-free probabilistic tool for the backbone assignment of IDPs that exploits the narrow, well-separated chemical shift distributions of disordered residues in the BMRB. Validated on three IDPs of increasing size, IAPP (37 residues), NUPR1 (82 residues) and JPT2 (218 residues), SpinForecast achieved 100% confidence assignments for 74%, 57% and 44% of peaks respectively, all with greater than 99% accuracy. Where a single confident assignment could not be determined, the correct assignment was contained within the returned set of candidate assignments in greater than 97% of cases. Performance was also assessed on CBP-TAZ4 (306 residues), a mixed ordered/disordered system included as a stress test; as expected, performance was lower. When using all available atom types for CBP-TAZ4, only 22% of peaks could be assigned with 100% confidence and the correct assignment was contained within the candidate set in 91% of cases, reflecting the limitations of applying disordered-residue chemical shift distributions to structured regions. These results demonstrate that the sharp distributions of backbone chemical shifts in IDPs contain sufficient information to enable accurate probabilistic assignment without requiring sequential connectivity between peaks.

The application of SpinForecast to the disordered protein NUPR1 is illustrated in **Figure 5**. Starting from an unassigned CON spectrum, SpinForecast compares the experimental chemical shifts of each peak against the residue-specific chemical shift distributions for every residue in the sequence, and calculates posterior assignment probabilities using Bayes’ theorem. Peaks assigned with 100% posterior probability are shown in the output spectrum, demonstrating how SpinForecast transforms an unassigned spectrum into a partially assigned one. Critically, no connectivities between peaks are produced at any stage; each peak is assigned independently, making SpinForecast robust to missing peaks that would break the chain in conventional assignment routines.

**Figure 5.**
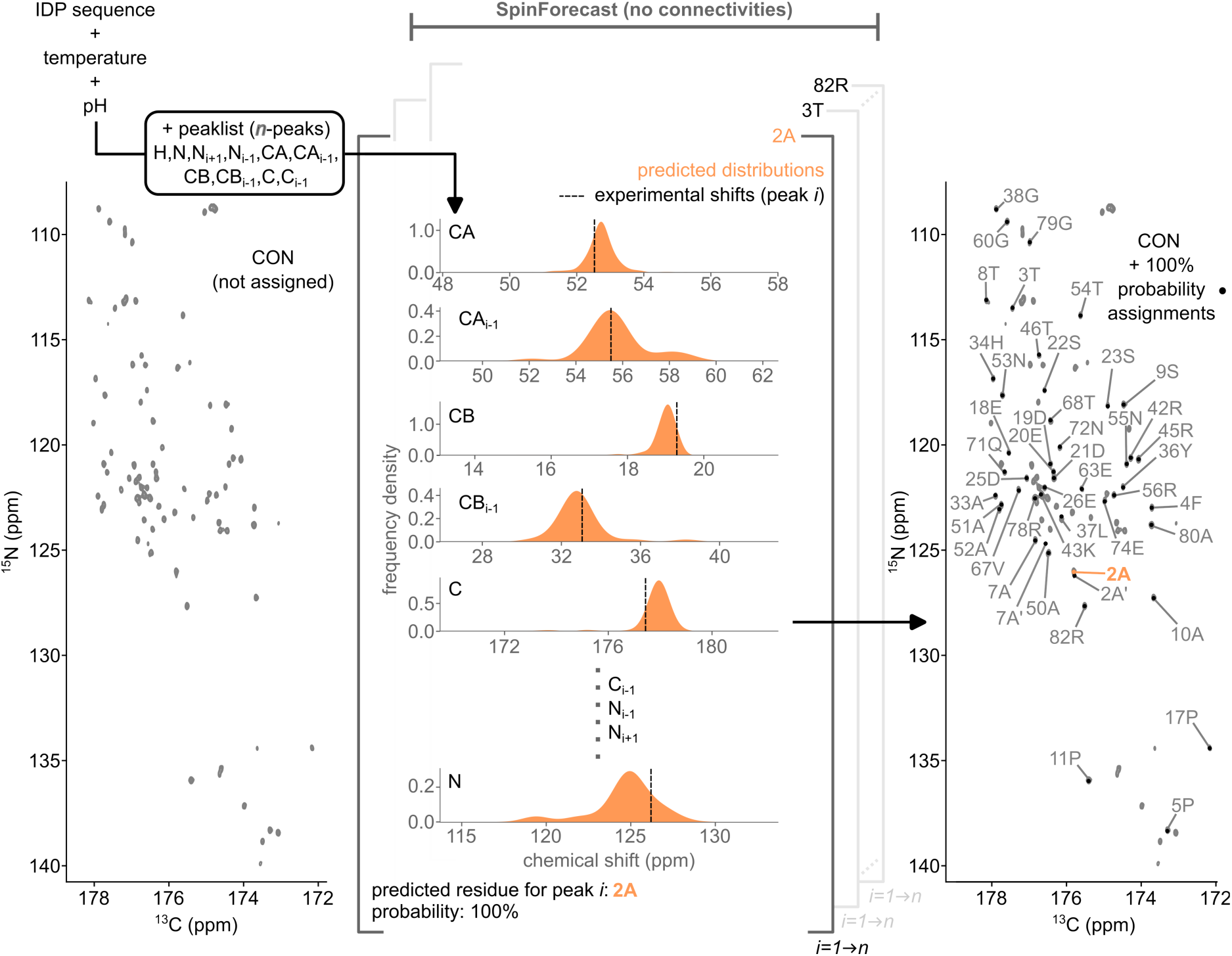
An illustration of SpinForecast applied to the 82-residue disordered protein NUPR1. Using the protein sequence, experimental conditions, and a set of N-dimensional (ND) peaklists as input, SpinForecast compares the experimental chemical shifts of each peak against the residue-specific chemical shift distributions for every residue in the sequence, calculating the posterior assignment probabilities using Bayes’ theorem. No connectivities between peaks are produced at any stage; each peak is assigned independently using only the chemical shift information contained within that entry, including any available i-1 and i+1 chemical shifts. Peaks assigned with 100% posterior probability are shown in the output spectrum on the right. In the figure, this is shown for a representative peak originating from residue 2A, which is assigned with 100% posterior probability. The peak labelled 2A’ is a minor population state arising from *cis*-proline isomerism at a nearby proline.

The 100% confidence assignments provided by SpinForecast serve as anchor points that are distributed throughout protein sequences, maximising their utility for accelerating traditional sequential assignment procedures. Rather than replacing existing assignment tools such as CCPN AnalysisAssign,^20^ MARS,^22^ or POKY,^24^ SpinForecast is designed to be used in conjunction with them; high confidence assignments established by SpinForecast provide a strong foundation from which sequential chains can be extended to fill in the remaining gaps. SpinForecast can also be used retrospectively as a validation tool, to determine whether the chemical shifts of an existing assignment are consistent with those expected for disordered residues in the BMRB. The assignment probabilities reported for each peak provide additional insight into which assignments to trust more than others, and the leave-one-out analysis identifies which atom types are driving each individual assignment prediction.

A key design principle of SpinForecast was to minimise the barrier to entry for new users. To this end, SpinForecast is freely available as an interactive online tool at https://tools.bindresearch.org/bindbox/SpinForecast, requiring no installation or programming experience (**Figure 6**). Users can input a protein sequence and experimental conditions directly into the interface and obtain residue-specific chemical shift distribution predictions, as well as interactive plots of predicted spectra for a range of experiment types. A peaklist can then be loaded by dragging and dropping onto the canvas, with support for comma separated values (.csv), tabular delimited (.tab/.list), and NMR exchange formats (.nef)^66–68^ files, as used by programs such as CCPN AnalysisAssign.^20^ The source code is openly available at https://github.com/bindresearch/BindBox, and the chemical shift distributions used by SpinForecast can be explored interactively at https://tools.bindresearch.org/bindbox/BMRB_Chemical_Shifts. By making SpinForecast freely accessible and user-friendly, we aim to encourage new researchers to study IDP targets using NMR.

**Figure 6.**
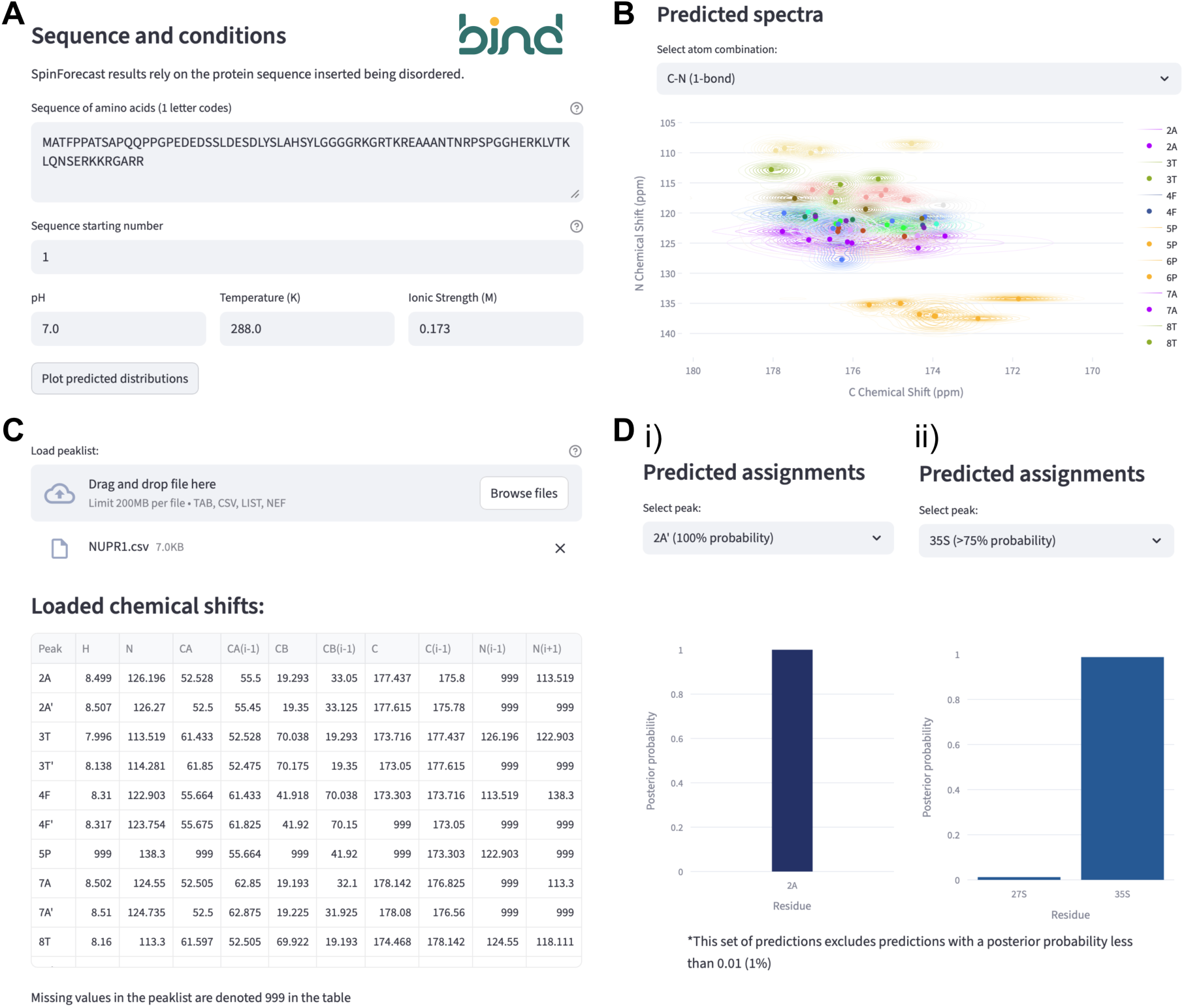
An illustration of the SpinForecast interface at https://tools.bindresearch.org/bindbox/SpinForecast when applied to the validation dataset of NUPR1. **A** The interface to input a disordered protein sequence and relevant experimental conditions. The input sequence and conditions were those used for NUPR1 throughout this paper and provided in **Supplementary Table 1**. **B** The predicted C-N (1-bond) spectrum, also known as a CON, for NUPR1 at the experimental conditions in A. Contours represent the two-dimensional (2D) KDEs for each residue and atom type combination, with scatter points at the maximum of each distribution. Predictions from different residue types are coloured differently. Other 2D predictions with different atom combinations can also be selected and plotted. **C** The interface to load a peaklist in SpinForecast, with an example from the protein NUPR1 included here (**Supplementary Table 4**). SpinForecast accepts comma separated values (.csv), tabular delimited (.tab/.list) and NMR exchange format (.nef) files. Note that peaks in this peaklist are denoted by their true assignment for illustrative purposes; these names are not used by SpinForecast to influence assignment predictions. **D** The predicted assignment of the peaks labelled 2A’ (i) and 35S (ii), from the peaklist in **Supplementary Table 4**, based on the residue-specific chemical shift distributions of NUPR1 at the experimental conditions shown in **A**. The peak 2A’ is a minor population state arising from *cis*-proline isomerism at either 5P or 6P. Each peak is assigned independently from all other peaks; using only the chemical shift information contained within that entry, including any available *i*-1 and *i*+1 chemical shifts. Panels A-D represent a selection of interface components from SpinForecast, other features are available on the webpage.

Because SpinForecast assigns each peak independently without requiring sequential connectivities, it is particularly well-suited to aid the assignment of low-intensity minor population states where it is often more difficult to find connectivities to other peaks in the spectrum. This is especially relevant for IDPs, which are enriched in proline residues relative to structured proteins^15^ and therefore have an increased potential for peaks arising from *cis*-proline conformations. Proline *cis* and *trans* conformations can be distinguished by their CB-CG chemical shifts, with values greater than 8.0 ppm indicating *cis*-proline and values less than 8.0 ppm indicating *trans*-proline.^16,69,70^ Calculation of the CB-CG chemical shifts for all proline residues where both CB and CG chemical shifts were reported in the disordered and structured chemical shift databases revealed that 4.43% of deposited structured proline residues adopt the *cis* conformation, reducing to 1.95% for disordered residues (**Supplementary Figure 3**). Whether this lower proportion reflects a genuine feature of IDPs or a lack of deposited data for *cis*-proline conformations in IDPs remains unclear. We anticipate that SpinForecast will facilitate the assignment of these minor population states, enabling more of them to be deposited in the BMRB and allowing their potential role in the behaviour of IDPs to be uncovered.

Peaks classified as out-of-distribution by SpinForecast showed a strong correlation with either regions of secondary structure propensity, or proximity to proline residues, in IAPP,^71^ NUPR1, JPT2 and CBP-TAZ4^60^ (**Figure 4**). The correlation with secondary structure propensity is perhaps unsurprising given that NMR-derived secondary structure propensity is commonly assessed by comparing experimental CA and CB chemical shifts to random coil values.^14,47,72–76^ Programs that assess secondary structure based on experimental chemical shifts such as the TALOS,^77^ SSP,^78^ CheZOD Z-score,^79^ TriZOD G-score^80^ heavily rely on these comparisons with random coil predictions. Residues with significant secondary structure propensity will therefore deviate from the disordered chemical shift distributions used by SpinForecast, leading to their classification as out-of-distribution. This behaviour is particularly pronounced in CBP-TAZ4, where the structured zinc-finger domain produces consecutive stretches of out-of-distribution peaks corresponding to regions of sustained secondary structure. The out-of-distribution classification can therefore provide a useful diagnostic for identifying regions of residual or stable secondary structure within a protein, complementing existing secondary structure propensity analyses such as those based on POTENCI.^47^ We also noticed that close proximity (within two positions) to proline residues was an additional contributor to out-of-distribution classifications. There are several potential contributing factors: 1) the chain-breaking nature of prolines results in an underrepresention of this disordered amino acid and its nearest neighbours in the disordered chemical shift dataset extracted from the BMRB, 2) SpinForecast only considers the *i*±1 nearest neighbours when filtering our disordered dataset. This results in the KDEs for prolines and their nearest neighbors (*i*±{1,2}) becoming artificially narrow leading to enhanced out-of-distribution classifications. We anticipate this will improve as more disordered chemical shift data is deposited in the BMRB.

The disorder classification used to filter the BMRB dataset relies on AlphaFold2 pLDDT scores,^49,53^ with a conservative threshold of pLDDT below 70 applied to stretches of at least 30 consecutive residues. Given that AlphaFold2 is known to over-fold disordered regions,^51,63^ this threshold was chosen to ensure that retained chemical shifts are genuinely disordered; however, as AlphaFold2 and other structure prediction tools continue to improve their handling of disordered regions, this threshold may warrant revisiting in future iterations of SpinForecast.

SpinForecast builds upon a rich body of previous work exploiting chemical shift statistics of IDPs for assignment purposes. Early work by Piai et al.^33^ and Romero et al.^34^ demonstrated that chemical shift values alone contain sufficient information for amino acid recognition in IDPs, and similar approaches using chemical shifts for common dipeptides in the IDPome have recently been reported as a method to aid assignment.^81^ SpinForecast extends these approaches by applying a substantially larger dataset of disorder-filtered chemical shifts and filtering by nearest-neighbour residue type, rather than residue type alone. Obtaining chemical shift information for additional atoms along the backbone collapses the probability distributions to reduce the number of candidate peak assignments that agree with the chemical shift data observed (**Figures 2** and **3**). Overall, this increases the number of 100% confident assignments along with their accuracy (**Figure 2**). Extending SpinForecast in the future to include chemical shifts of additional side-chain nuclei not currently implemented would further increase the number of 100% confident assignments obtained. Utilising higher dimensional (≥4D) experiments would enable chemical shift information for many atoms to be linked and classified to a particular peak automatically.^27–34^ The obtained peak list could then be instantly inserted into SpinForecast to assess assignment candidates and probabilities. Amino acid selective experiments could then be employed to resolve any remaining ambiguous assignments.^33^

The dataset used by SpinForecast contains 217,357 re-referenced backbone chemical shifts from 847 unique sequences and 957 BMRB entries, compared to 47,757 chemical shifts from 137 proteins used to train POTENCI.^47^ Like POTENCI, SpinForecast applies temperature and pH corrections to account for experimental conditions; however, rather than returning a single point estimate for each residue, SpinForecast returns a full distribution of chemical shifts, enabling probabilistic assignment by Bayes’ theorem. SpinForecast also reports uncertainties for each prediction, providing the user with a measure of confidence in the returned distributions that is not available from single-point predictors such as POTENCI.

SpinForecast represents a new approach to the assignment of IDPs by NMR, exploiting the largely untapped chemical shift data held in the BMRB to enable chain-free probabilistic assignment from chemical shifts alone. As the BMRB continues to grow with new depositions of disordered protein chemical shifts, the residue-specific distributions used by SpinForecast will become increasingly precise, improving assignment performance for a wider range of sequence contexts and experimental conditions. Similarly, continued development of disorder prediction tools beyond pLDDT-based classification^80,82^ may allow more accurate and comprehensive filtering of the BMRB in future iterations of SpinForecast. By making such tools freely accessible, we aim to lower the barrier to entry for protein NMR assignment and encourage new researchers to study IDPs using solution-state NMR spectroscopy.

## Author Contributions

GTH and GK conceptualised the project. GK filtered the BMRB chemical shift database by pLDDT score. JTE and GK wrote the SpinForecast program and performed benchmarking. JC, JTE, and GTH performed the assignment of NUPR1. KMS, JC, JTE, and GTH performed the assignment of JPT2. TL and JTE productionised SpinForecast and its online architecture. PC and JC provided helpful suggestions on SpinForecast features. JTE, GK, and GTH wrote the manuscript with input from all authors.

## Acknowledgements

The authors thank Sören von Bülow (Bind Research), Charles J. Buchanan (University College London, Max-Planck-Institute for Terrestrial Microbiology), T. Reid Alderson (Helmholtz Munich), D. Flemming Hansen (University College London), Christopher A. Waudby (University College London), Nicolas L Fawzi (Brown University), Sandy Wang (Brown University), and Tongyin Zheng (Brown University) for helpful discussions in the development of SpinForecast. The authors acknowledge Jonathan Marchant (Medical College of Wisconsin), Sushil Kumar (Medical College of Wisconsin) and Sandip Patel (University College London) for helpful discussion about the purification of JPT2. The authors acknowledge Geoff Kelly, Peter J Simpson and Alain Oregioni from the Francis Crick Institute for their assistance in recording NMR assignment spectra. Bind Research is supported by the UK Government via the Department for Science, Innovation and Technology’s Research Ventures Catalyst Programme, Eric and Wendy Schmidt, and the Klaff Family Foundation. GTH is supported by a Biotechnology and Biological Sciences Research Council (BBSRC) Discovery Fellowship (BB/X009955/1). KS was supported by the University College London-Birkbeck Medical Research Council Doctoral Training Programme (MR/W006774/1). JC was supported by a Steel Perlot Early Investigator Grant and by Erika Alden DeBenedictis (Francis Crick Institute). NMR assignment experiments were supported by the Francis Crick Institute through provision of access to the MRC Biomedical NMR Centre. The Francis Crick Institute receives its core funding from Cancer Research UK (CC1078); UK Medical Research Council (CC1078); Wellcome Trust (CC1078). For the purpose of open access, the authors have applied a Creative Commons Attribution (CC BY) licence to any Author Accepted Manuscript version arising from this submission.

## Data availability

The source code for SpinForecast at the time of writing is available on https://github.com/bindresearch/BindBox/releases/tag/v1.1. The chemical shift distributions of disordered and structured proteins are available in the .parquet files within the GitHub repository. The latest version of SpinForecast can be accessed online at https://tools.bindresearch.org/bindbox/SpinForecast. The chemical shift distributions can also be explored interactively at https://tools.bindresearch.org/bindbox/BMRB_Chemical_Shifts, noting that this page may be updated as new BMRB data are released. A video tutorial is available at https://www.youtube.com/@BindResearch. Chemical shift assignments of NUPR1 and JPT2 are available in **Supplementary Tables 4** and **5**, and are deposited in the BMRB (accession numbers 53937 and 53938, respectively). Unprocessed NMR data and processing scripts for NUPR1 and JPT2 are included in their respective BMRB submissions.

## Notes

https://tools.bindresearch.org/bindbox/SpinForecast

https://tools.bindresearch.org/bindbox/BMRB_Chemical_Shifts

https://github.com/bindresearch/BindBox/

https://www.youtube.com/@BindResearch

### Competing Interest Statement

The authors have declared no competing interest.

