## Supplementary Materials for "SpinForecast: chain-free probabilistic backbone assignment of intrinsically disordered proteins from NMR chemical shifts"

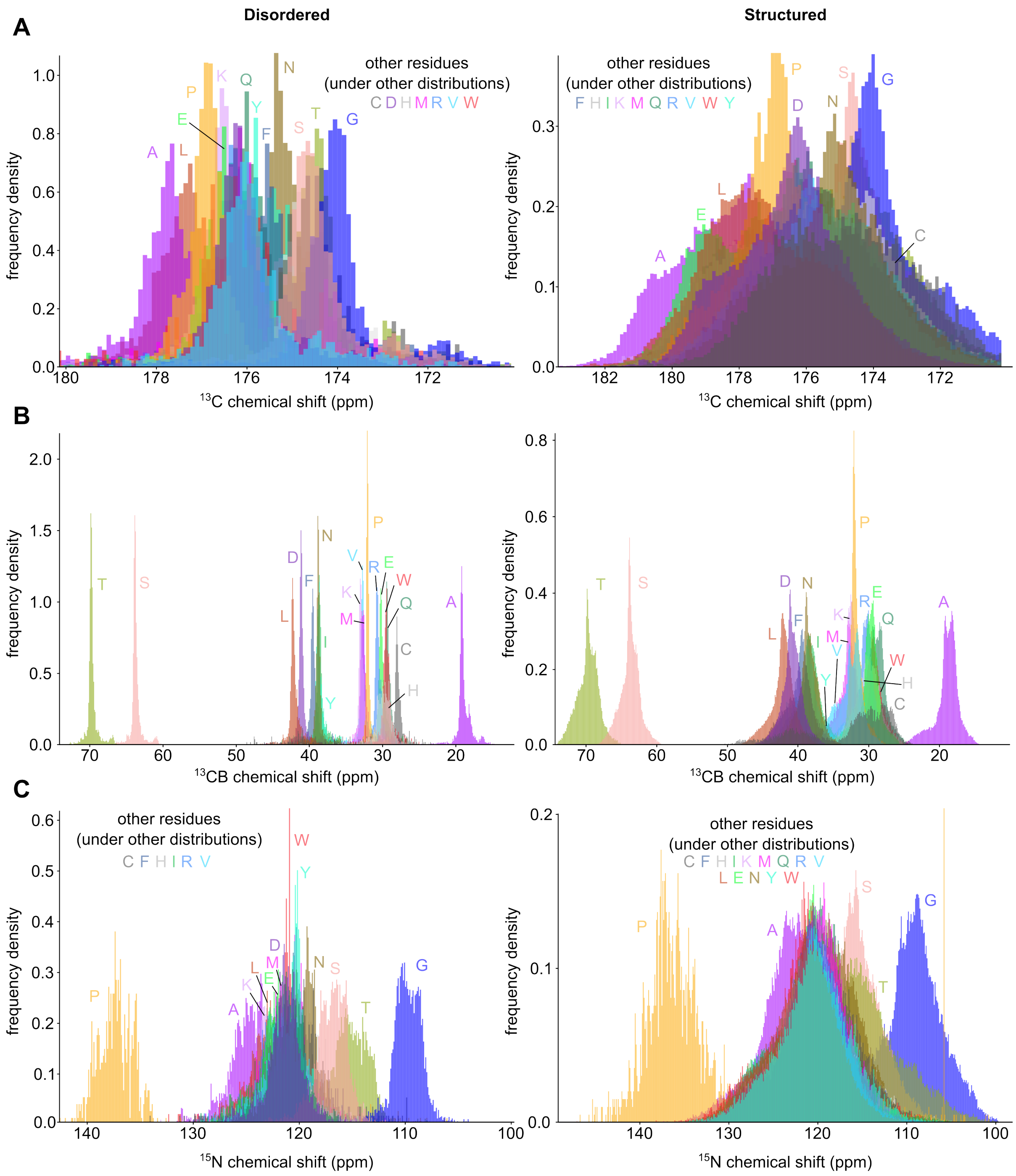


**Supplementary Figure 1** Overlays of C (**A**), CB (**B**) and N (**C**) chemical shift distributions for all amino acid types for disordered residues (left) and structured residues (right) within the BMRB database. Disordered chemical shifts were extracted using the disorder classification procedure described in the Materials and Methods, and structured chemical shifts contain all values extracted from the BMRB not classified as disordered. The plots were produced using the dashboard at https://tools.bindresearch.org/bindbox/BMRB_Chemical_Shifts using histogram bin widths of 0.1 ppm.

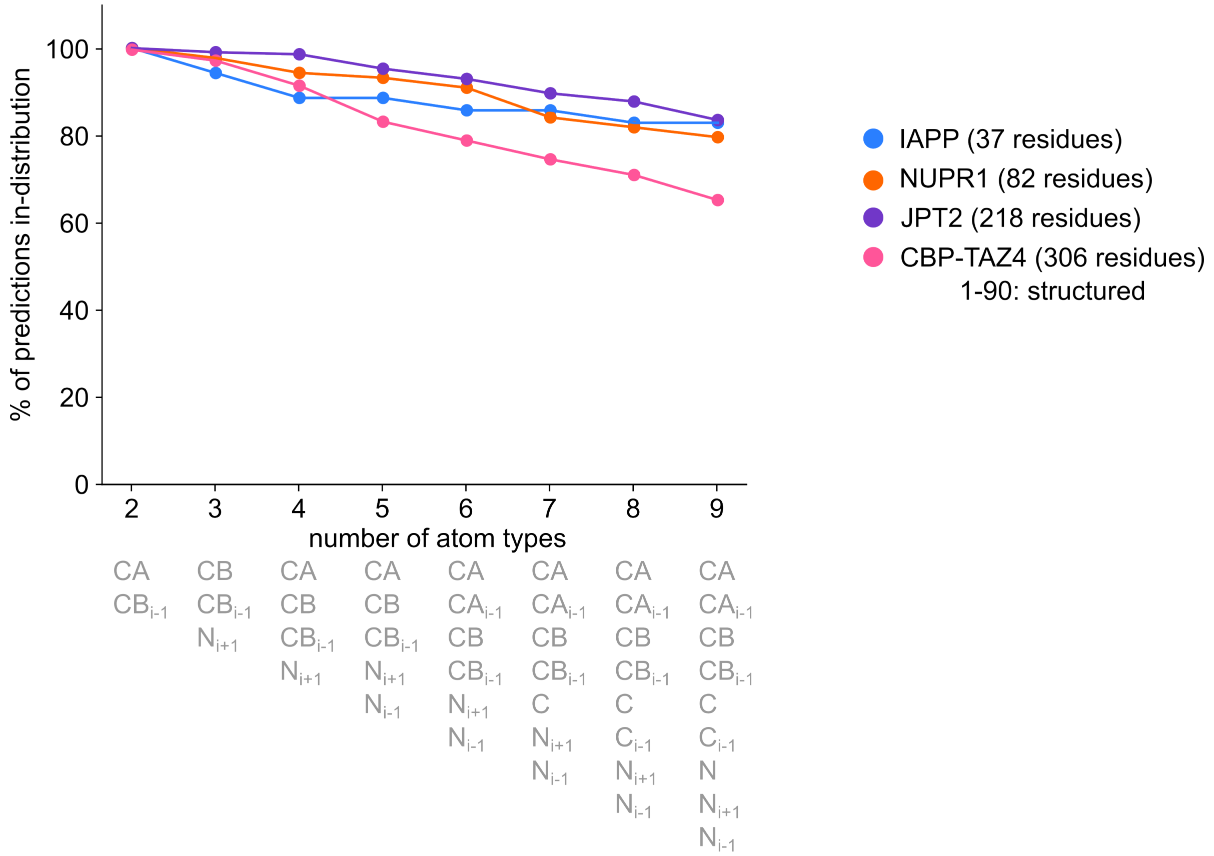


**Supplementary Figure 2** The percentage of assignment predictions made by SpinForecast that were classified as in-distribution as a function of the number of atom types used. The percentage of in-distribution predictions decreases as more atom types are included, reflecting the increasingly stringent requirements for a peak to be consistent with the residue-specific chemical shift distributions across all atom types. The atom type combinations shown below each data point give the optimal combination for that number of atom types. The optimal combination is defined as the one that maximises the sum of the percentage of 100% probability predictions and the accuracy of those predictions.

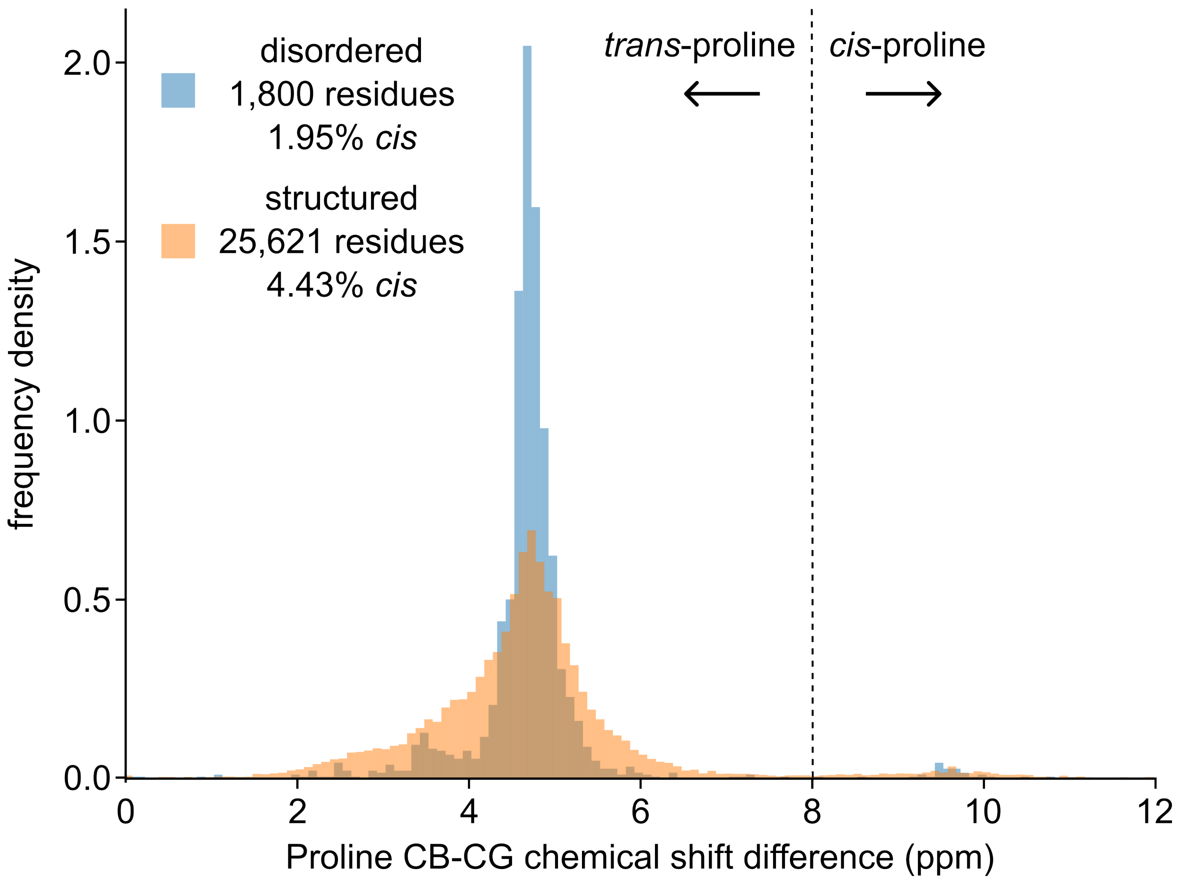


**Supplementary Figure 3** Distributions of CB-CG chemical shift values for disordered (1,800 residues) and structured (25,621 residues) proline residues in the BMRB (as of August 2025). Note that the set of chemical shifts for this analysis was un-referenced as the difference between CB and CG shifts is independent of any referencing error. Values of CB-CG greater than 8.0 ppm were classified as originating from *cis*-proline conformations, following established criteria.[1,2] The percentage of reported values in the BMRB classified as *cis* are shown for both structured and disordered residues. The histogram bin widths were set to 0.1 ppm.

**Supplementary Table 1** A table of proteins used to test the performance of SpinForecast. For the ionic strength calculations, it was assumed that the sample was prepared and pH corrected at a temperature of 298K. The ionic strengths were calculated by combining the contributions from salts and the anionic/cationic buffer contribution, calculated using the Henderson-Hasselbalch equation (assuming the presence of monovalent anionic/cationic counterions to achieve charge conservation with a net zero charge).[3] For sodium acetate in the CBP-TAZ4 buffer, it was assumed that any loss of negative charge from the acetate component upon pH correction to 5.5 was replaced by the equivalent amount of another monovalent negative counterion in order to reach overall charge neutrality. Any additional cations or anions added during pH correction were unknown and neglected. The following pKa values were used: 7.55 (HEPES),[4,] 6.27 (MES),[4] 8.06 (Tris),[4] 4.76 (acetate),[4] assuming a temperature of 298K.

| Protein name | BMRB ID | Temp. (K) | pH | Buffer components | Ionic strength (M) | Sequence |
|---|---|---|---|---|---|---|
| **IAPP** | 51259[5] | 283 | 6.5 | 25 mM MES, 25 mM $MgCl_2$ | 0.091 | KCNTATCATQRLANFLVHSSNNFG AILSSTNVGSNTY |
| **NUPR1** | 53937 | 288 | 7.0 | 25 mM Tris, 150 mM NaCl, 1 mM TCEP, 0.05% $NaN_3$ | 0.173 | MATFPPATSAPQQPPGPEDEDSSLD ESDLYSLAHSYLGGGGRKGRTKRE AAANTNRPSPGGHERKLVTKLQNS ERKKRGARR |
| **JPT2** | 53938 | 278 | 7.2 | 20 mM HEPES, 110 mM KCl, 10 mM NaCl, 1 mM TCEP | 0.126 | MFQVPDSEGGRAGSRKWQLLTGS LASTSPSLLSGQGPWAPLQRAMKP PGGESSNLFGSPEEATPSSRPNRMA SNIFGPTEEPQNIPKRTNPPGGKGS GIFDESTPVQTRQHLNPPGGKTSDI FGSPVTATSRLAHPNKPKDHVFLC EGEEPKSDLKAARSIPAGAEPGEKG SARKAGPAKEQEPMPTVDSHEPRL GPRPRSHNKVLNPPGGKSSISFY |
| **CBP-TAZ4** | 52655[6] | 298 | 5.5 | 25 mM sodium acetate, 50 mM NaCl, 0.1 mM $ZnCl_2$, 2 mM TCEP, 0.05% $NaN_3$ | 0.075 | MGSPQESRRLSIQRAIQSLVHAAQC RNANCSLPSCQKMKRVVQHTKGC KRKTNGGCPVCKQLIALAAYHAK HCQENKCPVPFCLNIKHKLRQQQI QHRLQQAQLMRRRMATMNTRNV PQQSLPSPTSAPPGTPTQQPSTPQTP QPPAQPQPSPVSMSPAGFPSVARTQ PPTTVSTGKPTSQVPAPPPPAQPPP AAVEAARQIEREAQQQQHLYRVNI NNSMPPGRTGMGTPGSQMAPVSL NVPRPNQVSGPVMPSMPPGQWQQ APLPQQQPMPGLPRPVISMQAQAA VAGPRMPSVQQLEHHHHHH |

**Supplementary Table 2** SpinForecast assignment performance metrics for the optimal atom combination for each number of atom types used.

| Protein name | Number of atom types | Atom types | Proportion of predictions with set size = 1 | Accuracy (posterior probability = 1) | Accuracy (posterior probability > 75%) | Accuracy of candidate assignment set | Median set size | Proportion in-distribution | Residue numbers out of distribution |
|---|---|---|---|---|---|---|---|---|---|
| **IAPP** | 2 | ['CA', 'CB(i-1)'] | 0.17 | 1.00 | 0.94 | 1.00 | 3.0 | 1.00 | [] |
| **NUPR1** | 2 | ['CA', 'CB(i-1)'] | 0.06 | 1.00 | 1.00 | 0.99 | 5.0 | 1.00 | [] |
| **JPT2** | 2 | ['CA', 'CB(i-1)'] | 0.02 | 1.00 | 1.00 | 0.98 | 9.0 | 1.00 | [] |
| **CBP_TAZ4** | 2 | ['CA', 'CB(i-1)'] | 0.01 | 1.00 | 1.00 | 0.82 | 12.0 | 1.00 | ['42'] |
| **IAPP** | 3 | ['CB', 'CB(i-1)', 'N(i+1)'] | 0.49 | 0.94 | 0.93 | 0.97 | 1.0 | 0.94 | ['4', '5'] |
| **NUPR1** | 3 | ['CB', 'CB(i-1)', 'N(i+1)'] | 0.39 | 1.00 | 0.98 | 0.99 | 2.0 | 0.98 | ['3', '4'] |
| **JPT2** | 3 | ['CB', 'CB(i-1)', 'N(i+1)'] | 0.19 | 1.00 | 0.97 | 0.99 | 3.0 | 0.99 | ['39', '217'] |
| **CBP_TAZ4** | 3 | ['CB', 'CB(i-1)', 'N(i+1)'] | 0.08 | 1.00 | 0.89 | 0.88 | 5.0 | 0.97 | ['11', '12', '19', '27', '35', '53', '64', '65'] |
| **IAPP** | 4 | ['CA', 'CB', 'CB(i-1)', 'N(i+1)'] | 0.69 | 1.00 | 0.97 | 1.00 | 1.0 | 0.89 | ['4', '5', '6', '8'] |
| **NUPR1** | 4 | ['CA', 'CB', 'CB(i-1)', 'N(i+1)'] | 0.49 | 1.00 | 0.98 | 0.99 | 1.0 | 0.94 | ['3', '4', '12', '30', '58'] |
| **JPT2** | 4 | ['CA', 'CB', 'CB(i-1)', 'N(i+1)'] | 0.30 | 1.00 | 0.96 | 0.99 | 2.0 | 0.99 | ['39', '176', '217'] |
| **CBP_TAZ4** | 4 | ['CA', 'CB', 'CB(i-1)', 'N(i+1)'] | 0.12 | 1.00 | 0.90 | 0.89 | 4.0 | 0.91 | ['6', '10', '12', '13', '16', '17', '27', '28', '29', '35', '41', '42', '44', '45', '46', '47', '53', '54', '63', '64', '77', '83', '105', '135'] |
| **IAPP** | 5 | ['CA', 'CB', 'CB(i-1)', 'N(i+1)', 'N(i-1)'] | 0.77 | 1.00 | 0.97 | 1.00 | 1.0 | 0.89 | ['4', '5', '7', '8'] |
| **NUPR1** | 5 | ['CA', 'CB', 'CB(i-1)', 'N(i+1)', 'N(i-1)'] | 0.53 | 1.00 | 1.00 | 0.99 | 1.0 | 0.93 | ['3', '4', '12', '30', '57', '58'] |
| **JPT2** | 5 | ['CA', 'CB', 'CB(i-1)', 'N(i+1)', 'N(i-1)'] | 0.44 | 1.00 | 0.97 | 0.99 | 2.0 | 0.95 | ['35', '39', '40', '62', '89', '165', '176', '184', '185', '217'] |

| Protein name | Number of atom types | Atom types | Proportion of predictions with set size = 1 | Accuracy (posterior probability = 1) | Accuracy (posterior probability > 75%) | Accuracy of candidate assignment set | Median set size | Proportion in-distribution | Residue numbers out of distribution |
|---|---|---|---|---|---|---|---|---|---|
| **CBP_TAZ4** | 5 | ['CA', 'CB', 'CB(i-1)', 'N(i+1)', 'N(i-1)'] | 0.22 | 0.97 | 0.94 | 0.89 | 2.0 | 0.83 | ['6', '7', '10', '12', '13', '16', '17', '22', '24', '25', '27', '28', '29', '30', '33', '35', '41', '42', '44', '45', '46', '47', '48', '50', '51', '52', '53', '54', '55', '56', '63', '64', '65', '68', '71', '77', '83', '131', '135', '141', '158', '167', '183', '184', '265', '273', '281'] |
| **IAPP** | 6 | ['CA', 'CA(i-1)', 'CB', 'CB(i-1)', 'N(i+1)', 'N(i-1)'] | 0.74 | 1.00 | 0.96 | 1.00 | 1.0 | 0.86 | ['4', '5', '7', '8', '9'] |
| **NUPR1** | 6 | ['CA', 'CA(i-1)', 'CB', 'CB(i-1)', 'N(i+1)', 'N(i-1)'] | 0.58 | 1.00 | 0.99 | 0.99 | 1.0 | 0.91 | ['3', '4', '12', '16', '30', '57', '58', '73'] |
| **JPT2** | 6 | ['CA', 'CA(i-1)', 'CB', 'CB(i-1)', 'N(i+1)', 'N(i-1)'] | 0.47 | 0.99 | 0.97 | 0.99 | 1.0 | 0.93 | ['5', '35', '36', '39', '40', '62', '89', '165', '166', '176', '177', '184', '185', '217', '218'] |
| **CBP_TAZ4** | 6 | ['CA', 'CA(i-1)', 'CB', 'CB(i-1)', 'N(i+1)', 'N(i-1)'] | 0.23 | 0.98 | 0.94 | 0.89 | 2.0 | 0.79 | ['6', '7', '8', '11', '12', '16', '18', '19', '21', '22', '24', '27', '28', '29', '31', '33', '35', '36', '40', '42', '44', '45', '46', '48', '50', '51', '52', '53', '54', '55', '56', '57', '60', '62', '63', '64', '66', '67', '68', '69', '71', '79', '83', '103', '131', '135', '138', '141', '156', '158', '167', '181', '183', '184', '265', '271', '273', '281', '293'] |
| **IAPP** | 7 | ['CA', 'CA(i-1)', 'CB', 'CB(i-1)', 'N(i+1)', 'N(i-1)', 'C'] | 0.77 | 1.00 | 0.96 | 1.00 | 1.0 | 0.86 | ['4', '5', '7', '8', '9'] |
| **NUPR1** | 7 | ['CA', 'CA(i-1)', 'CB', 'CB(i-1)', 'N(i+1)', 'N(i-1)', 'C'] | 0.57 | 1.00 | 0.98 | 0.99 | 1.0 | 0.84 | ['3', '4', '12', '13', '16', '27', '29', '30', '31', '32', '57', '58', '73', '81'] |

| Protein name | Number of atom types | Atom types | Proportion of predictions with set size = 1 | Accuracy (posterior probability = 1) | Accuracy (posterior probability > 75%) | Accuracy of candidate assignment set | Median set size | Proportion in-distribution | Residue numbers out of distribution |
|---|---|---|---|---|---|---|---|---|---|
| **JPT2** | 7 | ['CA', 'CA(i-1)', 'CB', 'CB(i-1)', 'N(i+1)', 'N(i-1)', 'C'] | 0.48 | 0.99 | 0.97 | 0.99 | 1.0 | 0.90 | ['3', '5', '35', '36', '39', '40', '46', '57', '62', '84', '89', '90', '112', '124', '165', '166', '176', '177', '184', '185', '217', '218'] |
| **CBP_TAZ4** | 7 | ['CA', 'CA(i-1)', 'CB', 'CB(i-1)', 'N(i+1)', 'N(i-1)', 'C'] | 0.23 | 1.00 | 0.97 | 0.89 | 2.0 | 0.74 | ['6', '7', '8', '11', '12', '16', '18', '19', '21', '24', '27', '28', '29', '31', '32', '35', '36', '38', '40', '42', '45', '46', '47', '48', '50', '51', '52', '53', '54', '55', '56', '58', '59', '62', '63', '64', '66', '67', '68', '69', '71', '72', '73', '75', '79', '83', '104', '128', '131', '135', '141', '156', '158', '168', '176', '181', '183', '184', '185', '196', '197', '203', '205', '220', '222', '249', '265', '271', '273', '281', '293'] |
| **IAPP** | 8 | ['CA', 'CA(i-1)', 'CB', 'CB(i-1)', 'N(i+1)', 'N(i-1)', 'C', 'C(i-1)'] | 0.74 | 1.00 | 0.96 | 1.00 | 1.0 | 0.83 | ['4', '5', '7', '8', '9', '37'] |
| **NUPR1** | 8 | ['CA', 'CA(i-1)', 'CB', 'CB(i-1)', 'N(i+1)', 'N(i-1)', 'C', 'C(i-1)'] | 0.58 | 1.00 | 0.98 | 1.00 | 1.0 | 0.82 | ['3', '4', '12', '13', '14', '16', '27', '28', '29', '30', '31', '32', '57', '58', '73', '81'] |
| **JPT2** | 8 | ['CA', 'CA(i-1)', 'CB', 'CB(i-1)', 'N(i+1)', 'N(i-1)', 'C', 'C(i-1)'] | 0.48 | 0.99 | 0.97 | 0.99 | 1.0 | 0.88 | ['3', '5', '35', '36', '39', '40', '46', '57', '62', '77', '84', '85', '89', '90', '112', '113', '124', '165', '166', '176', '177', '184', '185', '207', '217', '218'] |

| Protein name | Number of atom types | Atom types | Proportion of predictions with set size = 1 | Accuracy (posterior probability = 1) | Accuracy (posterior probability > 75%) | Accuracy of candidate assignment set | Median set size | Proportion in-distribution | Residue numbers out of distribution |
|---|---|---|---|---|---|---|---|---|---|
| **CBP_TAZ4** | 8 | ['CA', 'CA(i-1)', 'CB', 'CB(i-1)', 'N(i+1)', 'N(i-1)', 'C', 'C(i-1)'] | 0.23 | 1.00 | 0.96 | 0.91 | 2.0 | 0.71 | ['6', '7', '8', '10', '11', '12', '16', '18', '19', '21', '22', '24', '27', '28', '29', '30', '31', '32', '35', '36', '38', '39', '40', '41', '42', '45', '46', '47', '48', '51', '53', '54', '55', '56', '58', '59', '62', '63', '64', '65', '66', '67', '68', '69', '71', '72', '73', '75', '77', '78', '79', '83', '104', '128', '131', '135', '141', '156', '158', '167', '168', '169', '176', '181', '183', '184', '185', '196', '197', '200', '203', '204', '205', '220', '249', '250', '265', '271', '273', '281', '293'] |
| **IAPP** | 9 | ['CA', 'CA(i-1)', 'CB', 'CB(i-1)', 'N(i+1)', 'N(i-1)', 'C', 'C(i-1)', 'N'] | 0.74 | 1.00 | 0.96 | 1.00 | 1.0 | 0.83 | ['4', '5', '7', '8', '9', '37'] |
| **NUPR1** | 9 | ['CA', 'CA(i-1)', 'CB', 'CB(i-1)', 'N(i+1)', 'N(i-1)', 'C', 'C(i-1)', 'N'] | 0.57 | 1.00 | 1.00 | 1.00 | 1.0 | 0.80 | ['3', '4', '12', '13', '14', '16', '27', '28', '29', '30', '31', '32', '57', '58', ’58'', '59', '73', '81'] |
| **JPT2** | 9 | ['CA', 'CA(i-1)', 'CB', 'CB(i-1)', 'N(i+1)', 'N(i-1)', 'C', 'C(i-1)', 'N'] | 0.44 | 0.99 | 0.98 | 0.99 | 1.0 | 0.83 | ['3', '4', '5', '28', '29', '35', '36', '39', '40', '46', '47', '57', '62', '77', '81', '84', '85', '86', '89', '90', '112', '113', '124', '161', '162', '165', '166', '176', '177', '184', '185', '186', '198', '207', '217'] |

| Protein name | Number of atom types | Atom types | Proportion of predictions with set size = 1 | Accuracy (posterior probability = 1) | Accuracy (posterior probability > 75%) | Accuracy of candidate assignment set | Median set size | Proportion in-distribution | Residue numbers out of distribution |
|---|---|---|---|---|---|---|---|---|---|
| **CBP_TAZ4** | 9 | ['CA', 'CA(i-1)', 'CB', 'CB(i-1)', 'N(i+1)', 'N(i-1)', 'C', 'C(i-1)', 'N'] | 0.22 | 1.00 | 0.97 | 0.91 | 2.0 | 0.65 | ['3', '6', '7', '10', '11', '12', '13', '16', '18', '19', '20', '21', '22', '24', '25', '27', '28', '29', '30', '31', '32', '35', '36', '39', '40', '41', '42', '45', '46', '47', '48', '49', '51', '53', '54', '55', '56', '58', '59', '60', '62', '63', '64', '65', '66', '67', '68', '69', '71', '72', '73', '75', '78', '79', '80', '82', '83', '104', '105', '128', '129', '131', '135', '136', '141', '142', '144', '149', '152', '156', '158', '167', '168', '169', '176', '177', '181', '183', '184', '185', '196', '197', '200', '203', '204', '205', '220', '222', '249', '250', '265', '271', '273', '277', '281', '282', '293'] |

**Supplementary Table 3** Peaklist of backbone chemical shifts for IAPP used to validate SpinForecast, obtained from BMRB entry 51259. Residue numbers correspond to the sequence in Supplementary Table 1. Missing values are denoted by 999.000.

| Residue number (IAPP) | H (ppm) | N (ppm) | CA (ppm) | CA(i-1) (ppm) | CB (ppm) | CB(i-1) (ppm) | C (ppm) | C(i-1) (ppm) | N(i-1) (ppm) | N(i+1) (ppm) |
|---|---|---|---|---|---|---|---|---|---|---|
| **3N** | 999.000 | 999.000 | 52.868 | 999.000 | 39.402 | 999.000 | 174.970 | 999.000 | 999.000 | 110.957 |
| **4T** | 7.616 | 110.957 | 61.093 | 52.868 | 70.942 | 39.402 | 174.773 | 174.970 | 999.000 | 122.826 |
| **5A** | 8.714 | 122.826 | 54.742 | 61.093 | 18.546 | 70.942 | 179.546 | 174.773 | 110.957 | 109.318 |
| **6T** | 7.934 | 109.318 | 62.802 | 54.742 | 69.365 | 18.546 | 175.208 | 179.546 | 122.826 | 120.517 |
| **7C** | 8.108 | 120.517 | 56.288 | 62.802 | 41.100 | 69.365 | 174.984 | 175.208 | 109.318 | 124.517 |
| **8A** | 8.201 | 124.517 | 53.533 | 56.288 | 18.829 | 41.100 | 178.434 | 174.984 | 120.517 | 113.557 |
| **9T** | 8.132 | 113.557 | 62.876 | 53.533 | 69.457 | 18.829 | 175.064 | 178.434 | 124.517 | 122.737 |
| **10Q** | 8.251 | 122.737 | 56.359 | 62.876 | 29.132 | 69.457 | 176.126 | 175.064 | 113.557 | 122.116 |
| **11R** | 8.314 | 122.116 | 56.590 | 56.359 | 30.418 | 29.132 | 176.570 | 176.126 | 122.737 | 123.013 |
| **12L** | 8.198 | 123.013 | 55.281 | 56.590 | 42.240 | 30.418 | 177.350 | 176.570 | 122.116 | 124.005 |
| **13A** | 8.228 | 124.005 | 52.885 | 55.281 | 18.953 | 42.240 | 177.614 | 177.350 | 123.013 | 116.955 |
| **14N** | 8.247 | 116.955 | 53.364 | 52.885 | 38.534 | 18.953 | 175.061 | 177.614 | 124.005 | 120.120 |
| **15F** | 8.035 | 120.120 | 58.087 | 53.364 | 39.207 | 38.534 | 175.558 | 175.061 | 116.955 | 123.080 |
| **16L** | 8.076 | 123.080 | 55.127 | 58.087 | 42.229 | 39.207 | 176.982 | 175.558 | 120.120 | 121.073 |
| **17V** | 7.984 | 121.073 | 62.518 | 55.127 | 32.637 | 42.229 | 175.944 | 176.982 | 123.080 | 123.237 |
| **18H** | 8.418 | 123.237 | 56.089 | 62.518 | 30.641 | 32.637 | 175.218 | 175.944 | 121.073 | 117.518 |
| **19S** | 8.301 | 117.518 | 58.290 | 56.089 | 63.819 | 30.641 | 174.567 | 175.218 | 123.237 | 117.904 |
| **20S** | 8.498 | 117.904 | 58.492 | 58.290 | 63.729 | 63.819 | 174.209 | 174.567 | 117.518 | 120.188 |
| **21N** | 8.391 | 120.188 | 53.128 | 58.492 | 38.644 | 63.729 | 174.761 | 174.209 | 117.904 | 118.781 |
| **22N** | 8.287 | 118.781 | 53.097 | 53.128 | 38.529 | 38.644 | 175.053 | 174.761 | 120.188 | 120.466 |
| **23F** | 8.253 | 120.466 | 58.252 | 53.097 | 38.856 | 38.529 | 176.368 | 175.053 | 118.781 | 110.271 |
| **24G** | 8.324 | 110.271 | 45.202 | 58.252 | 999.000 | 38.856 | 173.713 | 176.368 | 120.466 | 123.644 |
| **25A** | 8.001 | 123.644 | 52.434 | 45.202 | 19.181 | 999.000 | 177.646 | 173.713 | 110.271 | 120.684 |
| **26I** | 8.202 | 120.684 | 61.068 | 52.434 | 38.481 | 19.181 | 176.433 | 177.646 | 123.644 | 127.079 |
| **27L** | 8.432 | 127.079 | 54.990 | 61.068 | 42.209 | 38.481 | 177.282 | 176.433 | 120.684 | 116.977 |
| **28S** | 8.382 | 116.977 | 58.204 | 54.990 | 63.785 | 42.209 | 174.643 | 177.282 | 127.079 | 117.909 |
| **29S** | 8.443 | 117.909 | 58.404 | 58.204 | 63.728 | 63.785 | 174.727 | 174.643 | 116.977 | 115.207 |
| **30T** | 8.183 | 115.207 | 61.886 | 58.404 | 69.615 | 63.728 | 174.201 | 174.727 | 117.909 | 121.483 |
| **31N** | 8.430 | 121.483 | 53.213 | 61.886 | 38.708 | 69.615 | 175.152 | 174.201 | 115.207 | 120.681 |
| **32V** | 8.202 | 120.681 | 62.629 | 53.213 | 32.437 | 38.708 | 176.726 | 175.152 | 121.483 | 112.576 |
| **33G** | 8.549 | 112.576 | 45.191 | 62.629 | 999.000 | 32.437 | 174.119 | 176.726 | 120.681 | 115.543 |
| **34S** | 8.243 | 115.543 | 58.205 | 45.191 | 63.794 | 999.000 | 174.243 | 174.119 | 112.576 | 120.716 |
| **35N** | 8.544 | 120.716 | 53.246 | 58.205 | 38.795 | 63.794 | 174.995 | 174.243 | 115.543 | 114.035 |
| **36T** | 8.060 | 114.035 | 61.632 | 53.246 | 69.877 | 38.795 | 173.166 | 174.995 | 120.716 | 127.036 |
| **37Y** | 7.777 | 127.036 | 59.048 | 61.632 | 39.339 | 69.877 | 180.329 | 173.166 | 114.035 | 999.000 |

**Supplementary Table 4** Peaklist of backbone chemical shifts for NUPR1 used to validate SpinForecast. Residue numbers correspond to the sequence in Supplementary Table 1. Missing values are denoted by 999.000. Residues numbers followed by prime (such as 2A') correspond to minor population states, for example arising from nearby proline residues adopting *cis* conformations. Assignments are deposited in the BMRB (accession number 53937).

| Residue number (NUPR1) | H (ppm) | N (ppm) | CA (ppm) | CA(i-1) (ppm) | CB (ppm) | CB(i-1) (ppm) | C (ppm) | C(i-1) (ppm) | N(i-1) (ppm) | N(i+1) (ppm) |
|---|---|---|---|---|---|---|---|---|---|---|
| 2A | 8.499 | 126.196 | 52.528 | 55.500 | 19.293 | 33.050 | 177.437 | 175.800 | 999.000 | 113.519 |
| 2A' | 8.507 | 126.270 | 52.500 | 55.450 | 19.350 | 33.125 | 177.615 | 175.780 | 999.000 | 114.281 |
| 3T | 7.996 | 113.519 | 61.433 | 52.528 | 70.038 | 19.293 | 173.716 | 177.437 | 126.196 | 122.903 |
| 3T' | 8.138 | 114.281 | 61.850 | 52.475 | 70.175 | 19.350 | 173.050 | 177.615 | 126.270 | 123.754 |
| 4F | 8.310 | 122.903 | 55.664 | 61.433 | 41.918 | 70.038 | 173.303 | 173.716 | 113.519 | 138.300 |
| 4F' | 8.317 | 123.754 | 55.675 | 61.825 | 41.920 | 70.150 | 999.000 | 173.050 | 114.281 | 999.000 |
| 5P | 999.000 | 138.300 | 999.000 | 55.664 | 999.000 | 41.920 | 999.000 | 173.303 | 122.903 | 999.000 |
| 7A | 8.502 | 124.550 | 52.505 | 62.850 | 19.193 | 32.100 | 178.142 | 176.825 | 999.000 | 113.300 |
| 7A' | 8.510 | 124.735 | 52.500 | 62.875 | 19.225 | 31.925 | 178.080 | 176.560 | 999.000 | 113.284 |
| 8T | 8.160 | 113.300 | 61.597 | 52.505 | 69.922 | 19.193 | 174.468 | 178.142 | 124.550 | 118.111 |
| 8T' | 8.167 | 113.284 | 61.600 | 52.475 | 69.925 | 19.225 | 999.000 | 178.080 | 124.735 | 999.000 |
| 9S | 8.302 | 118.111 | 58.028 | 61.597 | 63.973 | 69.922 | 173.666 | 174.468 | 113.300 | 127.280 |
| 10A | 8.340 | 127.280 | 50.633 | 58.028 | 18.203 | 63.973 | 175.403 | 173.666 | 118.111 | 135.950 |
| 11P | 999.000 | 135.950 | 63.020 | 50.633 | 32.050 | 18.203 | 176.910 | 175.403 | 127.280 | 121.010 |
| 12Q | 8.546 | 121.010 | 55.651 | 63.020 | 29.611 | 32.050 | 175.846 | 176.910 | 135.950 | 123.236 |
| 13Q | 8.488 | 123.236 | 53.540 | 55.651 | 28.909 | 29.611 | 173.494 | 175.846 | 121.010 | 138.850 |
| 14P | 999.000 | 138.850 | 999.000 | 53.540 | 999.000 | 28.909 | 999.000 | 173.494 | 123.236 | 999.000 |
| 16G | 8.472 | 109.745 | 44.434 | 63.100 | 999.000 | 32.225 | 172.161 | 177.175 | 999.000 | 134.400 |
| 16G' | 8.445 | 109.398 | 44.410 | 63.350 | 999.000 | 999.000 | 999.000 | 177.260 | 999.000 | 999.000 |
| 17P | 999.000 | 134.400 | 63.470 | 44.434 | 32.150 | 999.000 | 177.540 | 172.161 | 109.745 | 120.264 |
| 18E | 8.749 | 120.264 | 56.673 | 63.470 | 29.937 | 32.150 | 176.433 | 177.540 | 134.400 | 120.870 |
| 18E' | 8.734 | 120.603 | 56.650 | 63.350 | 30.050 | 32.230 | 176.400 | 177.495 | 999.000 | 121.028 |
| 19D | 8.231 | 120.870 | 54.465 | 56.673 | 41.343 | 29.937 | 176.340 | 176.433 | 120.264 | 121.285 |
| 19D' | 8.275 | 121.028 | 54.425 | 56.680 | 41.390 | 30.050 | 999.000 | 176.400 | 120.603 | 999.000 |
| 20E | 8.334 | 121.285 | 56.767 | 54.465 | 30.462 | 41.343 | 176.339 | 176.340 | 120.870 | 121.596 |
| 21D | 8.472 | 121.596 | 54.435 | 56.767 | 41.156 | 30.462 | 176.585 | 176.339 | 121.285 | 117.412 |
| 22S | 8.346 | 117.412 | 58.725 | 54.435 | 63.761 | 41.156 | 174.895 | 176.585 | 121.596 | 118.145 |
| 23S | 8.444 | 118.145 | 58.838 | 58.725 | 63.769 | 63.761 | 174.664 | 174.895 | 117.412 | 123.684 |
| 24L | 8.140 | 123.684 | 55.289 | 58.838 | 42.344 | 63.769 | 177.064 | 174.664 | 118.145 | 121.578 |
| 25D | 8.376 | 121.578 | 54.280 | 55.289 | 41.391 | 42.344 | 176.579 | 177.064 | 123.684 | 122.055 |
| 26E | 8.514 | 122.055 | 57.216 | 54.280 | 29.948 | 41.391 | 176.971 | 176.579 | 121.578 | 116.200 |
| 27S | 8.398 | 116.200 | 59.547 | 57.216 | 63.666 | 29.948 | 174.925 | 176.971 | 122.055 | 122.332 |
| 28D | 8.316 | 122.332 | 54.770 | 59.547 | 41.000 | 63.666 | 176.799 | 174.925 | 116.200 | 121.515 |
| 29L | 8.030 | 121.515 | 56.447 | 54.770 | 41.971 | 41.000 | 178.025 | 176.799 | 122.332 | 118.980 |
| 30Y | 8.072 | 118.980 | 58.744 | 56.447 | 38.312 | 41.971 | 176.635 | 178.025 | 121.515 | 116.113 |
| 31S | 8.018 | 116.113 | 59.084 | 58.744 | 63.604 | 38.312 | 175.353 | 176.635 | 118.980 | 123.456 |
| 32L | 8.186 | 123.456 | 56.009 | 59.084 | 42.018 | 63.604 | 177.913 | 175.353 | 116.113 | 122.421 |

| Residue number (NUPR1) | H (ppm) | N (ppm) | CA (ppm) | CA(i-1) (ppm) | CB (ppm) | CB(i-1) (ppm) | C (ppm) | C(i-1) (ppm) | N(i-1) (ppm) | N(i+1) (ppm) |
|---|---|---|---|---|---|---|---|---|---|---|
| **33A** | 8.052 | 122.421 | 53.170 | 56.009 | 18.812 | 42.018 | 177.961 | 177.913 | 123.456 | 116.850 |
| **34H** | 7.974 | 116.850 | 56.349 | 53.170 | 30.139 | 18.812 | 175.422 | 177.961 | 122.421 | 116.107 |
| **35S** | 8.048 | 116.107 | 58.789 | 56.349 | 63.793 | 30.139 | 174.470 | 175.422 | 116.850 | 121.948 |
| **36Y** | 8.197 | 121.948 | 58.183 | 58.789 | 38.356 | 63.793 | 176.134 | 174.470 | 116.107 | 123.400 |
| **37L** | 8.150 | 123.400 | 55.343 | 58.183 | 42.074 | 38.356 | 177.883 | 176.134 | 121.948 | 108.660 |
| **38G** | 7.964 | 108.660 | 45.458 | 55.343 | 999.000 | 42.074 | 174.802 | 177.883 | 123.400 | 108.730 |
| **39G** | 8.275 | 108.730 | 45.410 | 45.458 | 999.000 | 999.000 | 175.033 | 174.802 | 108.660 | 108.924 |
| **40G** | 8.418 | 108.924 | 45.437 | 45.410 | 999.000 | 999.000 | 174.867 | 175.033 | 108.730 | 108.730 |
| **41G** | 8.348 | 108.730 | 45.284 | 45.437 | 999.000 | 999.000 | 174.268 | 174.867 | 108.924 | 120.639 |
| **42R** | 8.247 | 120.639 | 56.400 | 45.284 | 30.800 | 999.000 | 176.683 | 174.268 | 108.730 | 122.353 |
| **43K** | 8.461 | 122.353 | 56.658 | 56.400 | 32.900 | 30.800 | 177.170 | 176.683 | 120.639 | 110.003 |
| **44G** | 8.411 | 110.003 | 45.235 | 56.658 | 999.000 | 32.900 | 174.064 | 177.170 | 122.353 | 120.698 |
| **45R** | 8.247 | 120.698 | 56.328 | 45.235 | 30.857 | 999.000 | 176.739 | 174.064 | 110.003 | 115.709 |
| **46T** | 8.304 | 115.709 | 62.121 | 56.328 | 70.052 | 30.857 | 174.634 | 176.739 | 120.698 | 124.094 |
| **47K** | 8.454 | 124.094 | 56.886 | 62.121 | 32.953 | 70.052 | 176.802 | 174.634 | 115.709 | 122.772 |
| **48R** | 8.421 | 122.772 | 56.626 | 56.886 | 30.735 | 32.953 | 176.649 | 176.802 | 124.094 | 122.416 |
| **49E** | 8.454 | 122.416 | 56.633 | 56.626 | 30.306 | 30.735 | 176.479 | 176.649 | 122.772 | 125.111 |
| **50A** | 8.363 | 125.111 | 52.768 | 56.633 | 19.124 | 30.306 | 177.803 | 176.479 | 122.416 | 123.077 |
| **51A** | 8.243 | 123.077 | 52.627 | 52.768 | 19.079 | 19.124 | 177.679 | 177.803 | 125.111 | 122.836 |
| **52A** | 8.220 | 122.836 | 52.689 | 52.627 | 19.141 | 19.079 | 177.717 | 177.679 | 123.077 | 117.647 |
| **53N** | 8.377 | 117.647 | 53.321 | 52.689 | 38.751 | 19.141 | 175.625 | 177.717 | 122.836 | 113.873 |
| **54T** | 8.098 | 113.873 | 62.084 | 53.321 | 69.609 | 38.751 | 174.382 | 175.625 | 117.647 | 120.866 |
| **55N** | 8.450 | 120.866 | 53.305 | 62.084 | 38.769 | 69.609 | 174.724 | 174.382 | 113.873 | 122.407 |
| **56R** | 8.222 | 122.407 | 53.962 | 53.305 | 30.197 | 38.769 | 173.982 | 174.724 | 120.866 | 137.160 |
| **57P** | 999.000 | 137.160 | 62.900 | 53.962 | 32.150 | 30.197 | 176.750 | 173.982 | 122.407 | 117.934 |
| **58S** | 8.572 | 117.934 | 56.444 | 62.900 | 63.312 | 32.150 | 173.066 | 176.750 | 137.160 | 138.425 |
| **58S'** | 8.278 | 116.343 | 55.520 | 63.200 | 64.250 | 32.175 | 999.000 | 176.050 | 999.000 | 999.000 |
| **59P** | 999.000 | 138.425 | 63.650 | 56.444 | 32.050 | 63.312 | 177.580 | 173.066 | 117.934 | 109.404 |
| **60G** | 8.532 | 109.404 | 45.282 | 63.650 | 999.000 | 32.050 | 174.771 | 177.580 | 138.425 | 108.777 |
| **61G** | 8.316 | 108.777 | 45.373 | 45.282 | 999.000 | 999.000 | 174.317 | 174.771 | 109.404 | 119.229 |
| **62H** | 8.270 | 119.229 | 56.428 | 45.373 | 30.651 | 999.000 | 175.579 | 174.317 | 108.777 | 122.050 |
| **63E** | 8.538 | 122.050 | 56.906 | 56.428 | 29.990 | 30.651 | 176.519 | 175.579 | 119.229 | 122.554 |
| **64R** | 8.406 | 122.554 | 56.238 | 56.906 | 30.676 | 29.990 | 176.266 | 176.519 | 122.050 | 122.937 |
| **65K** | 8.353 | 122.937 | 56.353 | 56.238 | 32.971 | 30.676 | 176.437 | 176.266 | 122.554 | 124.019 |
| **66L** | 8.303 | 124.019 | 55.223 | 56.353 | 42.247 | 32.971 | 177.274 | 176.437 | 122.937 | 122.179 |
| **67V** | 8.227 | 122.179 | 62.503 | 55.223 | 32.782 | 42.247 | 176.427 | 177.274 | 124.019 | 118.942 |
| **68T** | 8.249 | 118.942 | 62.083 | 62.503 | 69.853 | 32.782 | 174.436 | 176.427 | 122.179 | 124.108 |
| **69K** | 8.363 | 124.108 | 56.621 | 62.083 | 33.015 | 69.853 | 176.669 | 174.436 | 118.942 | 123.578 |
| **70L** | 8.309 | 123.578 | 55.435 | 56.621 | 42.232 | 33.015 | 177.658 | 176.669 | 124.108 | 121.305 |
| **71Q** | 8.455 | 121.305 | 56.132 | 55.435 | 29.360 | 42.232 | 176.177 | 177.658 | 123.578 | 120.116 |
| **72N** | 8.548 | 120.116 | 53.801 | 56.132 | 38.811 | 29.360 | 175.775 | 176.177 | 121.305 | 116.344 |

| Residue number (NUPR1) | H (ppm) | N (ppm) | CA (ppm) | CA(i-1) (ppm) | CB (ppm) | CB(i-1) (ppm) | C (ppm) | C(i-1) (ppm) | N(i-1) (ppm) | N(i+1) (ppm) |
|---|---|---|---|---|---|---|---|---|---|---|
| **73S** | 8.370 | 116.344 | 59.149 | 53.801 | 63.601 | 38.811 | 174.970 | 175.775 | 120.116 | 122.638 |
| **74E** | 8.386 | 122.638 | 57.031 | 59.149 | 30.039 | 63.601 | 176.886 | 174.970 | 116.344 | 121.759 |
| **75R** | 8.248 | 121.759 | 56.746 | 57.031 | 30.466 | 30.039 | 176.744 | 176.886 | 122.638 | 122.047 |
| **76K** | 8.252 | 122.047 | 56.606 | 56.746 | 32.934 | 30.466 | 176.867 | 176.744 | 121.759 | 122.538 |
| **77K** | 8.280 | 122.538 | 56.600 | 56.606 | 32.987 | 32.934 | 176.825 | 176.867 | 122.047 | 122.493 |
| **78R** | 8.399 | 122.493 | 56.550 | 56.600 | 30.760 | 32.987 | 176.983 | 176.825 | 122.538 | 110.357 |
| **79G** | 8.445 | 110.357 | 45.128 | 56.550 | 999.000 | 30.760 | 173.725 | 176.983 | 122.493 | 123.768 |
| **80A** | 8.168 | 123.768 | 52.453 | 45.128 | 19.520 | 999.000 | 177.749 | 173.725 | 110.357 | 120.913 |
| **81R** | 8.377 | 120.913 | 56.228 | 52.453 | 30.734 | 19.520 | 175.515 | 177.749 | 123.768 | 127.572 |
| **82R** | 8.039 | 127.572 | 57.497 | 56.228 | 31.436 | 30.734 | 173.994 | 175.515 | 120.913 | 999.000 |

**Supplementary Table 5** Peaklist of backbone chemical shifts for JPT2 used to validate SpinForecast. Missing values are denoted by 999.000. Residue numbers correspond to the sequence in Supplementary Table 1. Assignments are deposited in the BMRB (accession number 53938).

| Residue number (JPT2) | H (ppm) | N (ppm) | CA (ppm) | CA(i-1) (ppm) | CB (ppm) | CB(i-1) (ppm) | C (ppm) | C(i-1) (ppm) | N(i-1) (ppm) | N(i+1) (ppm) |
|---|---|---|---|---|---|---|---|---|---|---|
| **2F** | 8.493 | 121.627 | 57.592 | 999.000 | 39.489 | 999.000 | 175.178 | 999.000 | 999.000 | 123.094 |
| **3Q** | 8.313 | 123.094 | 55.046 | 57.592 | 29.556 | 39.489 | 175.053 | 175.178 | 121.627 | 123.902 |
| **4V** | 8.357 | 123.902 | 59.902 | 55.046 | 32.111 | 29.556 | 174.550 | 175.053 | 123.094 | 139.698 |
| **5P** | 999.000 | 139.698 | 62.995 | 59.902 | 32.230 | 32.111 | 176.723 | 174.550 | 123.902 | 121.393 |
| **6D** | 8.599 | 121.393 | 54.582 | 62.995 | 40.975 | 32.230 | 176.743 | 176.723 | 139.698 | 116.266 |
| **7S** | 8.411 | 116.266 | 58.788 | 54.582 | 63.686 | 40.975 | 175.026 | 176.743 | 121.393 | 122.474 |
| **8E** | 8.519 | 122.474 | 56.938 | 58.788 | 29.908 | 63.686 | 177.390 | 175.026 | 116.266 | 109.367 |
| **9G** | 8.454 | 109.367 | 45.703 | 56.938 | 999.000 | 29.908 | 175.056 | 177.390 | 122.474 | 108.814 |
| **10G** | 8.327 | 108.814 | 45.364 | 45.703 | 999.000 | 999.000 | 174.540 | 175.056 | 109.367 | 120.642 |
| **11R** | 8.207 | 120.642 | 56.343 | 45.364 | 30.648 | 999.000 | 176.616 | 174.540 | 108.814 | 124.904 |
| **12A** | 8.485 | 124.904 | 53.149 | 56.343 | 18.789 | 30.648 | 178.568 | 176.616 | 120.642 | 108.315 |
| **13G** | 8.312 | 108.315 | 45.350 | 53.149 | 999.000 | 18.789 | 174.481 | 178.568 | 124.904 | 115.797 |
| **14S** | 8.192 | 115.797 | 58.595 | 45.350 | 63.891 | 999.000 | 174.894 | 174.481 | 108.315 | 123.089 |
| **15R** | 8.476 | 123.089 | 56.700 | 58.595 | 30.324 | 63.891 | 176.724 | 174.894 | 115.797 | 121.627 |
| **16K** | 8.313 | 121.627 | 57.300 | 56.700 | 32.468 | 30.324 | 176.838 | 176.724 | 123.089 | 121.180 |
| **17W** | 8.067 | 121.180 | 57.849 | 57.300 | 29.352 | 32.468 | 176.619 | 176.838 | 121.627 | 121.491 |
| **18Q** | 8.147 | 121.491 | 56.423 | 57.849 | 29.543 | 29.352 | 175.916 | 176.619 | 121.180 | 122.306 |
| **19L** | 8.137 | 122.306 | 55.489 | 56.423 | 42.177 | 29.543 | 177.689 | 175.916 | 121.491 | 122.664 |
| **20L** | 8.253 | 122.664 | 55.477 | 55.489 | 42.070 | 42.177 | 177.755 | 177.689 | 122.306 | 113.440 |
| **21T** | 8.057 | 113.440 | 62.083 | 55.477 | 69.869 | 42.070 | 175.248 | 177.755 | 122.664 | 111.050 |
| **22G** | 8.390 | 111.050 | 45.410 | 62.083 | 999.000 | 69.869 | 174.376 | 175.248 | 113.440 | 115.927 |
| **23S** | 8.289 | 115.927 | 58.474 | 45.410 | 63.798 | 999.000 | 174.748 | 174.376 | 111.050 | 123.914 |
| **24L** | 8.354 | 123.914 | 55.239 | 58.474 | 42.262 | 63.798 | 177.355 | 174.748 | 115.927 | 124.376 |
| **25A** | 8.288 | 124.376 | 52.608 | 55.239 | 19.150 | 42.262 | 177.914 | 177.355 | 123.914 | 114.787 |
| **26S** | 8.318 | 114.787 | 58.337 | 52.608 | 63.701 | 19.150 | 174.852 | 177.914 | 124.376 | 115.311 |
| **27T** | 8.171 | 115.311 | 61.566 | 58.337 | 69.700 | 63.701 | 174.331 | 174.852 | 114.787 | 119.534 |
| **28S** | 8.349 | 119.534 | 56.476 | 61.566 | 63.293 | 69.700 | 172.839 | 174.331 | 115.311 | 138.348 |
| **29P** | 999.000 | 138.348 | 63.581 | 56.476 | 31.874 | 63.293 | 177.102 | 172.839 | 119.534 | 115.702 |
| **30S** | 8.406 | 115.702 | 58.520 | 63.581 | 63.588 | 31.874 | 174.889 | 177.102 | 138.348 | 124.306 |
| **31L** | 8.319 | 124.306 | 55.407 | 58.520 | 42.089 | 63.588 | 177.632 | 174.889 | 115.702 | 121.945 |
| **32L** | 8.197 | 121.945 | 55.037 | 55.407 | 42.063 | 42.089 | 177.553 | 177.632 | 124.306 | 115.930 |
| **33S** | 8.242 | 115.930 | 58.634 | 55.037 | 63.674 | 42.063 | 175.249 | 177.553 | 121.945 | 110.910 |
| **34G** | 8.483 | 110.910 | 45.463 | 58.634 | 999.000 | 63.674 | 174.097 | 175.249 | 115.930 | 119.303 |
| **35Q** | 8.278 | 119.303 | 55.515 | 45.463 | 29.822 | 999.000 | 176.083 | 174.097 | 110.910 | 110.252 |
| **36G** | 8.332 | 110.252 | 44.494 | 55.515 | 999.000 | 29.822 | 171.928 | 176.083 | 119.303 | 133.849 |
| **37P** | 999.000 | 133.849 | 63.299 | 44.494 | 31.799 | 999.000 | 176.548 | 171.928 | 110.252 | 120.999.000 |
| **38W** | 8.163 | 120.999.000 | 57.234 | 63.299 | 29.289 | 31.799 | 175.459 | 176.548 | 133.849 | 127.595 |

| Residue number (JPT2) | H (ppm) | N (ppm) | CA (ppm) | CA(i-1) (ppm) | CB (ppm) | CB(i-1) (ppm) | C (ppm) | C(i-1) (ppm) | N(i-1) (ppm) | N(i+1) (ppm) |
|---|---|---|---|---|---|---|---|---|---|---|
| **39A** | 7.816 | 127.595 | 50.490 | 57.234 | 18.572 | 29.289 | 174.512 | 175.459 | 120.999.000 | 134.487 |
| **40P** | 999.000 | 134.487 | 63.094 | 50.490 | 32.004 | 18.572 | 177.057 | 174.512 | 127.595 | 121.576 |
| **41L** | 8.343 | 121.576 | 55.318 | 63.094 | 42.005 | 32.004 | 177.566 | 177.057 | 134.487 | 121.382 |
| **42Q** | 8.365 | 121.382 | 55.671 | 55.318 | 29.417 | 42.005 | 175.904 | 177.566 | 121.576 | 122.703 |
| **43R** | 8.407 | 122.703 | 56.067 | 55.671 | 30.903 | 29.417 | 175.852 | 175.904 | 121.382 | 125.207 |
| **44A** | 8.410 | 125.207 | 52.331 | 56.067 | 19.089 | 30.903 | 177.537 | 175.852 | 122.703 | 120.365 |
| **45M** | 8.413 | 120.365 | 55.216 | 52.331 | 33.061 | 19.089 | 175.933 | 177.537 | 125.207 | 124.435 |
| **46K** | 8.424 | 124.435 | 54.092 | 55.216 | 32.506 | 33.061 | 173.912 | 175.933 | 120.365 | 138.820 |
| **47P** | 999.000 | 138.820 | 61.284 | 54.092 | 30.669 | 32.506 | 999.000 | 173.912 | 124.435 | 999.000 |
| **49G** | 8.636 | 109.521 | 45.206 | 999.000 | 999.000 | 999.000 | 174.844 | 999.000 | 999.000 | 108.825 |
| **50G** | 8.361 | 108.825 | 45.800 | 45.206 | 999.000 | 999.000 | 174.353 | 174.844 | 109.521 | 120.768 |
| **51E** | 8.618 | 120.768 | 56.785 | 45.800 | 30.062 | 999.000 | 176.877 | 174.353 | 108.825 | 116.904 |
| **52S** | 8.549 | 116.904 | 58.462 | 56.785 | 63.673 | 30.062 | 174.670 | 176.877 | 120.768 | 117.770 |
| **53S** | 8.410 | 117.770 | 58.722 | 58.462 | 63.779 | 63.673 | 174.211 | 174.670 | 116.904 | 120.742 |
| **54N** | 8.488 | 120.742 | 53.258 | 58.722 | 38.633 | 63.779 | 175.151 | 174.211 | 117.770 | 122.176 |
| **55L** | 8.204 | 122.176 | 55.363 | 53.258 | 41.950 | 38.633 | 177.234 | 175.151 | 120.742 | 119.940 |
| **56F** | 8.265 | 119.940 | 57.663 | 55.363 | 39.334 | 41.950 | 176.225 | 177.234 | 122.176 | 110.404 |
| **57G** | 8.296 | 110.404 | 45.040 | 57.663 | 999.000 | 39.334 | 173.672 | 176.225 | 119.940 | 117.049 |
| **58S** | 8.297 | 117.049 | 56.538 | 45.040 | 63.339 | 999.000 | 172.955 | 173.672 | 110.404 | 137.987 |
| **59P** | 999.000 | 137.987 | 63.569 | 56.538 | 32.014 | 63.339 | 177.148 | 172.955 | 117.049 | 120.740 |
| **60E** | 8.597 | 120.740 | 56.797 | 63.569 | 30.052 | 32.014 | 176.625 | 177.148 | 137.987 | 122.082 |
| **61E** | 8.379 | 122.082 | 56.353 | 56.797 | 30.447 | 30.052 | 176.049 | 176.625 | 120.740 | 125.484 |
| **62A** | 8.399 | 125.484 | 52.419 | 56.353 | 19.152 | 30.447 | 177.656 | 176.049 | 122.082 | 116.204 |
| **63T** | 8.302 | 116.204 | 59.788 | 52.419 | 69.578 | 19.152 | 173.084 | 177.656 | 125.484 | 139.107 |
| **64P** | 999.000 | 139.107 | 63.475 | 59.788 | 32.001 | 69.578 | 177.185 | 173.084 | 116.204 | 115.874 |
| **65S** | 8.537 | 115.874 | 58.595 | 63.475 | 63.729 | 32.001 | 174.669 | 177.185 | 139.107 | 117.838 |
| **66S** | 8.352 | 117.838 | 58.262 | 58.595 | 63.793 | 63.729 | 174.058 | 174.669 | 115.874 | 123.759 |
| **67R** | 8.334 | 123.759 | 54.105 | 58.262 | 30.175 | 63.793 | 174.155 | 174.058 | 117.838 | 136.959 |
| **68P** | 999.000 | 136.959 | 63.126 | 54.105 | 32.133 | 30.175 | 176.671 | 174.155 | 123.759 | 119.233 |
| **69N** | 8.642 | 119.233 | 53.171 | 63.126 | 38.608 | 32.133 | 175.385 | 176.671 | 136.959 | 121.942 |
| **70R** | 8.437 | 121.942 | 56.285 | 53.171 | 30.640 | 38.608 | 176.294 | 175.385 | 119.233 | 121.388 |
| **71M** | 8.468 | 121.388 | 55.427 | 56.285 | 32.658 | 30.640 | 176.127 | 176.294 | 121.942 | 125.403 |
| **72A** | 8.391 | 125.403 | 52.692 | 55.427 | 19.150 | 32.658 | 177.665 | 176.127 | 121.388 | 114.769 |
| **73S** | 8.342 | 114.769 | 58.296 | 52.692 | 63.717 | 19.150 | 174.226 | 177.665 | 125.403 | 120.724 |
| **74N** | 8.473 | 120.724 | 53.110 | 58.296 | 38.616 | 63.717 | 174.979 | 174.226 | 114.769 | 120.275 |
| **75I** | 7.966 | 120.275 | 61.350 | 53.110 | 38.404 | 38.616 | 175.891 | 174.979 | 120.724 | 123.471 |
| **76F** | 8.340 | 123.471 | 57.433 | 61.350 | 39.630 | 38.404 | 175.829 | 175.891 | 120.275 | 110.513 |
| **77G** | 8.228 | 110.513 | 44.579 | 57.433 | 999.000 | 39.630 | 171.521 | 175.829 | 123.471 | 134.232 |
| **78P** | 999.000 | 134.232 | 63.233 | 44.579 | 32.237 | 999.000 | 175.850 | 171.521 | 110.513 | 114.664 |
| **79T** | 8.413 | 114.664 | 61.941 | 63.233 | 69.733 | 32.237 | 174.430 | 175.850 | 134.232 | 123.396 |

| Residue number (JPT2) | H (ppm) | N (ppm) | CA (ppm) | CA(i-1) (ppm) | CB (ppm) | CB(i-1) (ppm) | C (ppm) | C(i-1) (ppm) | N(i-1) (ppm) | N(i+1) (ppm) |
|---|---|---|---|---|---|---|---|---|---|---|
| **80E** | 8.429 | 123.396 | 56.051 | 61.941 | 30.557 | 69.733 | 176.024 | 174.430 | 114.664 | 124.109 |
| **81E** | 8.586 | 124.109 | 54.275 | 56.051 | 29.608 | 30.557 | 174.605 | 176.024 | 123.396 | 137.685 |
| **82P** | 999.000 | 137.685 | 63.350 | 54.275 | 32.067 | 29.608 | 177.006 | 174.605 | 124.109 | 120.280 |
| **83Q** | 8.689 | 120.280 | 55.675 | 63.350 | 29.573 | 32.067 | 175.753 | 177.006 | 137.685 | 119.819 |
| **84N** | 8.557 | 119.819 | 53.094 | 55.675 | 38.747 | 29.573 | 174.609 | 175.753 | 120.280 | 123.161 |
| **85I** | 8.148 | 123.161 | 58.805 | 53.094 | 38.280 | 38.747 | 174.441 | 174.609 | 119.819 | 140.231 |
| **86P** | 999.000 | 140.231 | 63.040 | 58.805 | 32.159 | 38.280 | 176.705 | 174.441 | 123.161 | 122.291 |
| **87K** | 8.518 | 122.291 | 56.178 | 63.040 | 33.096 | 32.159 | 176.679 | 176.705 | 140.231 | 123.363 |
| **88R** | 8.554 | 123.363 | 55.943 | 56.178 | 30.933 | 33.096 | 176.340 | 176.679 | 122.291 | 116.475 |
| **89T** | 8.386 | 116.475 | 61.591 | 55.943 | 69.906 | 30.933 | 173.934 | 176.340 | 123.363 | 122.441 |
| **90N** | 8.668 | 122.441 | 51.405 | 61.591 | 38.452 | 69.906 | 172.514 | 173.934 | 116.475 | 137.976 |
| **91P** | 999.000 | 137.976 | 61.313 | 51.405 | 30.676 | 38.452 | 999.000 | 172.514 | 122.441 | 999.000 |
| **94G** | 8.331 | 108.674 | 45.100 | 999.000 | 999.000 | 999.000 | 174.287 | 999.000 | 999.000 | 121.138 |
| **95K** | 8.492 | 121.138 | 56.623 | 45.100 | 32.871 | 999.000 | 177.369 | 174.287 | 108.674 | 110.658 |
| **96G** | 8.645 | 110.658 | 45.190 | 56.623 | 999.000 | 32.871 | 174.244 | 177.369 | 121.138 | 115.727 |
| **97S** | 8.389 | 115.727 | 58.541 | 45.190 | 63.981 | 999.000 | 175.163 | 174.244 | 110.658 | 111.255 |
| **98G** | 8.613 | 111.255 | 45.274 | 58.541 | 999.000 | 63.981 | 173.813 | 175.163 | 115.727 | 119.989 |
| **99I** | 7.932 | 119.989 | 61.129 | 45.274 | 38.485 | 999.000 | 175.970 | 173.813 | 111.255 | 124.711 |
| **100F** | 8.465 | 124.711 | 57.432 | 61.129 | 39.627 | 38.485 | 175.063 | 175.970 | 119.989 | 123.033 |
| **101D** | 8.314 | 123.033 | 53.888 | 57.432 | 41.131 | 39.627 | 176.073 | 175.063 | 124.711 | 122.498 |
| **102E** | 8.573 | 122.498 | 56.726 | 53.888 | 30.006 | 41.131 | 176.572 | 176.073 | 123.033 | 116.877 |
| **103S** | 8.520 | 116.877 | 58.832 | 56.726 | 63.789 | 30.006 | 174.429 | 176.572 | 122.498 | 119.065 |
| **104T** | 8.151 | 119.065 | 60.205 | 58.832 | 69.645 | 63.789 | 172.733 | 174.429 | 116.877 | 139.409 |
| **105P** | 999.000 | 139.409 | 63.169 | 60.205 | 32.153 | 69.645 | 177.068 | 172.733 | 119.065 | 121.406 |
| **106V** | 8.401 | 121.406 | 62.759 | 63.169 | 32.596 | 32.153 | 176.500 | 177.068 | 139.409 | 124.357 |
| **107Q** | 8.632 | 124.357 | 55.884 | 62.759 | 29.472 | 32.596 | 176.225 | 176.500 | 121.406 | 116.248 |
| **108T** | 8.331 | 116.248 | 62.048 | 55.884 | 69.762 | 29.472 | 174.562 | 176.225 | 124.357 | 123.215 |
| **109R** | 8.440 | 123.215 | 56.338 | 62.048 | 30.578 | 69.762 | 176.193 | 174.562 | 116.248 | 121.596 |
| **110Q** | 8.494 | 121.596 | 56.088 | 56.338 | 29.367 | 30.578 | 175.524 | 176.193 | 123.215 | 120.956 |
| **111H** | 8.425 | 120.956 | 56.329 | 56.088 | 30.759 | 29.367 | 175.186 | 175.524 | 121.596 | 123.349 |
| **112L** | 8.202 | 123.349 | 54.969 | 56.329 | 42.415 | 30.759 | 176.663 | 175.186 | 120.956 | 120.354 |
| **113N** | 8.598 | 120.354 | 51.336 | 54.969 | 38.559 | 42.415 | 172.345 | 176.663 | 123.349 | 137.905 |
| **114P** | 999.000 | 137.905 | 61.358 | 51.336 | 30.771 | 38.559 | 999.000 | 172.345 | 120.354 | 999.000 |
| **116G** | 8.647 | 109.744 | 45.140 | 999.000 | 999.000 | 999.000 | 174.768 | 999.000 | 999.000 | 108.653 |
| **117G** | 8.293 | 108.653 | 999.000 | 45.140 | 999.000 | 999.000 | 173.963 | 174.768 | 109.744 | 120.992 |
| **118K** | 8.414 | 120.992 | 56.230 | 999.000 | 33.220 | 999.000 | 177.129 | 173.963 | 108.653 | 116.023 |
| **119T** | 8.419 | 116.023 | 61.970 | 56.230 | 69.703 | 33.220 | 174.603 | 177.129 | 120.992 | 117.529 |
| **120S** | 8.380 | 117.529 | 58.231 | 61.970 | 63.940 | 69.703 | 174.187 | 174.603 | 116.023 | 122.891 |
| **121D** | 8.423 | 122.891 | 54.329 | 58.231 | 41.097 | 63.940 | 176.356 | 174.187 | 117.529 | 120.025 |
| **122I** | 8.052 | 120.025 | 61.496 | 54.329 | 38.543 | 41.097 | 176.218 | 176.356 | 122.891 | 123.859 |
| **123F** | 8.372 | 123.859 | 57.993 | 61.496 | 39.345 | 38.543 | 176.311 | 176.218 | 120.025 | 110.752 |

| Residue number (JPT2) | H (ppm) | N (ppm) | CA (ppm) | CA(i-1) (ppm) | CB (ppm) | CB(i-1) (ppm) | C (ppm) | C(i-1) (ppm) | N(i-1) (ppm) | N(i+1) (ppm) |
|---|---|---|---|---|---|---|---|---|---|---|
| **124G** | 8.291 | 110.752 | 45.030 | 57.993 | 999.000 | 39.345 | 173.685 | 176.311 | 123.859 | 117.174 |
| **125S** | 8.253 | 117.174 | 56.470 | 45.030 | 63.199 | 999.000 | 172.571 | 173.685 | 110.752 | 137.948 |
| **126P** | 999.000 | 137.948 | 63.250 | 56.470 | 31.849 | 63.199 | 177.024 | 172.571 | 117.174 | 121.201 |
| **127V** | 8.402 | 121.201 | 62.526 | 63.250 | 32.617 | 31.849 | 176.589 | 177.024 | 137.948 | 118.554 |
| **128T** | 8.329 | 118.554 | 61.531 | 62.526 | 70.001 | 32.617 | 174.404 | 176.589 | 121.201 | 126.702 |
| **129A** | 8.536 | 126.702 | 52.903 | 61.531 | 19.119 | 70.001 | 178.182 | 174.404 | 118.554 | 113.176 |
| **130T** | 8.227 | 113.176 | 62.253 | 52.903 | 69.626 | 19.119 | 174.914 | 178.182 | 126.702 | 118.556 |
| **131S** | 8.333 | 118.556 | 58.687 | 62.253 | 63.612 | 69.626 | 174.691 | 174.914 | 113.176 | 123.237 |
| **132R** | 8.432 | 123.237 | 56.258 | 58.687 | 30.594 | 63.612 | 176.255 | 174.691 | 118.556 | 122.826 |
| **133L** | 8.170 | 122.826 | 54.937 | 56.258 | 42.206 | 30.594 | 176.892 | 176.255 | 123.237 | 124.631 |
| **134A** | 8.240 | 124.631 | 52.264 | 54.937 | 19.292 | 42.206 | 177.058 | 176.892 | 122.826 | 120.084 |
| **135H** | 8.309 | 120.084 | 54.183 | 52.264 | 30.470 | 19.292 | 173.699 | 177.058 | 124.631 | 137.008 |
| **136P** | 999.000 | 137.008 | 63.451 | 54.183 | 32.153 | 30.470 | 176.751 | 173.699 | 120.084 | 119.153 |
| **137N** | 8.879 | 119.153 | 53.062 | 63.451 | 38.880 | 32.153 | 174.837 | 176.751 | 137.008 | 123.162 |
| **138K** | 8.294 | 123.162 | 54.156 | 53.062 | 32.627 | 38.880 | 174.288 | 174.837 | 119.153 | 137.581 |
| **139P** | 999.000 | 137.581 | 63.050 | 54.156 | 32.170 | 32.627 | 176.966 | 174.288 | 123.162 | 121.868 |
| **140K** | 8.590 | 121.868 | 56.554 | 63.050 | 32.811 | 32.170 | 176.460 | 176.966 | 137.581 | 120.914 |
| **141D** | 8.372 | 120.914 | 54.332 | 56.554 | 40.923 | 32.811 | 175.829 | 176.460 | 121.868 | 119.991 |
| **142H** | 8.258 | 119.991 | 56.547 | 54.332 | 30.845 | 40.923 | 174.920 | 175.829 | 120.914 | 122.540 |
| **143V** | 7.956 | 122.540 | 62.226 | 56.547 | 32.784 | 30.845 | 175.395 | 174.920 | 119.991 | 124.592 |
| **144F** | 8.445 | 124.592 | 57.584 | 62.226 | 39.529 | 32.784 | 175.389 | 175.395 | 122.540 | 124.934 |
| **145L** | 8.328 | 124.934 | 54.843 | 57.584 | 42.562 | 39.529 | 176.729 | 175.389 | 124.592 | 121.018 |
| **146C** | 8.491 | 121.018 | 58.405 | 54.843 | 28.033 | 42.562 | 174.564 | 176.729 | 124.934 | 124.012 |
| **147E** | 8.706 | 124.012 | 57.037 | 58.405 | 30.097 | 28.033 | 176.972 | 174.564 | 121.018 | 110.795 |
| **148G** | 8.602 | 110.795 | 45.126 | 57.037 | 999.000 | 30.097 | 173.833 | 176.972 | 124.012 | 120.336 |
| **149E** | 8.213 | 120.336 | 55.902 | 45.126 | 30.782 | 999.000 | 176.275 | 173.833 | 110.795 | 124.211 |
| **150E** | 8.640 | 124.211 | 54.427 | 55.902 | 29.500 | 30.782 | 174.473 | 176.275 | 120.336 | 137.690 |
| **151P** | 999.000 | 137.690 | 63.027 | 54.427 | 31.952 | 29.500 | 176.957 | 174.473 | 124.211 | 122.285 |
| **152K** | 8.625 | 122.285 | 56.129 | 63.027 | 32.972 | 31.952 | 176.866 | 176.957 | 137.690 | 117.189 |
| **153S** | 8.465 | 117.189 | 58.235 | 56.129 | 63.916 | 32.972 | 174.221 | 176.866 | 122.285 | 122.836 |
| **154D** | 8.540 | 122.836 | 54.203 | 58.235 | 40.915 | 63.916 | 176.395 | 174.221 | 117.189 | 122.811 |
| **155L** | 8.260 | 122.811 | 55.622 | 54.203 | 41.942 | 40.915 | 177.807 | 176.395 | 122.836 | 121.656 |
| **156K** | 8.337 | 121.656 | 56.532 | 55.622 | 32.616 | 41.942 | 176.638 | 177.807 | 122.811 | 124.582 |
| **157A** | 8.179 | 124.582 | 52.442 | 56.532 | 19.103 | 32.616 | 177.632 | 176.638 | 121.656 | 123.522 |
| **158A** | 8.307 | 123.522 | 52.489 | 52.442 | 19.105 | 19.103 | 177.812 | 177.632 | 124.582 | 120.403 |
| **159R** | 8.349 | 120.403 | 55.943 | 52.489 | 30.912 | 19.105 | 176.299 | 177.812 | 123.522 | 117.940 |
| **160S** | 8.437 | 117.940 | 58.079 | 55.943 | 63.621 | 30.912 | 173.941 | 176.299 | 120.403 | 124.805 |
| **161I** | 8.354 | 124.805 | 58.709 | 58.079 | 38.609 | 63.621 | 174.521 | 173.941 | 117.940 | 140.661 |
| **162P** | 999.000 | 140.661 | 63.014 | 58.709 | 32.158 | 38.609 | 176.514 | 174.521 | 124.805 | 124.915 |
| **163A** | 8.568 | 124.915 | 52.684 | 63.014 | 19.079 | 32.158 | 178.490 | 176.514 | 140.661 | 109.044 |
| **164G** | 8.534 | 109.044 | 44.940 | 52.684 | 999.000 | 19.079 | 173.639 | 178.490 | 124.915 | 123.581 |

| Residue number (JPT2) | H (ppm) | N (ppm) | CA (ppm) | CA(i-1) (ppm) | CB (ppm) | CB(i-1) (ppm) | C (ppm) | C(i-1) (ppm) | N(i-1) (ppm) | N(i+1) (ppm) |
|---|---|---|---|---|---|---|---|---|---|---|
| **165A** | 8.197 | 123.581 | 52.038 | 44.940 | 19.517 | 999.000 | 177.627 | 173.639 | 109.044 | 122.092 |
| **166E** | 8.592 | 122.092 | 54.449 | 52.038 | 29.435 | 19.517 | 174.862 | 177.627 | 123.581 | 137.298 |
| **167P** | 999.000 | 137.298 | 63.543 | 54.449 | 32.078 | 29.435 | 177.618 | 174.862 | 122.092 | 109.187 |
| **168G** | 8.563 | 109.187 | 45.085 | 63.543 | 999.000 | 32.078 | 174.190 | 177.618 | 137.298 | 120.789 |
| **169E** | 8.340 | 120.789 | 56.547 | 45.085 | 30.214 | 999.000 | 176.960 | 174.190 | 109.187 | 122.852 |
| **170K** | 8.623 | 122.852 | 56.760 | 56.547 | 32.694 | 30.214 | 177.465 | 176.960 | 120.789 | 110.284 |
| **171G** | 8.587 | 110.284 | 45.330 | 56.760 | 999.000 | 32.694 | 174.359 | 177.465 | 122.852 | 115.771 |
| **172S** | 8.255 | 115.771 | 58.399 | 45.330 | 63.900 | 999.000 | 174.441 | 174.359 | 110.284 | 125.997 |
| **173A** | 8.434 | 125.997 | 52.562 | 58.399 | 19.058 | 63.900 | 177.739 | 174.441 | 115.771 | 120.734 |
| **174R** | 8.343 | 120.734 | 55.981 | 52.562 | 30.740 | 19.058 | 176.308 | 177.739 | 125.997 | 123.908 |
| **175K** | 8.491 | 123.908 | 56.386 | 55.981 | 33.046 | 30.740 | 176.190 | 176.308 | 120.734 | 126.129 |
| **176A** | 8.500 | 126.129 | 52.356 | 56.386 | 19.511 | 33.046 | 177.846 | 176.190 | 123.908 | 108.828 |
| **177G** | 8.338 | 108.828 | 44.347 | 52.356 | 999.000 | 19.511 | 171.521 | 177.846 | 126.129 | 134.310 |
| **178P** | 999.000 | 134.310 | 62.900 | 44.347 | 32.200 | 999.000 | 176.845 | 171.521 | 108.828 | 124.809 |
| **179A** | 8.543 | 124.809 | 52.375 | 62.900 | 19.081 | 32.200 | 177.859 | 176.845 | 134.310 | 121.283 |
| **180K** | 8.435 | 121.283 | 56.214 | 52.375 | 33.060 | 19.081 | 176.660 | 177.859 | 124.809 | 122.694 |
| **181E** | 8.592 | 122.694 | 56.536 | 56.214 | 30.101 | 33.060 | 176.298 | 176.660 | 121.283 | 121.591 |
| **182Q** | 8.529 | 121.591 | 55.501 | 56.536 | 29.643 | 30.101 | 175.731 | 176.298 | 122.694 | 124.641 |
| **183E** | 8.593 | 124.641 | 54.474 | 55.501 | 29.601 | 29.643 | 174.369 | 175.731 | 121.591 | 137.047 |
| **184P** | 999.000 | 137.047 | 62.883 | 54.474 | 32.163 | 29.601 | 177.031 | 174.369 | 124.641 | 123.013 |
| **185M** | 8.602 | 123.013 | 53.911 | 62.883 | 30.019 | 32.163 | 174.288 | 177.031 | 137.047 | 137.290 |
| **186P** | 999.000 | 137.290 | 63.050 | 53.911 | 32.179 | 30.019 | 176.944 | 174.288 | 123.013 | 115.801 |
| **187T** | 8.472 | 115.801 | 61.841 | 63.050 | 69.962 | 32.179 | 174.726 | 176.944 | 137.290 | 122.803 |
| **188V** | 8.353 | 122.803 | 62.148 | 61.841 | 32.907 | 69.962 | 175.731 | 174.726 | 115.801 | 124.644 |
| **189D** | 8.551 | 124.644 | 54.291 | 62.148 | 41.239 | 32.907 | 176.200 | 175.731 | 122.803 | 116.921 |
| **190S** | 8.345 | 116.921 | 58.417 | 54.291 | 63.574 | 41.239 | 174.358 | 176.200 | 124.644 | 121.367 |
| **191H** | 8.486 | 121.367 | 56.164 | 58.417 | 30.300 | 63.574 | 174.934 | 174.358 | 116.921 | 123.567 |
| **192E** | 8.280 | 123.567 | 54.267 | 56.164 | 29.693 | 30.300 | 174.354 | 174.934 | 121.367 | 137.748 |
| **193P** | 999.000 | 137.748 | 63.309 | 54.267 | 32.093 | 29.693 | 176.956 | 174.354 | 123.567 | 121.661 |
| **194R** | 8.629 | 121.661 | 56.161 | 63.309 | 30.528 | 32.093 | 176.350 | 176.956 | 137.748 | 123.448 |
| **195L** | 8.412 | 123.448 | 54.896 | 56.161 | 42.504 | 30.528 | 177.539 | 176.350 | 121.661 | 109.778 |
| **196G** | 8.277 | 109.778 | 44.345 | 54.896 | 999.000 | 42.504 | 171.408 | 177.539 | 123.448 | 133.935 |
| **197P** | 999.000 | 133.935 | 62.779 | 44.345 | 32.034 | 999.000 | 176.723 | 171.408 | 109.778 | 122.404 |
| **198R** | 8.601 | 122.404 | 53.175 | 62.779 | 32.514 | 32.034 | 174.403 | 176.723 | 133.935 | 137.212 |
| **199P** | 999.000 | 137.212 | 62.930 | 53.175 | 32.157 | 32.514 | 176.873 | 174.403 | 122.404 | 121.973 |
| **200R** | 8.636 | 121.973 | 56.259 | 62.930 | 30.784 | 32.157 | 176.464 | 176.873 | 137.212 | 116.886 |
| **201S** | 8.428 | 116.886 | 58.141 | 56.259 | 63.741 | 30.784 | 174.255 | 176.464 | 121.973 | 121.965 |
| **202H** | 8.427 | 121.965 | 56.409 | 58.141 | 30.990 | 63.741 | 175.086 | 174.255 | 116.886 | 120.372 |
| **203N** | 8.416 | 120.372 | 53.191 | 56.409 | 38.745 | 30.990 | 174.931 | 175.086 | 121.965 | 122.498 |
| **204K** | 8.493 | 122.498 | 56.436 | 53.191 | 32.879 | 38.745 | 176.349 | 174.931 | 120.372 | 122.495 |
| **205V** | 8.251 | 122.495 | 62.413 | 56.436 | 32.573 | 32.879 | 175.966 | 176.349 | 122.498 | 126.463 |

| Residue number (JPT2) | H (ppm) | N (ppm) | CA (ppm) | CA(i-1) (ppm) | CB (ppm) | CB(i-1) (ppm) | C (ppm) | C(i-1) (ppm) | N(i-1) (ppm) | N(i+1) (ppm) |
|---|---|---|---|---|---|---|---|---|---|---|
| **206L** | 8.404 | 126.463 | 54.985 | 62.413 | 42.353 | 32.573 | 176.581 | 175.966 | 122.495 | 120.157 |
| **207N** | 8.523 | 120.157 | 51.230 | 54.985 | 38.621 | 42.353 | 172.187 | 176.581 | 126.463 | 137.861 |
| **208P** | 999.000 | 137.861 | 61.367 | 51.230 | 30.598 | 38.621 | 999.000 | 172.187 | 120.157 | 999.000 |
| **210G** | 8.619 | 109.602 | 45.204 | 999.000 | 999.000 | 999.000 | 174.789 | 999.000 | 999.000 | 108.664 |
| **211G** | 8.299 | 108.664 | 999.000 | 45.204 | 999.000 | 999.000 | 174.142 | 174.789 | 109.602 | 121.039 |
| **212K** | 8.402 | 121.039 | 56.366 | 999.000 | 32.918 | 999.000 | 176.820 | 174.142 | 108.664 | 117.301 |
| **213S** | 8.518 | 117.301 | 58.252 | 56.366 | 63.782 | 32.918 | 174.612 | 176.820 | 121.039 | 118.270 |
| **214S** | 8.465 | 118.270 | 58.408 | 58.252 | 63.662 | 63.782 | 174.296 | 174.612 | 117.301 | 121.971 |
| **215I** | 8.140 | 121.971 | 61.148 | 58.408 | 38.709 | 63.662 | 175.938 | 174.296 | 118.270 | 119.811 |
| **216S** | 8.277 | 119.811 | 57.876 | 61.148 | 63.943 | 38.709 | 173.599 | 175.938 | 121.971 | 122.703 |
| **217F** | 8.287 | 122.703 | 57.858 | 57.876 | 39.792 | 63.943 | 174.384 | 173.599 | 119.811 | 125.571 |
| **218Y** | 7.712 | 125.571 | 59.168 | 57.858 | 39.553 | 39.792 | 173.240 | 174.384 | 122.703 | 999.000 |

**Supplementary Table 6** Peaklist of backbone chemical shifts for CBP-TAZ4 used to validate SpinForecast, obtained from BMRB entry 52655. Residue numbers correspond to the sequence in Supplementary Table 1. Missing values are denoted by 999.000.

| Residue (CBP-TAZ4) | H (ppm) | N (ppm) | CA (ppm) | CA(i-1) (ppm) | CB (ppm) | CB(i-1) (ppm) | C (ppm) | C(i-1) (ppm) | N(i-1) (ppm) | N(i+1) (ppm) |
|---|---|---|---|---|---|---|---|---|---|---|
| **2G** | 999.000 | 999.000 | 43.120 | 999.000 | 999.000 | 999.000 | 999.000 | 999.000 | 999.000 | 116.980 |
| **3S** | 8.730 | 116.980 | 56.570 | 43.120 | 63.150 | 999.000 | 173.190 | 999.000 | 999.000 | 999.000 |
| **4P** | 999.000 | 999.000 | 64.090 | 56.570 | 31.750 | 63.150 | 177.930 | 173.190 | 116.980 | 119.020 |
| **5Q** | 8.480 | 119.020 | 57.300 | 64.090 | 28.530 | 31.750 | 177.200 | 177.930 | 999.000 | 122.080 |
| **6E** | 8.220 | 122.080 | 57.680 | 57.300 | 29.770 | 28.530 | 177.470 | 177.200 | 119.020 | 116.300 |
| **7S** | 8.320 | 116.300 | 59.890 | 57.680 | 63.040 | 29.770 | 176.050 | 177.470 | 122.080 | 122.710 |
| **8R** | 8.210 | 122.710 | 58.080 | 59.890 | 30.020 | 63.040 | 177.550 | 176.050 | 116.300 | 120.170 |
| **9R** | 7.990 | 120.170 | 58.590 | 58.080 | 30.070 | 30.020 | 178.090 | 177.550 | 122.710 | 120.770 |
| **10L** | 8.150 | 120.770 | 57.110 | 58.590 | 41.530 | 30.070 | 178.750 | 178.090 | 120.170 | 115.810 |
| **11S** | 8.110 | 115.810 | 60.980 | 57.110 | 62.550 | 41.530 | 177.290 | 178.750 | 120.770 | 123.930 |
| **12I** | 8.070 | 123.930 | 64.190 | 60.980 | 37.690 | 62.550 | 177.130 | 177.290 | 115.810 | 119.260 |
| **13Q** | 8.040 | 119.260 | 59.290 | 64.190 | 28.110 | 37.690 | 179.110 | 177.130 | 123.930 | 118.810 |
| **14R** | 8.200 | 118.810 | 59.000 | 59.290 | 29.910 | 28.110 | 178.960 | 179.110 | 119.260 | 124.160 |
| **15A** | 7.970 | 124.160 | 55.210 | 59.000 | 18.540 | 29.910 | 180.490 | 178.960 | 118.810 | 118.670 |
| **16I** | 8.670 | 118.670 | 65.200 | 55.210 | 37.450 | 18.540 | 177.650 | 180.490 | 124.160 | 118.950 |
| **17Q** | 8.110 | 118.950 | 59.210 | 65.200 | 27.840 | 37.450 | 179.530 | 177.650 | 118.670 | 115.800 |
| **18S** | 8.130 | 115.800 | 61.720 | 59.210 | 69.580 | 27.840 | 174.110 | 179.530 | 118.950 | 999.000 |
| **19L** | 999.000 | 999.000 | 58.320 | 61.720 | 42.040 | 69.580 | 176.840 | 174.110 | 115.800 | 118.610 |
| **20V** | 8.550 | 118.610 | 66.670 | 58.320 | 31.620 | 42.040 | 178.400 | 176.840 | 999.000 | 115.630 |
| **21H** | 7.940 | 115.630 | 60.030 | 66.670 | 27.550 | 31.620 | 177.950 | 178.400 | 118.610 | 121.320 |
| **22A** | 8.580 | 121.320 | 54.990 | 60.030 | 19.620 | 27.550 | 179.110 | 177.950 | 115.630 | 116.310 |
| **23A** | 8.250 | 116.310 | 53.640 | 54.990 | 18.200 | 19.620 | 177.230 | 179.110 | 121.320 | 111.660 |
| **24Q** | 7.200 | 111.660 | 54.250 | 53.640 | 30.440 | 18.200 | 174.150 | 177.230 | 116.310 | 124.360 |
| **25C** | 7.240 | 124.360 | 60.560 | 54.250 | 28.920 | 30.440 | 176.020 | 174.150 | 111.660 | 129.540 |
| **26R** | 8.890 | 129.540 | 54.500 | 60.560 | 29.990 | 28.920 | 175.050 | 176.020 | 124.360 | 120.280 |
| **27N** | 7.860 | 120.280 | 51.620 | 54.500 | 38.780 | 29.990 | 175.590 | 175.050 | 129.540 | 130.310 |
| **28A** | 9.170 | 130.310 | 54.300 | 51.620 | 18.160 | 38.780 | 177.960 | 175.590 | 120.280 | 113.970 |
| **29N** | 8.050 | 113.970 | 51.760 | 54.300 | 38.620 | 18.160 | 173.970 | 177.960 | 130.310 | 123.090 |
| **30C** | 7.140 | 123.090 | 61.510 | 51.760 | 29.790 | 38.620 | 176.760 | 173.970 | 113.970 | 124.510 |
| **31S** | 8.840 | 124.510 | 57.780 | 61.510 | 63.840 | 29.790 | 174.680 | 176.760 | 123.090 | 128.650 |
| **32L** | 8.500 | 128.650 | 54.110 | 57.780 | 40.620 | 63.840 | 176.630 | 174.680 | 124.510 | 999.000 |
| **33P** | 999.000 | 999.000 | 65.270 | 54.110 | 31.720 | 40.620 | 178.300 | 176.630 | 128.650 | 111.970 |
| **34S** | 8.380 | 111.970 | 60.280 | 65.270 | 62.690 | 31.720 | 174.290 | 178.300 | 999.000 | 125.710 |
| **35C** | 7.500 | 125.710 | 63.640 | 60.280 | 29.520 | 62.690 | 176.520 | 174.290 | 111.970 | 116.240 |
| **36Q** | 8.200 | 116.240 | 59.710 | 63.640 | 28.000 | 29.520 | 179.200 | 176.520 | 125.710 | 117.140 |
| **37K** | 8.010 | 117.140 | 59.400 | 59.710 | 32.630 | 28.000 | 179.320 | 179.200 | 116.240 | 118.060 |
| **38M** | 8.190 | 118.060 | 56.540 | 59.400 | 31.630 | 32.630 | 178.890 | 179.320 | 117.140 | 122.110 |
| **39K** | 9.600 | 122.110 | 61.270 | 56.540 | 32.450 | 31.630 | 179.610 | 178.890 | 118.060 | 117.690 |

| Residue (CBP-TAZ4) | H (ppm) | N (ppm) | CA (ppm) | CA(i-1) (ppm) | CB (ppm) | CB(i-1) (ppm) | C (ppm) | C(i-1) (ppm) | N(i-1) (ppm) | N(i+1) (ppm) |
|---|---|---|---|---|---|---|---|---|---|---|
| **40R** | 7.550 | 117.690 | 59.450 | 61.270 | 29.680 | 32.450 | 179.650 | 179.610 | 122.110 | 121.530 |
| **41V** | 7.540 | 121.530 | 66.310 | 59.450 | 33.640 | 29.680 | 177.830 | 179.650 | 117.690 | 120.830 |
| **42V** | 8.480 | 120.830 | 67.300 | 66.310 | 31.980 | 33.640 | 999.000 | 177.830 | 121.530 | 999.000 |
| **44H** | 999.000 | 999.000 | 59.730 | 999.000 | 27.110 | 999.000 | 178.030 | 999.000 | 999.000 | 110.250 |
| **45T** | 7.900 | 110.250 | 65.480 | 59.730 | 69.440 | 27.110 | 175.730 | 178.030 | 999.000 | 120.130 |
| **46K** | 7.270 | 120.130 | 58.470 | 65.480 | 31.890 | 69.440 | 177.670 | 175.730 | 110.250 | 103.950 |
| **47G** | 7.160 | 103.950 | 44.500 | 58.470 | 999.000 | 31.890 | 173.690 | 177.670 | 120.130 | 123.200 |
| **48C** | 6.880 | 123.200 | 60.860 | 44.500 | 29.630 | 999.000 | 176.610 | 173.690 | 103.950 | 128.100 |
| **49K** | 9.000 | 128.100 | 56.350 | 60.860 | 32.610 | 29.630 | 177.360 | 176.610 | 123.200 | 122.470 |
| **50R** | 8.510 | 122.470 | 57.400 | 56.350 | 31.510 | 32.610 | 176.750 | 177.360 | 128.100 | 118.730 |
| **51K** | 8.450 | 118.730 | 57.490 | 57.400 | 32.030 | 31.510 | 177.100 | 176.750 | 122.470 | 113.240 |
| **52T** | 7.820 | 113.240 | 63.720 | 57.490 | 68.450 | 32.030 | 175.910 | 177.100 | 118.730 | 119.660 |
| **53N** | 8.370 | 119.660 | 53.870 | 63.720 | 38.080 | 68.450 | 175.860 | 175.910 | 113.240 | 107.360 |
| **54G** | 7.960 | 107.360 | 45.620 | 53.870 | 999.000 | 38.080 | 175.060 | 175.860 | 119.660 | 107.200 |
| **55G** | 7.810 | 107.200 | 45.370 | 45.620 | 999.000 | 999.000 | 173.190 | 175.060 | 107.360 | 122.430 |
| **56C** | 8.030 | 122.430 | 59.670 | 45.370 | 30.420 | 999.000 | 174.480 | 173.190 | 107.200 | 999.000 |
| **57P** | 999.000 | 999.000 | 64.820 | 59.670 | 32.140 | 30.420 | 178.790 | 174.480 | 122.430 | 122.500 |
| **58V** | 7.810 | 122.500 | 65.920 | 64.820 | 32.470 | 32.140 | 178.910 | 178.790 | 999.000 | 124.250 |
| **59C** | 8.900 | 124.250 | 66.070 | 65.920 | 29.160 | 32.470 | 177.510 | 178.910 | 122.500 | 117.330 |
| **60K** | 8.090 | 117.330 | 59.920 | 66.070 | 32.350 | 29.160 | 179.070 | 177.510 | 124.250 | 118.000 |
| **61Q** | 7.380 | 118.000 | 58.280 | 59.920 | 28.220 | 32.350 | 178.250 | 179.070 | 117.330 | 120.620 |
| **62L** | 7.810 | 120.620 | 57.670 | 58.280 | 41.610 | 28.220 | 178.930 | 178.250 | 118.000 | 120.270 |
| **63I** | 8.480 | 120.270 | 65.420 | 57.670 | 37.320 | 41.610 | 176.930 | 178.930 | 120.620 | 121.680 |
| **64A** | 7.310 | 121.680 | 54.880 | 65.420 | 17.790 | 37.320 | 180.790 | 176.930 | 120.270 | 117.520 |
| **65L** | 7.550 | 117.520 | 57.550 | 54.880 | 42.450 | 17.790 | 178.970 | 180.790 | 121.680 | 121.480 |
| **66A** | 8.870 | 121.480 | 55.630 | 57.550 | 18.940 | 42.450 | 179.300 | 178.970 | 117.520 | 119.330 |
| **67A** | 8.600 | 119.330 | 55.410 | 55.630 | 17.550 | 18.940 | 179.730 | 179.300 | 121.480 | 117.630 |
| **68Y** | 7.510 | 117.630 | 61.130 | 55.410 | 38.340 | 17.550 | 178.830 | 179.730 | 119.330 | 999.000 |
| **69H** | 999.000 | 999.000 | 58.590 | 61.130 | 27.750 | 38.340 | 176.720 | 178.830 | 117.630 | 122.060 |
| **70A** | 9.220 | 122.060 | 54.600 | 58.590 | 19.750 | 27.750 | 178.670 | 176.720 | 999.000 | 113.560 |
| **71K** | 6.910 | 113.560 | 58.290 | 54.600 | 32.440 | 19.750 | 176.370 | 178.670 | 122.060 | 112.470 |
| **72H** | 7.100 | 112.470 | 53.910 | 58.290 | 30.050 | 32.440 | 173.320 | 176.370 | 113.560 | 123.270 |
| **73C** | 6.910 | 123.270 | 59.820 | 53.910 | 29.780 | 30.050 | 176.060 | 173.320 | 112.470 | 128.520 |
| **74Q** | 9.170 | 128.520 | 54.510 | 59.820 | 29.600 | 29.780 | 175.490 | 176.060 | 123.270 | 999.000 |
| **75E** | 999.000 | 999.000 | 55.560 | 54.510 | 30.110 | 29.600 | 176.650 | 175.490 | 128.520 | 126.170 |
| **76N** | 9.000 | 126.170 | 55.070 | 55.560 | 38.220 | 30.110 | 175.810 | 176.650 | 999.000 | 120.790 |
| **77K** | 8.810 | 120.790 | 54.220 | 55.070 | 30.740 | 38.220 | 174.770 | 175.810 | 126.170 | 123.520 |
| **78C** | 6.780 | 123.520 | 57.780 | 54.220 | 30.410 | 30.740 | 175.830 | 174.770 | 120.790 | 999.000 |
| **79P** | 999.000 | 999.000 | 62.740 | 57.780 | 31.880 | 30.410 | 176.770 | 175.830 | 123.520 | 128.050 |
| **80V** | 9.060 | 128.050 | 62.230 | 62.740 | 32.610 | 31.880 | 173.960 | 176.770 | 999.000 | 999.000 |
| **82F** | 8.010 | 111.250 | 60.740 | 999.000 | 37.020 | 999.000 | 175.350 | 999.000 | 999.000 | 123.800 |

| Residue (CBP-TAZ4) | H (ppm) | N (ppm) | CA (ppm) | CA(i-1) (ppm) | CB (ppm) | CB(i-1) (ppm) | C (ppm) | C(i-1) (ppm) | N(i-1) (ppm) | N(i+1) (ppm) |
|---|---|---|---|---|---|---|---|---|---|---|
| **83C** | 7.780 | 123.800 | 64.880 | 60.740 | 29.870 | 37.020 | 176.830 | 175.350 | 111.250 | 999.000 |
| **97H** | 999.000 | 999.000 | 56.260 | 999.000 | 29.390 | 999.000 | 175.970 | 999.000 | 999.000 | 121.460 |
| **98R** | 8.350 | 121.460 | 56.030 | 56.260 | 28.960 | 29.390 | 176.070 | 175.970 | 999.000 | 122.810 |
| **99L** | 8.260 | 122.810 | 55.320 | 56.030 | 42.240 | 28.960 | 177.380 | 176.070 | 121.460 | 120.740 |
| **100Q** | 8.330 | 120.740 | 56.430 | 55.320 | 30.190 | 42.240 | 176.280 | 177.380 | 122.810 | 120.580 |
| **101Q** | 8.330 | 120.580 | 57.940 | 56.430 | 29.170 | 30.190 | 177.660 | 176.280 | 120.740 | 122.630 |
| **102A** | 8.090 | 122.630 | 54.260 | 57.940 | 18.110 | 29.170 | 179.730 | 177.660 | 120.580 | 118.030 |
| **103Q** | 8.030 | 118.030 | 57.720 | 54.260 | 28.450 | 18.110 | 177.820 | 179.730 | 122.630 | 121.200 |
| **104L** | 7.970 | 121.200 | 57.030 | 57.720 | 41.810 | 28.450 | 178.760 | 177.820 | 118.030 | 118.760 |
| **105M** | 8.050 | 118.760 | 56.940 | 57.030 | 31.960 | 41.810 | 177.480 | 178.760 | 121.200 | 999.000 |
| **109M** | 999.000 | 999.000 | 55.950 | 999.000 | 32.670 | 999.000 | 176.560 | 999.000 | 999.000 | 123.760 |
| **110A** | 8.130 | 123.760 | 52.890 | 55.950 | 18.900 | 32.670 | 178.220 | 176.560 | 999.000 | 112.010 |
| **111T** | 7.970 | 112.010 | 62.050 | 52.890 | 69.500 | 18.900 | 174.820 | 178.220 | 123.760 | 121.990 |
| **112M** | 8.200 | 121.990 | 55.740 | 62.050 | 32.690 | 69.500 | 175.920 | 174.820 | 112.010 | 119.720 |
| **113N** | 8.430 | 119.720 | 53.160 | 55.740 | 38.620 | 32.690 | 175.420 | 175.920 | 121.990 | 114.280 |
| **114T** | 8.090 | 114.280 | 61.990 | 53.160 | 69.560 | 38.620 | 174.500 | 175.420 | 119.720 | 122.560 |
| **115R** | 8.280 | 122.560 | 56.130 | 61.990 | 30.570 | 69.560 | 175.800 | 174.500 | 114.280 | 119.980 |
| **116N** | 8.420 | 119.980 | 52.970 | 56.130 | 38.640 | 30.570 | 174.590 | 175.800 | 122.560 | 121.590 |
| **117V** | 8.040 | 121.590 | 59.550 | 52.970 | 32.360 | 38.640 | 174.290 | 174.590 | 119.980 | 139.360 |
| **118P** | 999.000 | 139.360 | 63.210 | 59.550 | 31.870 | 32.360 | 176.850 | 174.290 | 121.590 | 120.930 |
| **119Q** | 8.500 | 120.930 | 55.640 | 63.210 | 29.350 | 31.870 | 176.020 | 176.850 | 139.360 | 121.560 |
| **120Q** | 8.460 | 121.560 | 55.760 | 55.640 | 29.540 | 29.350 | 175.710 | 176.020 | 120.930 | 117.550 |
| **121S** | 8.380 | 117.550 | 57.960 | 55.760 | 63.660 | 29.540 | 173.850 | 175.710 | 121.560 | 125.280 |
| **122L** | 8.260 | 125.280 | 52.770 | 57.960 | 41.670 | 63.660 | 175.090 | 173.850 | 117.550 | 136.130 |
| **123P** | 999.000 | 136.130 | 62.780 | 52.770 | 31.870 | 41.670 | 176.620 | 175.090 | 125.280 | 117.570 |
| **124S** | 8.420 | 117.570 | 56.090 | 62.780 | 63.200 | 31.870 | 173.110 | 176.620 | 136.130 | 138.060 |
| **125P** | 999.000 | 138.060 | 63.230 | 56.090 | 31.880 | 63.200 | 177.140 | 173.110 | 117.570 | 113.330 |
| **126T** | 8.180 | 113.330 | 61.590 | 63.230 | 69.620 | 31.880 | 174.490 | 177.140 | 138.060 | 117.990 |
| **127S** | 8.170 | 117.990 | 57.950 | 61.590 | 63.920 | 69.620 | 173.500 | 174.490 | 113.330 | 126.990 |
| **128A** | 8.270 | 126.990 | 50.390 | 57.950 | 18.130 | 63.920 | 174.880 | 173.500 | 117.990 | 137.260 |
| **129P** | 999.000 | 137.260 | 999.000 | 50.390 | 999.000 | 18.130 | 174.850 | 174.880 | 126.990 | 135.840 |
| **130P** | 999.000 | 135.840 | 63.150 | 999.000 | 31.870 | 999.000 | 177.440 | 174.850 | 137.260 | 109.360 |
| **131G** | 8.490 | 109.360 | 44.910 | 63.150 | 999.000 | 31.870 | 174.000 | 177.440 | 135.840 | 116.650 |
| **132T** | 8.000 | 116.650 | 59.980 | 44.910 | 69.640 | 999.000 | 172.930 | 174.000 | 109.360 | 139.420 |
| **133P** | 999.000 | 139.420 | 63.240 | 59.980 | 31.880 | 69.640 | 177.020 | 172.930 | 116.650 | 114.800 |
| **134T** | 8.250 | 114.800 | 61.840 | 63.240 | 69.670 | 31.880 | 174.410 | 177.020 | 139.420 | 122.780 |
| **135Q** | 8.360 | 122.780 | 55.430 | 61.840 | 29.400 | 69.670 | 175.460 | 174.410 | 114.800 | 123.320 |
| **136Q** | 8.500 | 123.320 | 53.520 | 55.430 | 28.740 | 29.400 | 173.940 | 175.460 | 122.780 | 137.350 |
| **137P** | 999.000 | 137.350 | 62.810 | 53.520 | 31.890 | 28.740 | 176.760 | 173.940 | 123.320 | 116.420 |
| **138S** | 8.480 | 116.420 | 58.060 | 62.810 | 63.670 | 31.890 | 174.290 | 176.760 | 137.350 | 117.990 |
| **139T** | 8.160 | 117.990 | 59.500 | 58.060 | 69.660 | 63.670 | 172.800 | 174.290 | 116.420 | 138.760 |

| Residue (CBP-TAZ4) | H (ppm) | N (ppm) | CA (ppm) | CA(i-1) (ppm) | CB (ppm) | CB(i-1) (ppm) | C (ppm) | C(i-1) (ppm) | N(i-1) (ppm) | N(i+1) (ppm) |
|---|---|---|---|---|---|---|---|---|---|---|
| **140P** | 999.000 | 138.760 | 63.150 | 59.500 | 31.880 | 69.660 | 176.800 | 172.800 | 117.990 | 120.840 |
| **141Q** | 8.520 | 120.840 | 55.410 | 63.150 | 29.440 | 31.880 | 175.930 | 176.800 | 138.760 | 118.250 |
| **142T** | 8.230 | 118.250 | 59.730 | 55.410 | 69.660 | 29.440 | 172.630 | 175.930 | 120.840 | 138.990 |
| **143P** | 999.000 | 138.990 | 63.060 | 59.730 | 31.870 | 69.660 | 176.600 | 172.630 | 118.250 | 122.180 |
| **144Q** | 8.470 | 122.180 | 53.010 | 63.060 | 28.950 | 31.870 | 173.590 | 176.600 | 138.990 | 138.880 |
| **145P** | 999.000 | 138.880 | 999.000 | 53.010 | 999.000 | 28.950 | 174.500 | 173.590 | 122.180 | 135.390 |
| **146P** | 999.000 | 135.390 | 62.740 | 999.000 | 31.800 | 999.000 | 176.510 | 174.500 | 138.880 | 124.320 |
| **147A** | 8.360 | 124.320 | 52.080 | 62.740 | 19.030 | 31.800 | 177.490 | 176.510 | 135.390 | 120.550 |
| **148Q** | 8.330 | 120.550 | 53.340 | 52.080 | 29.750 | 19.030 | 173.900 | 177.490 | 124.320 | 137.180 |
| **149P** | 999.000 | 137.180 | 62.890 | 53.340 | 31.970 | 29.750 | 176.560 | 173.900 | 120.550 | 121.660 |
| **150Q** | 8.490 | 121.660 | 53.380 | 62.890 | 28.870 | 31.970 | 173.970 | 176.560 | 137.180 | 137.170 |
| **151P** | 999.000 | 137.170 | 62.810 | 53.380 | 31.890 | 28.870 | 176.590 | 173.970 | 121.660 | 117.810 |
| **152S** | 8.440 | 117.810 | 56.240 | 62.810 | 63.200 | 31.890 | 172.860 | 176.590 | 137.170 | 137.980 |
| **153P** | 999.000 | 137.980 | 63.200 | 56.240 | 31.860 | 63.200 | 176.930 | 172.860 | 117.810 | 119.810 |
| **154V** | 8.150 | 119.810 | 62.170 | 63.200 | 32.540 | 31.860 | 176.160 | 176.930 | 137.980 | 119.080 |
| **155S** | 8.300 | 119.080 | 57.940 | 62.170 | 63.690 | 32.540 | 174.290 | 176.160 | 119.810 | 122.650 |
| **156M** | 8.380 | 122.650 | 54.930 | 57.940 | 32.820 | 63.690 | 175.860 | 174.290 | 119.080 | 118.540 |
| **157S** | 8.350 | 118.540 | 56.230 | 54.930 | 63.130 | 32.820 | 172.770 | 175.860 | 122.650 | 138.210 |
| **158P** | 999.000 | 138.210 | 63.310 | 56.230 | 31.630 | 63.130 | 176.670 | 172.770 | 118.540 | 123.440 |
| **159A** | 8.250 | 123.440 | 52.420 | 63.310 | 18.920 | 31.630 | 178.030 | 176.670 | 138.210 | 107.390 |
| **160G** | 8.190 | 107.390 | 44.790 | 52.420 | 999.000 | 18.920 | 173.350 | 178.030 | 123.440 | 120.710 |
| **161F** | 7.980 | 120.710 | 55.500 | 44.790 | 38.880 | 999.000 | 173.910 | 173.350 | 107.390 | 137.310 |
| **162P** | 999.000 | 137.310 | 63.070 | 55.500 | 31.880 | 38.880 | 176.710 | 173.910 | 120.710 | 116.420 |
| **163S** | 8.380 | 116.420 | 58.180 | 63.070 | 63.590 | 31.880 | 174.640 | 176.710 | 137.310 | 121.650 |
| **164V** | 8.120 | 121.650 | 61.910 | 58.180 | 32.750 | 63.590 | 175.690 | 174.640 | 116.420 | 127.650 |
| **165A** | 8.340 | 127.650 | 52.160 | 61.910 | 19.070 | 32.750 | 177.510 | 175.690 | 121.650 | 120.680 |
| **166R** | 8.340 | 120.680 | 55.880 | 52.160 | 30.670 | 19.070 | 176.390 | 177.510 | 127.650 | 115.640 |
| **167T** | 8.160 | 115.640 | 61.520 | 55.880 | 69.750 | 30.670 | 174.100 | 176.390 | 120.680 | 124.020 |
| **168Q** | 8.410 | 124.020 | 53.380 | 61.520 | 28.990 | 69.750 | 173.360 | 174.100 | 115.640 | 138.800 |
| **169P** | 999.000 | 138.800 | 999.000 | 53.380 | 999.000 | 28.990 | 174.640 | 173.360 | 124.020 | 135.270 |
| **170P** | 999.000 | 135.270 | 62.840 | 999.000 | 31.920 | 999.000 | 176.990 | 174.640 | 138.800 | 114.370 |
| **171T** | 8.260 | 114.370 | 61.720 | 62.840 | 69.690 | 31.920 | 174.670 | 176.990 | 135.270 | 116.810 |
| **172T** | 8.170 | 116.810 | 61.600 | 61.720 | 69.830 | 69.690 | 174.330 | 174.670 | 114.370 | 122.620 |
| **173V** | 8.200 | 122.620 | 62.010 | 61.600 | 32.610 | 69.830 | 176.100 | 174.330 | 116.810 | 119.850 |
| **174S** | 8.480 | 119.850 | 57.960 | 62.010 | 63.680 | 32.610 | 174.830 | 176.100 | 122.620 | 115.750 |
| **175T** | 8.220 | 115.750 | 61.750 | 57.960 | 69.620 | 63.680 | 175.050 | 174.830 | 119.850 | 110.920 |
| **176G** | 8.380 | 110.920 | 45.070 | 61.750 | 999.000 | 69.620 | 173.570 | 175.050 | 115.750 | 121.990 |
| **177K** | 8.140 | 121.990 | 53.870 | 45.070 | 32.450 | 999.000 | 174.580 | 173.570 | 110.920 | 137.350 |
| **178P** | 999.000 | 137.350 | 63.090 | 53.870 | 31.940 | 32.450 | 177.160 | 174.580 | 121.990 | 114.530 |
| **179T** | 8.310 | 114.530 | 61.700 | 63.090 | 69.630 | 31.940 | 174.590 | 177.160 | 137.350 | 117.680 |
| **180S** | 8.260 | 117.680 | 58.050 | 61.700 | 63.760 | 69.630 | 174.200 | 174.590 | 114.530 | 122.440 |

| Residue (CBP-TAZ4) | H (ppm) | N (ppm) | CA (ppm) | CA(i-1) (ppm) | CB (ppm) | CB(i-1) (ppm) | C (ppm) | C(i-1) (ppm) | N(i-1) (ppm) | N(i+1) (ppm) |
|---|---|---|---|---|---|---|---|---|---|---|
| **181Q** | 8.430 | 122.440 | 55.470 | 58.050 | 29.350 | 63.760 | 175.570 | 174.200 | 117.680 | 123.170 |
| **182V** | 8.250 | 123.170 | 59.750 | 55.470 | 32.330 | 29.350 | 174.290 | 175.570 | 122.440 | 139.760 |
| **183P** | 999.000 | 139.760 | 62.910 | 59.750 | 31.980 | 32.330 | 176.040 | 174.290 | 123.170 | 125.720 |
| **184A** | 8.320 | 125.720 | 50.070 | 62.910 | 17.950 | 31.980 | 175.000 | 176.040 | 139.760 | 136.850 |
| **185P** | 999.000 | 136.850 | 999.000 | 50.070 | 999.000 | 17.950 | 174.190 | 175.000 | 125.720 | 136.560 |
| **186P** | 999.000 | 136.560 | 999.000 | 999.000 | 999.000 | 999.000 | 174.160 | 174.190 | 136.850 | 136.230 |
| **187P** | 999.000 | 136.230 | 999.000 | 999.000 | 999.000 | 999.000 | 174.590 | 174.160 | 136.560 | 135.070 |
| **188P** | 999.000 | 135.070 | 62.660 | 999.000 | 31.880 | 999.000 | 176.580 | 174.590 | 136.230 | 124.320 |
| **189A** | 8.350 | 124.320 | 52.190 | 62.660 | 19.090 | 31.880 | 177.490 | 176.580 | 135.070 | 120.550 |
| **190Q** | 8.300 | 120.550 | 53.190 | 52.190 | 28.940 | 19.090 | 173.440 | 177.490 | 124.320 | 138.770 |
| **191P** | 999.000 | 138.770 | 999.000 | 53.190 | 999.000 | 28.940 | 174.000 | 173.440 | 120.550 | 136.310 |
| **192P** | 999.000 | 136.310 | 999.000 | 999.000 | 999.000 | 999.000 | 174.630 | 174.000 | 138.770 | 135.110 |
| **193P** | 999.000 | 135.110 | 63.210 | 999.000 | 31.790 | 999.000 | 177.030 | 174.630 | 136.310 | 123.430 |
| **194A** | 8.410 | 123.430 | 52.810 | 63.210 | 18.870 | 31.790 | 178.150 | 177.030 | 135.110 | 122.670 |
| **195A** | 8.230 | 122.670 | 52.970 | 52.810 | 18.770 | 18.870 | 178.530 | 178.150 | 123.430 | 119.580 |
| **196V** | 7.970 | 119.580 | 63.150 | 52.970 | 32.290 | 18.770 | 176.880 | 178.530 | 122.670 | 123.260 |
| **197E** | 8.340 | 123.260 | 57.250 | 63.150 | 29.730 | 32.290 | 177.140 | 176.880 | 119.580 | 124.280 |
| **198A** | 8.220 | 124.280 | 53.410 | 57.250 | 18.550 | 29.730 | 178.470 | 177.140 | 123.260 | 121.680 |
| **199A** | 8.070 | 121.680 | 53.260 | 53.410 | 18.500 | 18.550 | 178.730 | 178.470 | 124.280 | 119.320 |
| **200R** | 8.110 | 119.320 | 56.990 | 53.260 | 30.490 | 18.500 | 177.220 | 178.730 | 121.680 | 999.000 |
| **201Q** | 999.000 | 999.000 | 56.830 | 56.990 | 28.490 | 30.490 | 176.720 | 177.220 | 119.320 | 121.290 |
| **202I** | 8.030 | 121.290 | 62.410 | 56.830 | 38.290 | 28.490 | 177.180 | 176.720 | 999.000 | 123.220 |
| **203E** | 8.210 | 123.220 | 57.450 | 62.410 | 29.700 | 38.290 | 177.400 | 177.180 | 121.290 | 121.080 |
| **204R** | 8.230 | 121.080 | 57.080 | 57.450 | 30.360 | 29.700 | 177.570 | 177.400 | 123.220 | 120.660 |
| **205E** | 8.440 | 120.660 | 57.670 | 57.080 | 29.480 | 30.360 | 177.330 | 177.570 | 121.080 | 123.560 |
| **206A** | 8.170 | 123.560 | 53.440 | 57.670 | 18.490 | 29.480 | 178.840 | 177.330 | 120.660 | 118.080 |
| **207Q** | 8.160 | 118.080 | 56.820 | 53.440 | 28.800 | 18.490 | 176.990 | 178.840 | 123.560 | 119.980 |
| **208Q** | 8.210 | 119.980 | 56.570 | 56.820 | 28.930 | 28.800 | 176.400 | 176.990 | 118.080 | 119.910 |
| **209Q** | 8.240 | 119.910 | 56.440 | 56.570 | 29.030 | 28.930 | 176.400 | 176.400 | 119.980 | 119.750 |
| **210Q** | 8.220 | 119.750 | 56.410 | 56.440 | 28.950 | 29.030 | 176.110 | 176.400 | 119.910 | 118.490 |
| **211H** | 8.320 | 118.490 | 55.760 | 56.410 | 29.120 | 28.950 | 174.610 | 176.110 | 119.750 | 122.120 |
| **212L** | 8.050 | 122.120 | 55.430 | 55.760 | 42.180 | 29.120 | 176.850 | 174.610 | 118.490 | 120.370 |
| **213Y** | 8.090 | 120.370 | 57.710 | 55.430 | 38.450 | 42.180 | 175.490 | 176.850 | 122.120 | 122.530 |
| **214R** | 8.110 | 122.530 | 55.810 | 57.710 | 30.820 | 38.450 | 175.750 | 175.490 | 120.370 | 120.640 |
| **215V** | 8.060 | 120.640 | 62.100 | 55.810 | 32.700 | 30.820 | 175.630 | 175.750 | 122.530 | 122.320 |
| **216N** | 8.470 | 122.320 | 52.940 | 62.100 | 38.760 | 32.700 | 175.180 | 175.630 | 120.640 | 121.220 |
| **217I** | 8.110 | 121.220 | 61.330 | 52.940 | 38.510 | 38.760 | 175.900 | 175.180 | 122.320 | 121.660 |
| **218N** | 8.450 | 121.660 | 53.280 | 61.330 | 38.630 | 38.510 | 174.930 | 175.900 | 121.220 | 119.520 |
| **219N** | 8.350 | 119.520 | 53.360 | 53.280 | 38.620 | 38.630 | 175.040 | 174.930 | 121.660 | 115.660 |
| **220S** | 8.210 | 115.660 | 58.380 | 53.360 | 63.630 | 38.620 | 174.010 | 175.040 | 119.520 | 122.930 |
| **221M** | 8.240 | 122.930 | 57.540 | 58.380 | 32.290 | 63.630 | 173.560 | 174.010 | 115.660 | 138.710 |

| Residue (CBP-TAZ4) | H (ppm) | N (ppm) | CA (ppm) | CA(i-1) (ppm) | CB (ppm) | CB(i-1) (ppm) | C (ppm) | C(i-1) (ppm) | N(i-1) (ppm) | N(i+1) (ppm) |
|---|---|---|---|---|---|---|---|---|---|---|
| **222P** | 999.000 | 138.710 | 999.000 | 57.540 | 999.000 | 32.290 | 174.690 | 173.560 | 122.930 | 135.250 |
| **223P** | 999.000 | 135.250 | 63.170 | 999.000 | 31.970 | 999.000 | 177.560 | 174.690 | 138.710 | 108.760 |
| **224G** | 8.430 | 108.760 | 45.150 | 63.170 | 999.000 | 31.970 | 174.160 | 177.560 | 135.250 | 120.410 |
| **225R** | 8.200 | 120.410 | 55.880 | 45.150 | 30.740 | 999.000 | 176.510 | 174.160 | 108.760 | 114.670 |
| **226T** | 8.210 | 114.670 | 61.850 | 55.880 | 69.820 | 30.740 | 175.020 | 176.510 | 120.410 | 110.950 |
| **227G** | 8.450 | 110.950 | 45.230 | 61.850 | 999.000 | 69.820 | 174.130 | 175.020 | 114.670 | 119.620 |
| **228M** | 8.260 | 119.620 | 55.330 | 45.230 | 32.800 | 999.000 | 176.680 | 174.130 | 110.950 | 109.890 |
| **229G** | 8.470 | 109.890 | 44.910 | 55.330 | 999.000 | 32.800 | 173.790 | 176.680 | 119.620 | 115.820 |
| **230T** | 8.090 | 115.820 | 59.570 | 44.910 | 69.470 | 999.000 | 173.100 | 173.790 | 109.890 | 139.350 |
| **231P** | 999.000 | 139.350 | 63.560 | 59.570 | 31.720 | 69.470 | 177.560 | 173.100 | 115.820 | 109.700 |
| **232G** | 8.530 | 109.700 | 45.230 | 63.560 | 999.000 | 31.720 | 174.350 | 177.560 | 139.350 | 115.370 |
| **233S** | 8.130 | 115.370 | 58.360 | 45.230 | 63.720 | 999.000 | 174.560 | 174.350 | 109.700 | 121.680 |
| **234Q** | 8.440 | 121.680 | 55.760 | 58.360 | 29.260 | 63.720 | 175.700 | 174.560 | 115.370 | 121.380 |
| **235M** | 8.290 | 121.380 | 55.080 | 55.760 | 32.890 | 29.260 | 175.420 | 175.700 | 121.680 | 126.710 |
| **236A** | 8.280 | 126.710 | 55.010 | 55.080 | 18.010 | 32.890 | 175.270 | 175.420 | 121.380 | 135.470 |
| **237P** | 999.000 | 135.470 | 62.830 | 55.010 | 31.880 | 18.010 | 176.900 | 175.270 | 126.710 | 120.180 |
| **238V** | 8.200 | 120.180 | 62.090 | 62.830 | 32.670 | 31.880 | 176.170 | 176.900 | 135.470 | 119.190 |
| **239S** | 8.330 | 119.190 | 57.800 | 62.090 | 63.660 | 32.670 | 174.240 | 176.170 | 120.180 | 124.390 |
| **240L** | 8.310 | 124.390 | 54.920 | 57.800 | 42.330 | 63.660 | 176.840 | 174.240 | 119.190 | 119.550 |
| **241N** | 8.420 | 119.550 | 53.080 | 54.920 | 38.580 | 42.330 | 174.490 | 176.840 | 124.390 | 121.520 |
| **242V** | 7.930 | 121.520 | 59.590 | 53.080 | 32.470 | 38.580 | 174.110 | 174.490 | 119.550 | 139.100 |
| **243P** | 999.000 | 139.100 | 62.910 | 59.590 | 31.940 | 32.470 | 176.460 | 174.110 | 121.520 | 122.230 |
| **244R** | 8.380 | 122.230 | 53.710 | 62.910 | 30.150 | 31.940 | 174.450 | 176.460 | 139.100 | 136.560 |
| **245P** | 999.000 | 136.560 | 63.150 | 53.710 | 31.960 | 30.150 | 176.640 | 174.450 | 122.230 | 118.140 |
| **246N** | 8.510 | 118.140 | 53.300 | 63.150 | 38.390 | 31.960 | 174.990 | 176.640 | 136.560 | 120.500 |
| **247Q** | 8.260 | 120.500 | 55.690 | 53.300 | 29.450 | 38.390 | 175.710 | 174.990 | 118.140 | 121.350 |
| **248V** | 8.220 | 121.350 | 62.080 | 55.690 | 32.600 | 29.450 | 176.010 | 175.710 | 120.500 | 119.500 |
| **249S** | 8.390 | 119.500 | 58.020 | 62.080 | 63.930 | 32.600 | 174.330 | 176.010 | 121.350 | 110.880 |
| **250G** | 8.200 | 110.880 | 44.500 | 58.020 | 999.000 | 63.930 | 171.350 | 174.330 | 119.500 | 134.260 |
| **251P** | 999.000 | 134.260 | 62.810 | 44.500 | 32.040 | 999.000 | 176.790 | 171.350 | 110.880 | 120.800 |
| **252V** | 8.250 | 120.800 | 62.200 | 62.810 | 32.510 | 32.040 | 176.070 | 176.790 | 134.260 | 126.010 |
| **253M** | 8.450 | 126.010 | 52.720 | 62.200 | 32.290 | 32.510 | 174.090 | 176.070 | 120.800 | 136.640 |
| **254P** | 999.000 | 136.640 | 62.910 | 52.720 | 32.000 | 32.290 | 176.610 | 174.090 | 126.010 | 115.840 |
| **255S** | 8.340 | 115.840 | 58.110 | 62.910 | 63.550 | 32.000 | 174.080 | 176.610 | 136.640 | 123.120 |
| **256M** | 8.270 | 123.120 | 52.910 | 58.110 | 32.630 | 63.550 | 173.560 | 174.080 | 115.840 | 138.740 |
| **257P** | 999.000 | 138.740 | 999.000 | 52.910 | 999.000 | 32.630 | 174.710 | 173.560 | 123.120 | 135.160 |
| **258P** | 999.000 | 135.160 | 63.150 | 999.000 | 31.790 | 999.000 | 177.560 | 174.710 | 138.740 | 108.720 |
| **259G** | 8.360 | 108.720 | 45.130 | 63.150 | 999.000 | 31.790 | 174.090 | 177.560 | 135.160 | 119.610 |
| **260Q** | 8.080 | 119.610 | 55.790 | 45.130 | 29.190 | 999.000 | 175.610 | 174.090 | 108.720 | 121.550 |
| **261W** | 8.100 | 121.550 | 57.090 | 55.790 | 29.250 | 29.190 | 175.930 | 175.610 | 119.610 | 121.850 |
| **262Q** | 8.050 | 121.850 | 55.610 | 57.090 | 29.460 | 29.250 | 175.140 | 175.930 | 121.550 | 121.280 |

| Residue (CBP-TAZ4) | H (ppm) | N (ppm) | CA (ppm) | CA(i-1) (ppm) | CB (ppm) | CB(i-1) (ppm) | C (ppm) | C(i-1) (ppm) | N(i-1) (ppm) | N(i+1) (ppm) |
|---|---|---|---|---|---|---|---|---|---|---|
| **263Q** | 8.150 | 121.280 | 55.380 | 55.610 | 29.430 | 29.460 | 175.080 | 175.140 | 121.850 | 127.080 |
| **264A** | 8.270 | 127.080 | 50.240 | 55.380 | 18.070 | 29.430 | 175.220 | 175.080 | 121.280 | 135.040 |
| **265P** | 999.000 | 135.040 | 62.620 | 50.240 | 31.790 | 18.070 | 176.570 | 175.220 | 127.080 | 123.580 |
| **266L** | 8.280 | 123.580 | 52.930 | 62.620 | 41.410 | 31.790 | 175.490 | 176.570 | 135.040 | 136.090 |
| **267P** | 999.000 | 136.090 | 63.000 | 52.930 | 31.790 | 41.410 | 176.840 | 175.490 | 123.580 | 120.020 |
| **268Q** | 8.440 | 120.020 | 55.930 | 63.000 | 29.250 | 31.790 | 175.930 | 176.840 | 136.090 | 121.610 |
| **269Q** | 8.390 | 121.610 | 55.480 | 55.930 | 29.380 | 29.250 | 175.570 | 175.930 | 120.020 | 122.960 |
| **270Q** | 8.430 | 122.960 | 53.510 | 55.480 | 28.770 | 29.380 | 173.750 | 175.570 | 121.610 | 136.760 |
| **271P** | 999.000 | 136.760 | 62.840 | 53.510 | 31.850 | 28.770 | 176.520 | 173.750 | 122.960 | 122.340 |
| **272M** | 8.350 | 122.340 | 53.700 | 62.840 | 29.950 | 31.850 | 174.250 | 176.520 | 136.760 | 136.850 |
| **273P** | 999.000 | 136.850 | 63.240 | 53.700 | 31.920 | 29.950 | 177.280 | 174.250 | 122.340 | 109.000 |
| **274G** | 8.410 | 109.000 | 44.860 | 63.240 | 999.000 | 31.920 | 173.700 | 177.280 | 136.850 | 122.840 |
| **275L** | 7.980 | 122.840 | 52.870 | 44.860 | 41.650 | 999.000 | 175.200 | 173.700 | 109.000 | 136.280 |
| **276P** | 999.000 | 136.280 | 62.860 | 52.870 | 31.900 | 41.650 | 176.520 | 175.200 | 122.840 | 121.920 |
| **277R** | 8.420 | 121.920 | 53.300 | 62.860 | 29.250 | 31.900 | 174.230 | 176.520 | 136.280 | 136.910 |
| **278P** | 999.000 | 136.910 | 62.910 | 53.300 | 32.050 | 29.250 | 176.630 | 174.230 | 121.920 | 121.150 |
| **279V** | 8.270 | 121.150 | 62.460 | 62.910 | 32.440 | 32.050 | 176.270 | 176.630 | 136.910 | 124.960 |
| **280I** | 8.220 | 124.960 | 60.650 | 62.460 | 38.450 | 32.440 | 176.020 | 176.270 | 121.150 | 120.030 |
| **281S** | 8.360 | 120.030 | 57.880 | 60.650 | 63.690 | 38.450 | 174.550 | 176.020 | 124.960 | 122.890 |
| **282M** | 8.470 | 122.890 | 55.590 | 57.880 | 32.450 | 63.690 | 176.360 | 174.550 | 120.030 | 121.170 |
| **283Q** | 8.340 | 121.170 | 55.930 | 55.590 | 29.140 | 32.450 | 175.860 | 176.360 | 122.890 | 125.220 |
| **284A** | 8.300 | 125.220 | 52.640 | 55.930 | 18.880 | 29.140 | 177.820 | 175.860 | 121.170 | 119.420 |
| **285Q** | 8.290 | 119.420 | 55.720 | 52.640 | 29.340 | 18.880 | 175.790 | 177.820 | 125.220 | 125.000 |
| **286A** | 8.230 | 125.000 | 52.300 | 55.720 | 18.970 | 29.340 | 177.400 | 175.790 | 119.420 | 123.150 |
| **287A** | 8.190 | 123.150 | 52.320 | 52.300 | 18.950 | 18.970 | 177.680 | 177.400 | 125.000 | 119.040 |
| **288V** | 8.000 | 119.040 | 61.950 | 52.320 | 32.760 | 18.950 | 175.730 | 177.680 | 123.150 | 127.760 |
| **289A** | 8.330 | 127.760 | 52.170 | 61.950 | 19.380 | 32.760 | 177.630 | 175.730 | 119.040 | 108.510 |
| **290G** | 8.150 | 108.510 | 44.380 | 52.170 | 999.000 | 19.380 | 171.600 | 177.630 | 127.760 | 134.270 |
| **291P** | 999.000 | 134.270 | 62.880 | 44.380 | 32.030 | 999.000 | 176.890 | 171.600 | 108.510 | 121.170 |
| **292R** | 8.430 | 121.170 | 55.720 | 62.880 | 30.550 | 32.030 | 176.130 | 176.890 | 134.270 | 123.220 |
| **293M** | 8.400 | 123.220 | 52.930 | 55.720 | 32.160 | 30.550 | 174.170 | 176.130 | 121.170 | 136.850 |
| **294P** | 999.000 | 136.850 | 63.080 | 52.930 | 32.050 | 32.160 | 176.710 | 174.170 | 123.220 | 116.220 |
| **295S** | 8.390 | 116.220 | 58.030 | 63.080 | 63.720 | 32.050 | 174.770 | 176.710 | 136.850 | 121.490 |
| **296V** | 8.190 | 121.490 | 62.580 | 58.030 | 32.370 | 63.720 | 176.360 | 174.770 | 116.220 | 123.170 |
| **297Q** | 8.370 | 123.170 | 56.150 | 62.580 | 29.130 | 32.370 | 176.130 | 176.360 | 121.490 | 999.000 |

**Supplementary Table 7** NMR experiments and acquisition parameters used for the backbone assignment of NUPR1. All experiments were performed on a 950 MHz (22.3 T) Bruker NMR spectrometer equipped with a 5 mm TCI probe (Cryogenic HNC inverse probe with cooled $^{1}H$, $^{15}N$, $^{13}C$ preamplifiers, $^{1}H$ inner coil and $^{13}C/^{15}N$ outer coil). Indirect dimension quadrature detection for all experiments was performed using States-TPPI. All data were processed using NMRPipe[7] and NUS reconstruction was performed using SMILE.[8] The assignment was performed on the following sample: 118 μM $^{13}C$, $^{15}N$ labelled NUPR1, 25 mM Tris, 150 mM NaCl, 1 mM TCEP, 0.05% $NaN_3$, pH 7.0, 5% $D_2O$ with the temperature set to 288K.

| Experiment name | Pulse sequence | D1 (s) | NS | DS | NUS sampled percentage (%) | Total points (real+ imaginary) (F1,F2,F3) | Offsets (F1,F2,F3 - ppm) | SW (F1, F2, F3 - ppm) | AQ (F1, F2, F3 - ms) | Nucleus (F1, F2, F3) |
|---|---|---|---|---|---|---|---|---|---|---|
| **HSQC** | hsqcfpf3gpphwg[9–12] | 1 | 4 | 16 | 100 (no NUS) | 2048, 512 | 4.69, 118.5 | 11.95, 24 | 90, 111 | H, N |
| **HNCO** | hncogpwg3d[13–15] | 1 | 4 | 32 | 16 | 2048, 84, 340 | 4.7, 118.5, 172.5 | 13.84, 24, 20 | 77.82, 18.17, 35.56 | H, N, C |
| **HN(CA)CO** | hncacogpwg3d[15,16] | 0.9 | 8 | 32 | 15 | 2048, 170, 170 | 4.7, 118.5, 172.5 | 13.84, 24, 7 | 77.82, 36.77, 50.80 | H, N, C |
| **HNCACB** | hncacbgpwg3d[17,18] | 1 | 8 | 32 | 30 | 2048, 170, 340 | 4.7, 118.5, 43 | 13.84, 24, 80 | 77.82, 36.77, 8.89 | H, N, C |
| **CBCA(CO)NH** | cbcaconhgpwg3d[18,19] | 1 | 16 | 32 | 25 | 2048, 112, 256 | 4.7, 118.5, 43 | 13.84, 24, 80 | 77.82, 24.22, 6.69 | H, N, C |
| **(H)N(COCA)NH** | hncocannhgpwg3d[20–22] | 1 | 16 | 64 | 25 | 2048, 160, 256 | 4.69, 118.5, 118.5 | 13.84, 24, 24 | 77.82, 34.61, 55.37 | H, N, N |
| **CON** | c_con_iasq[23,24] | 1 | 40 | 64 | 100 (no NUS) | 1024, 256 | 173.5, 124 | 12.09, 38.00 | 177.2, 35.0 | C, N |

**Supplementary Table 8** NMR experiments and acquisition parameters used for the backbone assignment of JPT2. The COCON and CBCACON experiments were performed on a 600 MHz (14.1 T) Bruker NMR Spectrometer equipped with an Avance III HD console and a 5 mm TXO cryoprobe (Cryogenic CNH probe with cooled $^{1}H$, $^{15}N$, $^{13}C$ preamplifiers, $^{13}C/^{15}N$ inner coil optimised to $^{13}C$, $^{1}H$ outer coil). All other experiments were performed on a 950 MHz (22.3 T) Bruker NMR spectrometer equipped with an Avance NEO console and a 5 mm TCI probe (Cryogenic HNC inverse probe with cooled $^{1}H$, $^{15}N$, $^{13}C$ preamplifiers, $^{1}H$ inner coil and $^{13}C/^{15}N$ outer coil). Indirect dimension quadrature detection for all experiments was performed using States-TPPI. All data were processed using NMRPipe[7] and NUS reconstruction was performed using SMILE.[8] The assignment was performed on the following sample: 205 µM $^{13}C$, $^{15}N$ labelled JPT2, 20 mM HEPES, 110 mM KCl, 10 mM NaCl, 1 mM TCEP, 0.05% $NaN_3$, pH 7.2, 5% $D_2O$ with the temperature set to 278K.

| Experiment name | Pulse sequence | D1 (s) | NS | DS | NUS sampled percentage (%) | Total points (real+imaginary) (F1,F2,F3) | Offsets (F1,F2,F3 - ppm) | SW (F1, F2, F3 - ppm) | AQ (F1, F2, F3 - ms) | Nucleus (F1, F2, F3) |
|---|---|---|---|---|---|---|---|---|---|---|
| **HNCO** | hncogpwg3d[13–15] | 0.7 | 8 | 16 | 20 | 2048,108,160 | 4.692, 118.5, 173.5 | 11.956, 24, 12 | 90.1, 23.3, 27.9 | H, N, C |
| **HSQC** | hsqcfpf3gpphwg[9–12] | 1 | 4 | 16 | 100 (no NUS) | 2048, 512 | 4.692, 118.5 | 11.956, 24 | 90.1, 110.7 | H, N |
| **CON** | c_con_iasq[23,24] | 1 | 40 | 40 | 100 (no NUS) | 1024, 256 | 173.5, 124 | 12.09, 38.0 | 177.2, 35.0 | C, N |
| **(H)N(COCA)NH** | hncocannhgpwg3d[20–22] | 1 | 16 | 16 | 32.3 | 2048, 160, 256 | 4.692, 118.5, 118.5 | 13.8438, 24, 24 | 77.8, 34.6, 55.4 | H, N, N |
| **CBCANCO** | c_hcbcanco_ia3d[25,26] | 0.7 | 8 | 32 | 100 (no NUS) | 1024, 82, 128 | 173.5, 124, 43 | 12.0911, 38, 70 | 177.2, 11.2, 3.8 | C, N, C |
| **HNCACB** | hncacbgpwg3d_sct[17,18] | 0.8 | 8 | 32 | 29.1 | 2048, 170, 340 | 4.69, 118.5, 43 | 11.956, 24, 80 | 90.1, 36.8, 8.9 | H, N, C |
| **CBCA(CO)NH** | cbcaconhgpwg3d[18,19] | 1 | 4 | 32 | 18.4 | 2048, 112, 256 | 4.69, 118.5, 43 | 11.956, 24, 80 | 90.1, 24.2, 6.7 | H, N, C |
| **HN(CA)CO** | hncacogpwg3d_sct[15,16] | 1 | 8 | 32 | 25.0 | 2048, 170, 170 | 4.69, 118.5, 172.5 | 11.956, 24, 7 | 90.1, 36.8, 50.8 | H, N, C |
| **COCON** | c_hcacocon_ia3d[27–29] | 0.9 | 16 | 32 | 25 | 1024, 128, 160 | 173, 121.5, 173 | 12.101, 37.0, 12 | 280.4, 28.4, 44.2 | C, N, C |
| **CBCACON** | c_hcbcacon_ia3d[23–26] | 0.7 | 16 | 32 | 20 | 1024, 136, 158 | 173.5, 124, 43 | 12.095, 38, 70 | 280.4, 29.4, 7.5 | C, N, C |

**Supplementary Methods**

**NUPR1 expression and purification**

Human NUPR1 was expressed and purified as follows: the plasmid pET-28a(+) harbouring the codon-optimized synthetic sequence coding for NUPR1 (UniProt ID O60356-1) was purchased from Genscript. This plasmid was expressed in the *Escherichia coli* (*E. coli*) strain C41(DE3) grown in M9 media with isotopically enriched $^{15}N$ $NH_4Cl$ (1 g/L) as the sole nitrogen source and $^{13}C$-labelled glucose (3 g/L) as the sole carbon source. Cultures were grown at 37°C and induced at an OD between 0.6 and 0.8 by addition of 0.5 mM isopropyl β-D-thiogalactopyranoside (IPTG) and left shaking for 4 hours. Cells were harvested by centrifugation. Pellets from 1L culture growths were dissolved in 20 mL 8.5 mM $NaH_2PO_4$, 11.5 mM $Na_2HPO_4$, 150 mM NaCl, 1 mM tris(2-carboxyethyl)phosphine (TCEP), 5% glycerol, 10 mM imidazole, pH 7, with the addition of Roche protease inhibitors tablets. Dissolved pellets were boiled in a heating block set to 120°C until sedimentation was observed. Pellets were vortexed and flash frozen in $LN_2$. After melting, small amounts of lysozyme and DNAse were added before centrifugation for 1 h at 40,000 × g at 5°C. The supernatant was filtered through 0.8 μm filter before being loaded onto a Ni-NTA column, washed with the lysis buffer, then matching buffer containing 2M NaCl to remove nucleic acids, and eluted with 500 mM imidazole in the absence of high salt. The His tag was cleaved by a TEV protease (containing its own 6xHis tag) at room temperature for 3-4 h. Imidazole was removed using a PD MidiTrap G-10 column (Cytiva) before passing the sample back over the Ni-NTA column to remove the TEV and cleaved His-tag. The protein was then concentrated and injected onto a Superdex 75 column (GE Healthcare) and subjected to size-exclusion chromatography at 5°C in 25 mM Tris, 150 mM NaCl, 1 mM TCEP, pH 7.0. Fractions containing the protein were purified further using a HiTrap SP ion exchange column (Cytiva) using a gradient from 30 mM to 750 mM NaCl, and the eluted protein was then exchanged into 25 mM Tris, 150 mM NaCl, 1 mM TCEP, pH 7.0. The protein was concentrated using Amicon Ultra Centrifugal filters with a 3 kDa cutoff and stored at −80°C until use.

**JPT2 expression and purification**[30,31]

For expression of recombinant JPT2.3, a pET-28a(+) plasmid harbouring the sequence coding for JPT2.3 (UniProt ID Q9H910-3) was expressed in the *E. coli* strain BL21 (DE3) grown in M9 media with isotopically enriched $^{15}N$ $NH_4Cl$ (1 g/L) as the sole nitrogen source and $^{13}C$-labelled glucose (3 g/L) as the sole carbon source. Cultures were grown at 37°C and induced at an OD of 0.8 by addition of 0.4 mM IPTG and left shaking for 4 hours. Cells were harvested by centrifugation. Pellets from 1L culture growths were dissolved in 30 ml of 50 mM Tris-Base pH 8.0, 300 mM NaCl, 5% glycerol, 5 mM imidazole, supplemented with EDTA-free protease inhibitor (Roche). Small amounts of lysozyme and DNase were added and the cell suspension was incubated on ice for 1 hour before sonication, addition of 1% Triton X-100, and centrifugation at 18000 × g for 1 hour. The supernatant was filtered through 0.8 μm filter before being loaded onto a Ni-NTA column, incubated with agitation for 1 hour, washed with 50 mM Tris-Base pH 8.0, 1 M NaCl, 5% glycerol, then washed with 50 mM Tris-Base pH 8.0, 300 mM NaCl, 5% glycerol, 20 mM imidazole and eluted with 50 mM Tris-Base pH 8.0, 300 mM NaCl, 5% glycerol, 250 mM imidazole. Imidazole was reduced to 20 mM by dilution and the N-terminal 8x His tags were removed using TEV protease (described above). The sample was passed back over a Ni-NTA column, and the untagged proteins were collected in the buffer used for lysis. The protein was further purified by size-exclusion chromatography on a Superdex 75 column (GE Healthcare) at 5°C in 20 mM HEPES, 110 mM KCl, 10 mM NaCl, 1 mM TCEP. The protein was concentrated using Amicon Ultra Centrifugal filters with a 10 kDa cutoff, flash frozen, and stored at −80°C until used.

**Supplementary Information References**
